# Mitochondrial stress in Fabry disease

**DOI:** 10.1101/2025.09.16.676526

**Authors:** L Lavalle, H Kurdi, D Moreno Martinez, V Muczynski, S Heales, DA Hughes

## Abstract

Fabry disease (FD) is clinically heterogeneous. As some GLA variants attain similar levels of residual activity but result in a range of phenotypes, our aim is to understand factors influencing phenotypic variability.

The mitochondrial unfolded protein response (mtUPR) is a stress response mechanism activated by multiple forms of mitochondrial dysfunction including accumulation of misfolded protein. As mitochondrial dysfunction has been reported, we investigated intracellular levels of heat shock protein 60 (Hsp60) by western blotting in 27 FD patients: 11 N215S (7 males) and 16 non-N215S (7 males) vs 4 heathy controls (HC, 3 males and 1 female). Serum Fibroblast Growth Factor-21 (FGF21), and Growth Differentiation Factor-15 (GDF-15) were also measured. Clinical outcomes explored included the Mainz Severity Score Index (MSSI), the Age-Adjusting Severity Scores (AASS), estimated glomerular filtration rate (eGFR) and left ventricular mass indexed to height (LVMI). Globotriaosylsphingosine (lyso-Gb3) data was available for a subset of participants.

Hsp60 showed no significant differences between groups (males FD: 0.29 vs HC: 0.11, females HC: 0.26 vs FD: 0.19 Hsp60/LC), however, differences among FD patients were noted. While some had over 2-fold that of HCs, others had less than half of HCs despite genotype and gender. When analysed in terms of severity scores, the N215S group with higher levels of Hsp60 corresponded with a milder phenotype. LVMI and eGFR also seemed to improve with higher levels of Hsp60 only for this group. In terms of FGF-21 and GDF-15, lower levels showed a trend with higher Hsp60 levels and LVMI in the N215S group.

To conclude, our findings suggest a potential role for mtUPR activation, as evidenced by intracellular Hsp60 levels, in modulating cardiac and renal manifestations in Fabry disease. These preliminary associations highlight the need for longitudinal studies to validate Hsp60 and mitokines as biomarkers of disease progression, aiming to inform personalized approaches that improve outcomes across Fabry disease phenotypes.

## 1.0 Introduction

FD is a highly heterogeneous condition and while some patients experience symptoms during childhood, others remain asymptomatic after middle age (Hughes 2017). The disorder is associated with reduced life expectancy, and the impact of aging on disease expression has been extensively documented (Waldek et al. 2009; Hughes et al. 2010; Ohshima et al. 1999; Kooman, Stenvinkel, and Shiels 2020). Aging is characterized by an increasing accumulation of misfolded proteins and proteins damaged by oxidative stress. Additionally, there is a reduction in the expression of endogenous chaperons (Brehme et al. 2014) and lysosomal dysfunction (Leeman et al. 2018), resulting in a cellular proteome imbalance. This imbalance leads to significative changes in protein levels, causing a subsequent loss of solubility and the accumulation of toxic aggregates (Walther et al. 2015). In mammals, these changes are cell type specific, with nondividing long-lived cells accumulating more damage (Sala, Bott, and Morimoto 2017).

For mutations that result in misfolded proteins, such as Fabry GLA missense mutations (Lenders, Stappers, and Brand 2020; Weidemann et al. 2022), the capacity of the proteostasic network is diminished compared to WT, which is potentially progressive as they get older due to the chronic nature of the proteostatic insult (Wang and Kaufman 2012; Roth et al. 2014; Morley et al. 2002; Morimoto 2011). Therefore, the proteostatic imbalance may be considered a major driver of degenerative disease (Hipp, Kasturi, and Hartl 2019; Jovaisaite, Mouchiroud, and Auwerx 2014; Balch, Roth, and Hutt 2011) and may be particularly important for lysosomal disease due to the role of the autophagic machinery in the clearance of proteasome-resistant misfolded proteins (Fregno and Molinari 2019).

As shown by Roth and collaborators, the continuous presence of misfolded proteins leads to an exacerbation of protein folding deficiency and prompts an integrated stress response (ISR), which is an essential precursor of the mitochondrial UPR (mtUPR) (Roth et al. 2014; Anderson and Haynes 2020). MAMs (mitochondrial-associated membranes) are the apposition (or transient contact) between the ER and the mitochondria, and are essential for signal transmission between these organelles (Vance 2014). The Protein Kinase R-like ER Kinase (PERK), a crucial component of the ER unfolded protein response (erUPR)(Harding, Zhang, and Ron 1999), is particularly abundant at these contact sites (Marciniak et al. 2006), and has been proposed to serve as a potential signal transduction hub connecting both the erUPR and the mtUPR (Kang et al. 2022; Rainbolt, Saunders, and Wiseman 2014).

The mtUPR has been described as a coordinated process between the nucleus and the mitochondrial network that aims to maintain mitochondrial health and proteostasis. This conserved adaptive transcriptional response is triggered by various stressors that activate specific kinases, which then phosphorylate the eIF2α translation initiation factor. In cases of ER perturbations, the specific kinase involved is PERK (Naresh & Haynes, 2019). The main transcriptional regulators of the mtUPR are ATF5, ATF4, and CHOP. Once upregulated, these will promote the expression of chaperones and proteases including as heat shock proteins 10 and 60 (Hsp10, Hsp60)(Anderson and Haynes 2020).

Beyond the cell-autonomous activation of the mitochondrial stress response, another regulatory mechanism involving intercellular communication has been proposed. This suggests that the mtUPR can be activated in distant cells, leading to its activation in distant tissues throughout the organism (Qian Zhang 2023; Durieux, Wolff, and Dillin 2011). The signalling molecules that are thought to mediate this communication have been termed mitokines (Kim et al. 2013). In mammals, these include FGF-21 and GDF-15, which exert an impact on systemic energy metabolism, thereby inducing global metabolic remodelling (Qian Zhang 2023) (Suomalainen and Battersby 2018).

A study found elevated serum GDF-15 in 52 subjects with classic FD undergoing ERT. These levels were more pronounced in those subjects with cardiomyopathy, whether accompanied by nephropathy or not. Additionally, correlations between GDF-15 levels and clinical outcomes including GFR and interventricular septal thickness were observed. Interestingly, patients who initiated ERT after the age of 40 displayed higher GDF-15 levels, potentially suggesting the presence of persistent underlying injuries and supporting the importance of timely initiation of this type of treatment (Gregório et al. 2022).

While mild transient stimulation of these mitokines can have positive effects on health and lifespan, prolonged exposure may have detrimental consequences. It could impede systemic stress responses, leading to resistance to mitokine signalling (analogous insulin resistance). Released during the early stages of mitochondrial dysfunction, FGF-21 and GDF-15 have been shown to reflect pathogenesis and tissue-specific metabolic remodelling. These can aid in medical diagnosis, monitor disease progression, and assess the effectiveness of therapy (Suomalainen 2023). Consequently, these mitokines were measured in our cohort of 35 Fabry patients.

## 2.0 Methods

The study received ethical approval from the Health Research Authority (HRA) and Health and Care Research Wales (HCRW) (REC number: 20/WM/0329). Patients are required to give written consent for study participation. Study documents and ethical approval can be found in Appendix 1.

### 2.1 Clinical data collection and analysis

All participants with FD included in this study were documented to have at least one GLA mutation and had measurement of either plasma or leukocyte enzyme activity available on patients notes, done prior to initiation of treatment. Medical history of all patients were retrospectively assessed and, when available, clinical data prior to initiation of treatment was collected and employed to calculate clinical severity scores, i.e., MSSI and AASS, avoiding any influence of treatment effect on organ manifestations. Data regarding clinical events after treatment commencement were used to cluster individuals with FD into clinical signatures.

Cardiac data included electrocardiogram (ECG) and cardiac magnetic resonance (CMR). Left ventricular mass was calculated using the Devereux formula (Devereux et al. 1986) and LVMI was calculated and adjusted for height. Left ventricular hypertrophy (LVH) was define as an LVMI >/= 78 and 74 g/m^2^ in males and females, respectively (Chuang et al. 2014). Renal function was assessed by estimated glomerular filtration rate (GFR), calculated using the CKD Epidemiology Collaboration equation (CKD-EPI)(Lamb, Levey, and Stevens 2013) and CKD was staged according to the Renal Association, UK. The presence of WMLs was evaluated by cerebral magnetic resonance imaging (MRI). Plasma lyso-Gb3 measurements were available for a subset of patients prior to treatment commencement.

#### Classical vs nonclassical FD

participants were classified as classical or late onset / nonclassical FD based on residual enzyme activity (only for males) and the presence or absence of characteristics Fabry symptoms, i.e., acroparesthesia, angiokeratoma, and / or cornea verticillate. Males were classified as having the classical phenotype if (1) their GLA activity was lower than 5% of the mean reference range and (2) had at least one of the characteristic Fabry symptoms. Alternatively, they were classified as nonclassical / late onset FD. For females, the presence of at least 1 characteristic Fabry symptom was sufficient to be consider as having a classical phenotype. Conversely, they were classified as nonclassical / late onset FD (Smid et al. 2014).

#### Clinical signatures

individuals with FD were subsequently clustered according to the predominant organ system affected, defining these as “clinical signatures”.

#### Fabry cardiomyopathy

was defined as (1) the presence of ECG abnormalities included Sokolow-Lyon voltage criteria for LVH, conduction abnormalities (long QT, short PR, and conduction blocks), repolarization abnormalities (T-wave inversions in at least two contiguous), increase voltages and arrhythmias including multifocal ventricular ectopics, ventricular fibrillation, non-sustained ventricular tachycardia, ventricular tachycardia, (2) LVH documented on image scan, and / or (3) the development of serious cardiac events included atrial fibrillation and other clinically significant arrythmia, implantation of an implantable cardiac defibrillator or pacemaker, and myocardial infarction (Nordin et al. 2019).

#### Fabry nephropathy

Latest expert reviews report characteristic FD features in kidney biopsies of Fabry patients with normal GFR and absence of proteinuria. These features include segmental podocyte foot processes effacement, focal sclerosis, Gb3 depositions, and vascular changes (Ortiz et al. 2008; Schiffmann et al. 2017). A more recent study reported glomerular hyperfiltration as a common feature in young patients with FD and highlighted its contribution to the loss of renal function and onset of nephropathy in type 2 diabetes, sickle cell anaemia, and in autosomal dominant polycystic kidney disease (Riccio et al. 2019). Therefore, in this study we defined Fabry nephropathy as (1) the presence of glomerular hyperfiltration (GFR> 130ml/min/1.73m2), (2) a low GFR: for adults under the age of 60 this was defined as an GFR< 90ml/min/1.73m2. For individuals aged 60 or older this was defined according to their age (Fernandes et al. 2015), (3) proteinuria (>150mg/24hs) or an increased albumin creatinine ratio (> 3mg/mmol), and / or (4) the development of serious renal events including CKD stage 3A (GFR<59ml/min/1.73m2), dialysis, and kidney transplant.

#### Fabry cerebrovascular disease

was define as (1) the presence of WML and (2) history of stroke or TIA.

### 2.2 Serum sample collection

Patients’ blood samples were collected in serum separator tubes (BD) and incubated for 2 hours at 4C. These were then spin down at 2500 rpm for 5 minutes and serum was aspirated into 500 ul aliquots and stored at -80 °C.

### 2.3 Cell cultures

Cells were incubated in a humidified atmosphere with 5% CO2 at 37°C (Haraeus Instruments, Michigan, US).

Fibroblast cell lines GM00302, GM00881, GM00882 (Coriell Institute) were grown in Eagle’s Minimum Essential Medium with Earle’s salts and non-essential amino acids (EMEM; Stemcell Technologies; 36550), supplemented with 15% heat-inactivated FBS (Thermo Fisher; 26140079), and 1% Penicillin-Streptomycin (Sigma; P0781).

Peripheral blood mononuclear cells (PBMC) were isolated from 10 ml of blood by Lymphoprep™ (Stemcell Technologies; 07801) gradient separation. PBMCs were grown in Roswell Park Memorial Institute (RPMI) 1640 medium (Thermo Fisher; A10491-01) with 10% heat inactivated FBS. PBMCs for all FD patients and healthy controls (HC) were cultured for a total of 6 days with 2 media renewals.

### 2.4 Cell lysate preparation

Fibroblasts were harvested with TrypLE™ Express Enzyme (Life Technologies; 12604013) and transferred into centrifuge tubes. PBMCs were transferred into centrifuge tubes and spin down at 1500 rpm for 10 minutes at 4°C. Whole cell lysates were obtained by washing twice with Hanksʹ Balanced Salt solution (HBSS; Sigma; H8264). Then, 100 ul of mammalian cell lysis solution (GE Healthcare; 28-941279) was added, supplemented with 1% protease cocktail inhibitor (Sigma; P8340) and 1% of 100mM phenylmethylsulfonyl fluoride (17.4mg/ml in DMSO; Sigma). Total protein quantification was determined using a bicinchoninic acid protein assay kit (BCA; Sigma; BCA1) and supernatants were stored at -80°C until analysis.

### 2.5 Western blot analysis

Lysates containing 10 ug of total protein were combined with 10 ul 4X Bolt™ LDS Sample Buffer (Invitrogen™; B0007), 4 ul of 10X Bolt™ Sample Reducing Agent (Invitrogen™; B0009), 6 ul distilled H20 and gently mixed. Samples were then incubated at 90°C for 5 minutes and centrifuged at 13000 rpm for 30 seconds. Samples were loaded on a Bolt™ 4-12% Bis-Tris Plus Gels (Invitrogen™; NW04120BOX) and the gel was run at 200 volts for 32 minutes. 10 ul of protein ladder (SeeBlue™ Plus2 Pre-stained Protein Standard; Invitrogen™; LC5925) was employed. Proteins were transferred to a nitrocellulose blotting membrane (GE Healthcare Life Sciences; 10600002) and this was blocked using 10 ml of blocking buffer (5% milk solution in 0.1% Tween 20 PBS (Sigma; 70166)) at room temperature for an hour. Then the membrane was treated with primary antibodies labelled with horseradish peroxidase: anti-Hsp60 monoclonal antibody (2.5 ng/ul; Bio-techne, NBP2-34670H) and anti-sodium potassium ATPase antibody (0.1 ng/ul; Abcam, ab185065) as loading control (LC). Development of the membrane was done by pipetting 2 ml of enhanced chemiluminescent substrate solution (SuperSignal™ West Pico PLUS Chemiluminescent Substrate; Thermo Fisher, 34577) at room temperature. Bands were visualised on a BioRad Molecular Imager (Invitrogen™) and signal quantifications were done in pre-saturating blots.

### 2.6 Enzyme-linked immunosorbent assay (ELISA)

ELISA kits were employed to assess mitokines serum levels according to manufacturer’s instructions, FGF-21 (DF2100; R&D) and GDF-15 (DGD150; R&D).

### 2.7 Statistical analysis

All data was analysed using GraphPad Prism 8.0™ (GraphPad Software Inc., California, USA). Parametric data is presented as the mean and standard deviation, while non-parametric data is represented using the median and interquartile range, with scatter plots visualizing individual data points. N value is defined as the number of repeats for each patient and for each independent cell culture experiment. Difference between two discrete populations were analysed by Mann-Whitney U tests and difference between three or more datasets was analysed by Kruskal-Wallis test with post hoc Dunn’s test. Relationship between two continuous variables was determined using Spearman’s rank correlation. P<0.05 was considered to be significant.

#### 2.7.1 mtUPR predictor analyses

Exploratory linear regression analyses were employed to investigate predictors of mtUPR markers, i.e., intracellular Hsp60, FGF-21, and GDF-15. The full list of candidate predictors included GLA protein, GLA activity, age when the sample was taken (‘age at sampling’), and plasma lyso-Gb3 (only available for males as per missingness in females’ dataset for mtUPR outcomes and for GLA protein, i.e., only 5 out of 13 females with hsp60 or GLA protein data also had plasma lyso-Gb3 data available). For the mitokines predictor analyses, Hsp60 was also included as a candidate predictor.

To investigate the relative contributions of mtUPR markers to clinical outcomes i.e., MISSI, AASS, LVMI, and GFR, multiple linear regression analyses were done. For MSSI and GFR, GLA protein and age at sampling were included as covariates. To account for substrate accumulation effect, GLA activity and plasma lyso-Gb3 data were also added as covariates but in separate models. For LVMI there was missingness, as only data for 11 subjects per biological gender group was available. Similarly, N215S males model building was restricted by the number of subjects as they were a total of 10. Therefore, only those variables that showed significant correlation (|sigma| > 0.4) were included in the full model for these datasets (supplementary fig. 2 and 3). Data distribution was assessed using D’Agostino and Pearson, Kolmogorov-Smirnov tests, and Shapiro-Wiki normality tests.

The model-building approach implemented included a variable selection process with an iterative stepwise selection mixed method whereby irrelevant or redundant variables were removed if they did not significantly improve model fit, based on the variable statistical significance. Starting with a full model (including all independent variables) followed by backward elimination allowed to assess the effect of all candidate variables on the dependent variable. A complete case analysis approach was employed to deal with missing data. Best models were then chosen upon their Akaike information criterion corrected for small sample size (AICc)(Chowdhury and Turin 2020). All analyses were done in GraphPad Prism, version 8.0.

## 3.0 Results

### 3.1 Intracellular levels of heat shock proteins in wild type and FD fibroblast

Two types of fibroblasts cell lines harbouring GLA mutants were studied and compared to WT. The nonsense variant R220X and the missense R301Q displayed differences in total levels of both heat shock proteins. For hsp60, R301Q showed an increase to over 186.6% and R227X a reduction to around 44% when compared to control. Similarly, hsp10 was increased to 126.8% and reduced to 18.2% in R301Q and R227X, respectively (fig. 1). These observations could suggest that GLA missense mutations elicit a different mitochondrial response than those mutations that abolish protein synthesis.

**Figure 1.**
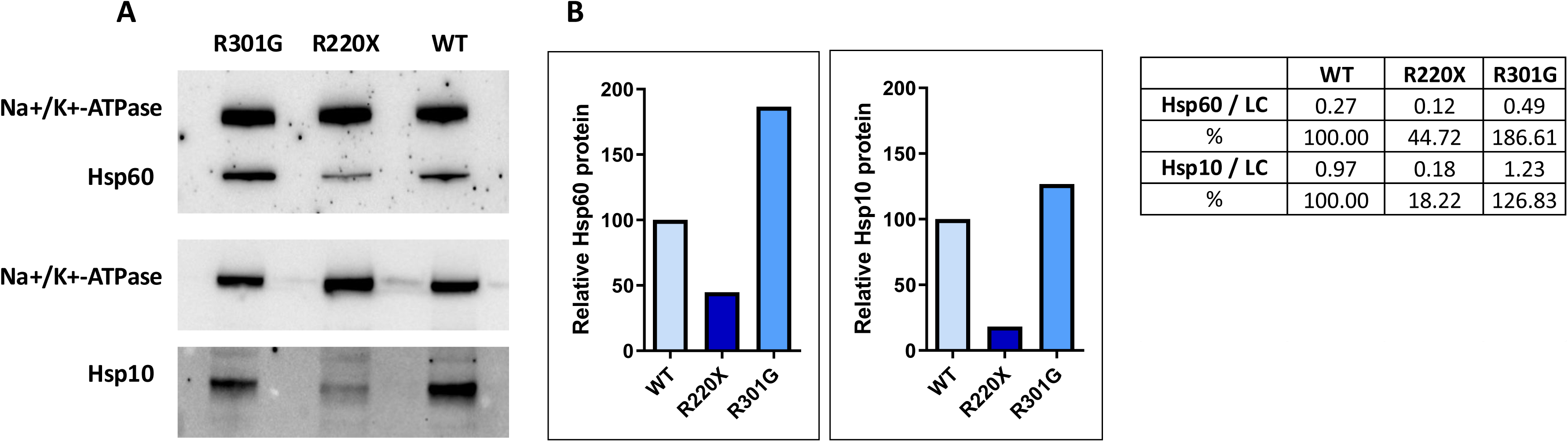
Quantification of heat shock proteins 60 and 10 in normal and FD fibroblasts. Cells lysates containing 30 ug of protein were subject to **(A)** SDS-PAGE and interaction with anti-heat shock protein 60 (Hsp60, 2.5 ng/ul, Bio-techne) antibody and another gel with anti-Hsp10 (1.0 ng/ul, Bio-techne) antibody. Both gels were also incubated with anti-sodium potassium ATPase antibody (Na+/K+-ATPase, 0.1 ng/ul, Abcam) antibody, as a loading control. **(B)** Quantification of target bands (Hsp60 and Hsp10) intensity in each lane was divided by that of Na+/K+-ATPase in the same lane, and the number obtained for wild type (WT) was considered to be 100. n= 1 each.

### 3.2 Intracellular levels of hsp60 in PBMCs from subjects with FD

Summary demographics of the individuals with FD recruited into the study are shown in table 1. Tables 2 and 3 present FD history per participant.

**Table 1.**
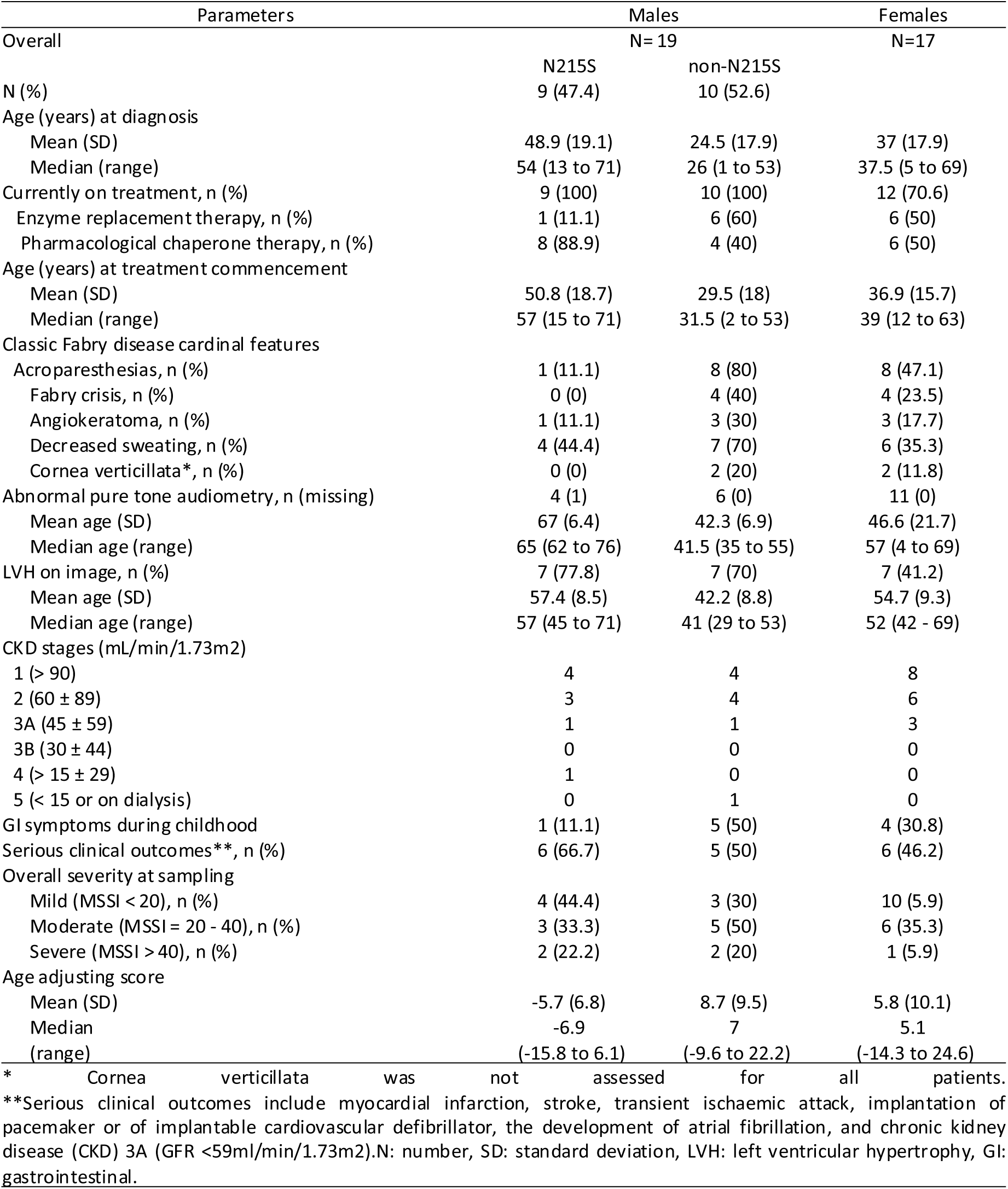
Demographics.

**Table 2:**
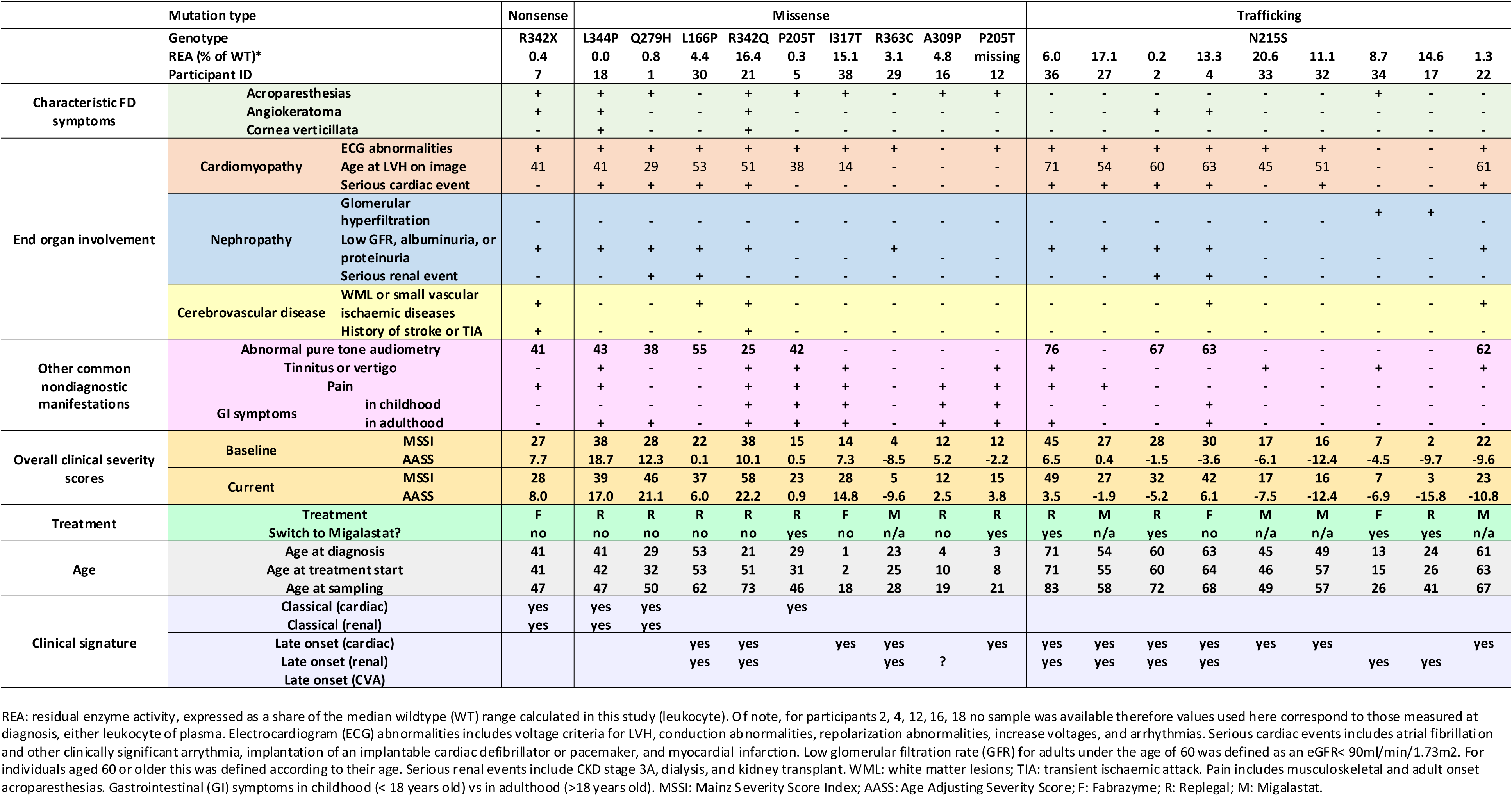
Individual history of Fabry disease (males)

**Table 3:**
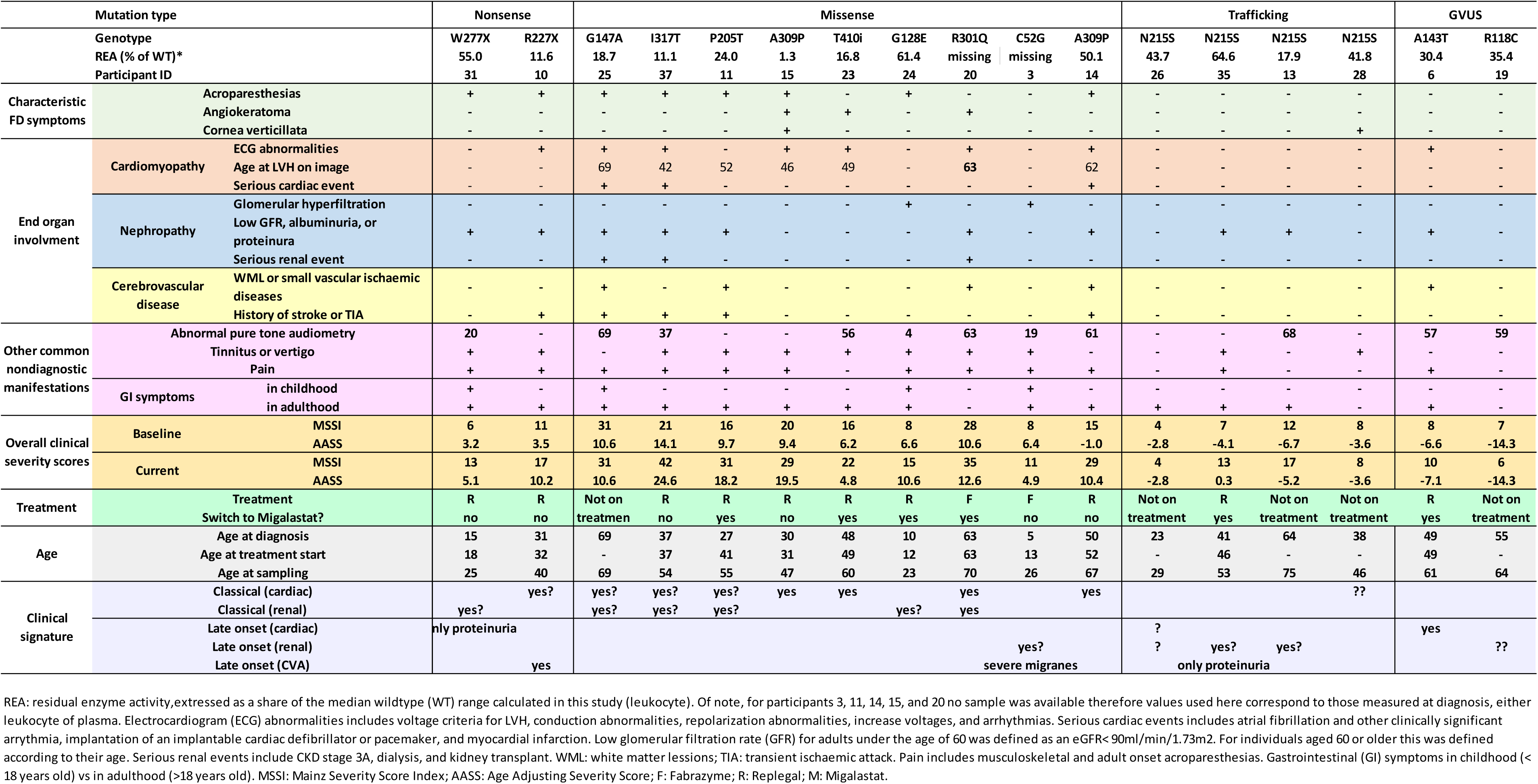
Individual history of Fabry disease (females)

No significant difference between Fabry patients and controls was found in Hsp60 intracellular levels regardless of genotype and gender (males FD: 254.1% vs HC:100%, females FD: 89% vs HC: 100%, p= ns, fig. 2C and supplemental fig. 6), however, differences among FD patients were noted. As Hsp60 levels are reported to increase with age (Cappello et al. 2013), results were analysed against the age at which the sample was taken from each participant. For HC (3 males and a female), Hsp60 levels were higher in older subjects (r2= 1.0 p= 0.04, fig. 3A), however, this was not the case for subjects with FD.

**Fig. 2.**
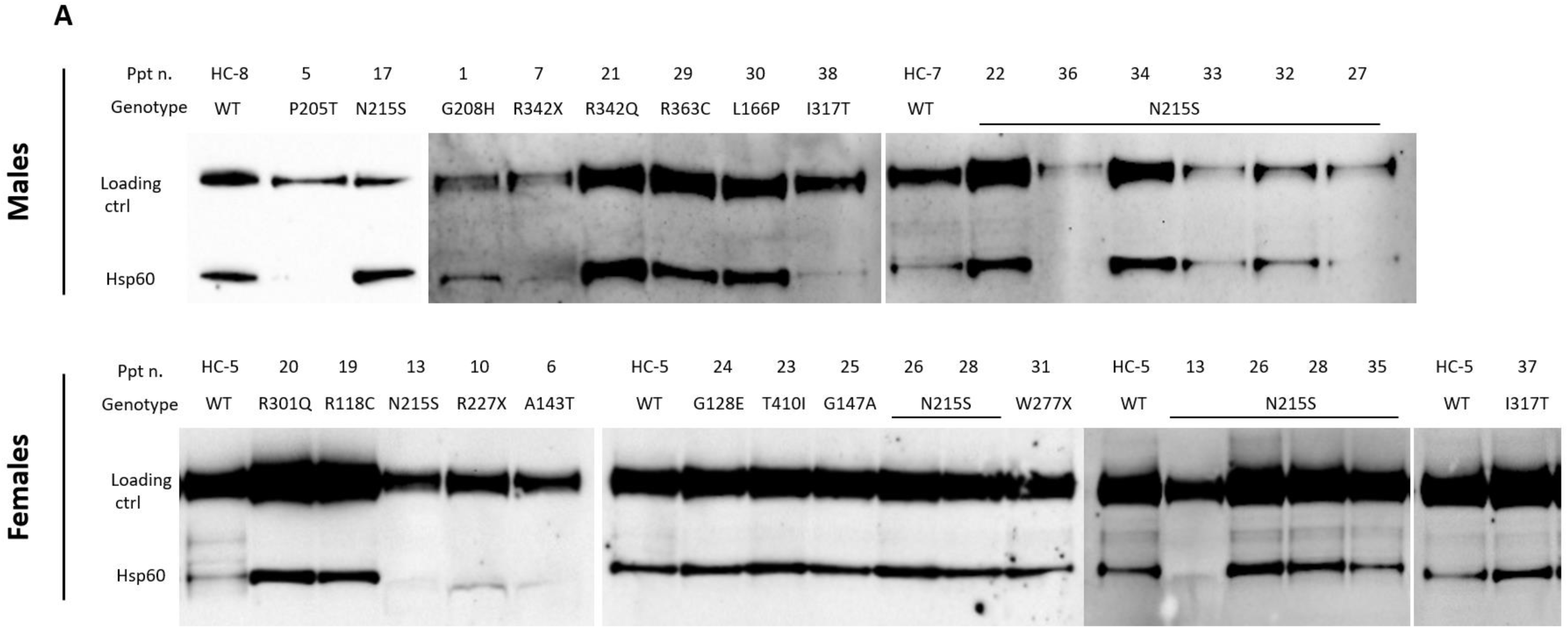
Quantification of Heat shock protein 60 in mononuclear cells from control and participants with Fabry disease. **(A)** Cells lysates containing 10 ug of protein prepared from peripheral blood mononuclear cells (PBMCs) of patients with FD and from a healthy control (HC) were subject to SDS-PAGE and interaction with anti – heat shock protein 60 (Hsp60; 2.5 ng/ul, Bio-techne) antibody and anti-sodium potassium ATPase antibody (Na+/K+-ATPase, 0.1 ng/ul, Abcam) antibody, as a loading control. **(B)** To normalise the results, Hsp60 intensity in each lane was divided by that of Na+/K+-ATPase in the same lane. **(C)** The median of the three male HC was considered to be 100. For females only one HC was available.

**Fig. 2.**
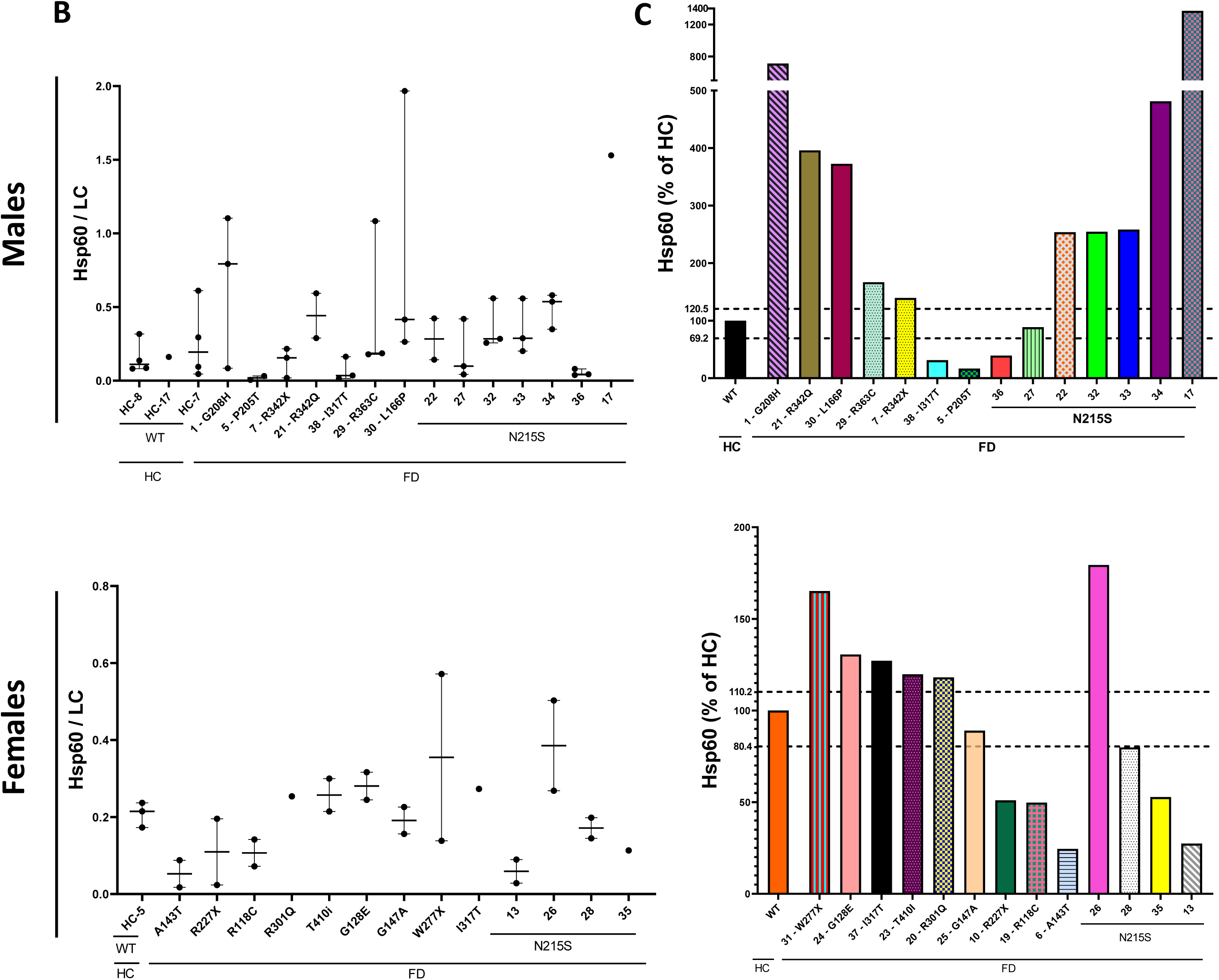
Quantification of Heat shock protein 60 in mononuclear cells from control and participants with Fabry disease. **(A)** Cells lysates containing 10 ug of protein prepared from peripheral blood mononuclear cells (PBMCs) of patients with FD and from a healthy control (HC) were subject to SDS-PAGE and interaction with anti – heat shock protein 60 (Hsp60; 2.5 ng/ul, Bio-techne) antibody and anti-sodium potassium ATPase antibody (Na+/K+-ATPase, 0.1 ng/ul, Abcam) antibody, as a loading control. **(B)** To normalise the results, Hsp60 intensity in each lane was divided by that of Na+/K+-ATPase in the same lane. **(C)** The median of the three male HC was considered to be 100. For females only one HC was available.

**Fig. 3.**
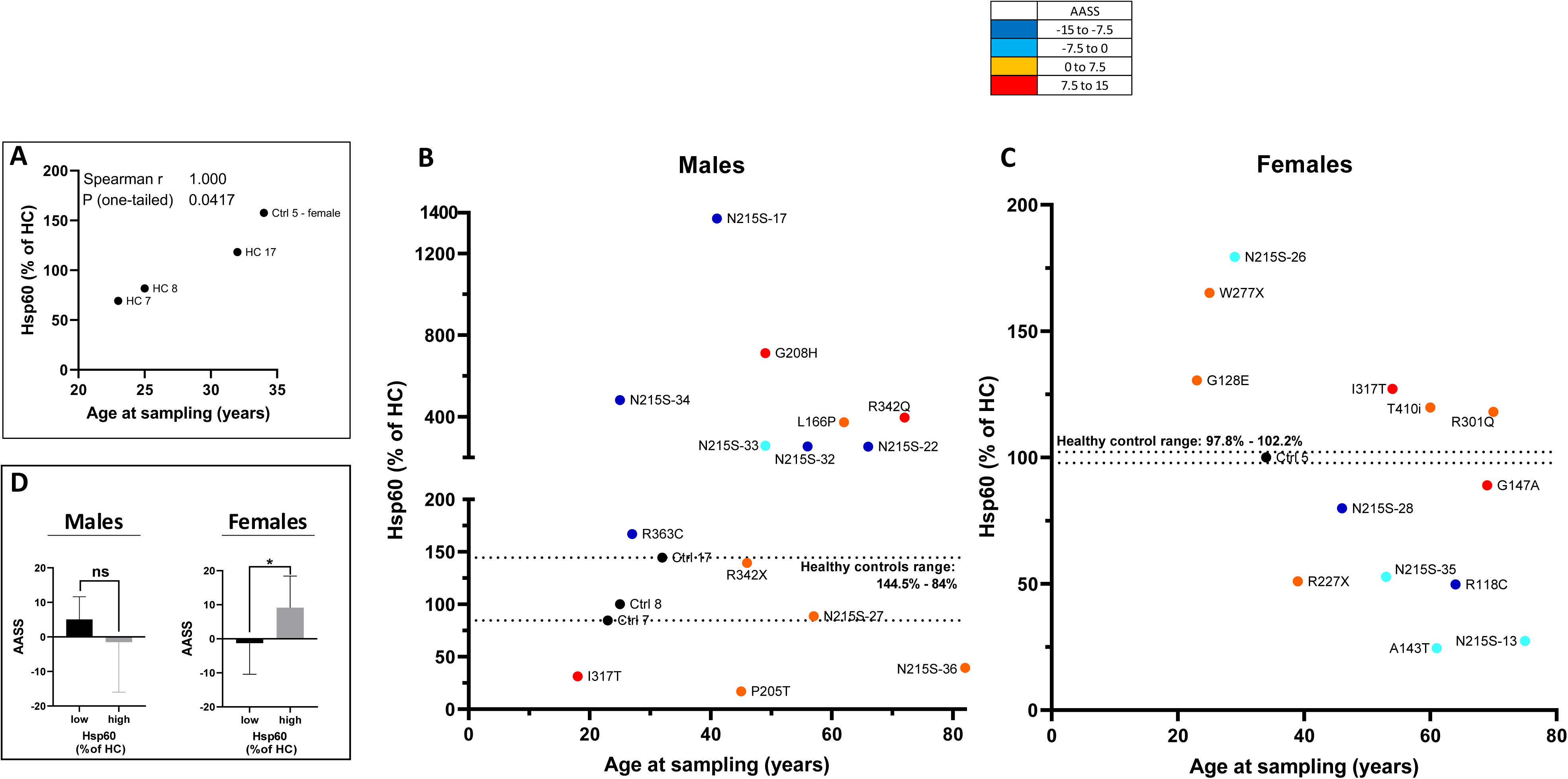
Relation between Heat shock protein 60 and aging. Analysis of relative Heat shock protein 60 (Hsp60) and age for **(A)** control subjects and individuals with Fabry disease (FD): males **(B)** and females **(C)**. Age adjusting Severity Score (AASS) and relative Hsp60 analysis in subjects with FD **(D)**.

some male patients showed Hsp60 levels over four times that of the age and gender-matched HC i.e., participant 34, others displayed the same level or less but at an older age, i.e., patients 27 and 36 (fig. 3B). For females only one HC was available and her Hsp60 levels suggested two patient groups, those with Hsp60 levels above the HC concentration and those under it, with significantly different AASS (-3.6 vs 7.9, p= 0.03, fig. 3D).

Females displayed a positive significant correlation between Hsp60 levels and current AASS (r2= 0.6 p=0.04, fig. 4). Conversely, males did not display differences in AASS when classified upon HC’s Hsp60 level (-7.5 vs 3.5, p= 0.12, fig. 3D), and two weak but opposite trends became apparent between the heat shock protein and the AASS when these were plotted against each other (fig. 4). While N215S males showed a negative trend, suggesting higher levels of Hsp60 corresponded to lower AASS, their non-N215S counterpart displayed a positive trend, hinting that elevated levels of the heat shock protein might be present in subjects with higher severity score. These trends strengthened with the MSSI only for males (N215S: r2= -0.94 p= 0.005, supplementary fig. 4).

**Fig. 4.**
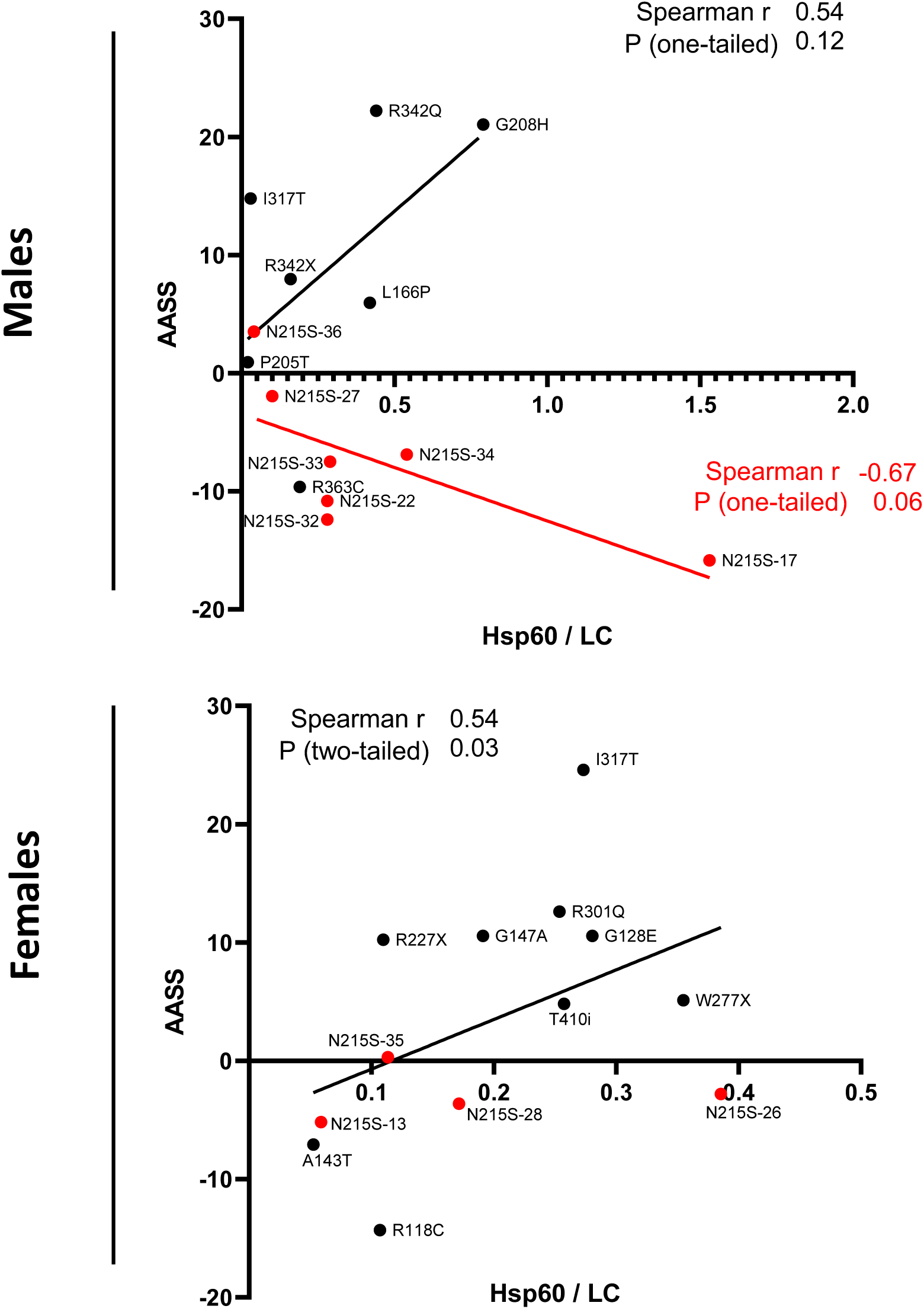
Relation between Heat shock protein 60 (Hsp60) and current clinical severity by the age-adjusting severity score (AASS). Males N215S (n= 7) vs non-N215S (n = 7). Females N215S (n= 4) and non-N215S (n = 9).

In terms of cardiac involvement, the four male patients whose Hsp60 level was lower than controls’ i.e., participants 5, 27, 36, and 38, had LVH. For patients 27 and 36, this was documented before FD diagnosis at 54 and 71 years old, respectively. However, patients 5 and 38 had been on ERT treatment for 7 and 12 years, respectively, before developing myocardial hypertrophy. A correlation analysis between Hsp60 and LVMI showed a negative trend, suggesting lower LVMI magnitudes corresponded with higher levels of Hsp60 in males (r2= -0.82 p= 0.01, fig. 5). The N215S participants 17 and 34 showed the highest levels of Hsp60 within this cohort and have not developed LVH yet. They are currently 41 and 26 years old, respectively. For females, the correlation showed an opposite trend, with higher levels of Hsp60 corresponding with higher LVMI values (r2= 0.66 p= 0.05, fig. 5). In terms of renal involvement, a correlation analysis between Hsp60 and GFR suggested two distinctively opposite trends for male patients. N215S individuals’ GFR was higher with higher levels of Hsp60 while their non-N215S counterpart was lower (N215S r2= 0.89 p= 0.01 vs non-N215S r2=-0.89 p= 0.01, fig. 5). For females, the correlation analysis showed a weak positive trend between Hsp60 levels and GFR (r2= 0.46 p=0.11, fig. 5).

**Fig. 5.**
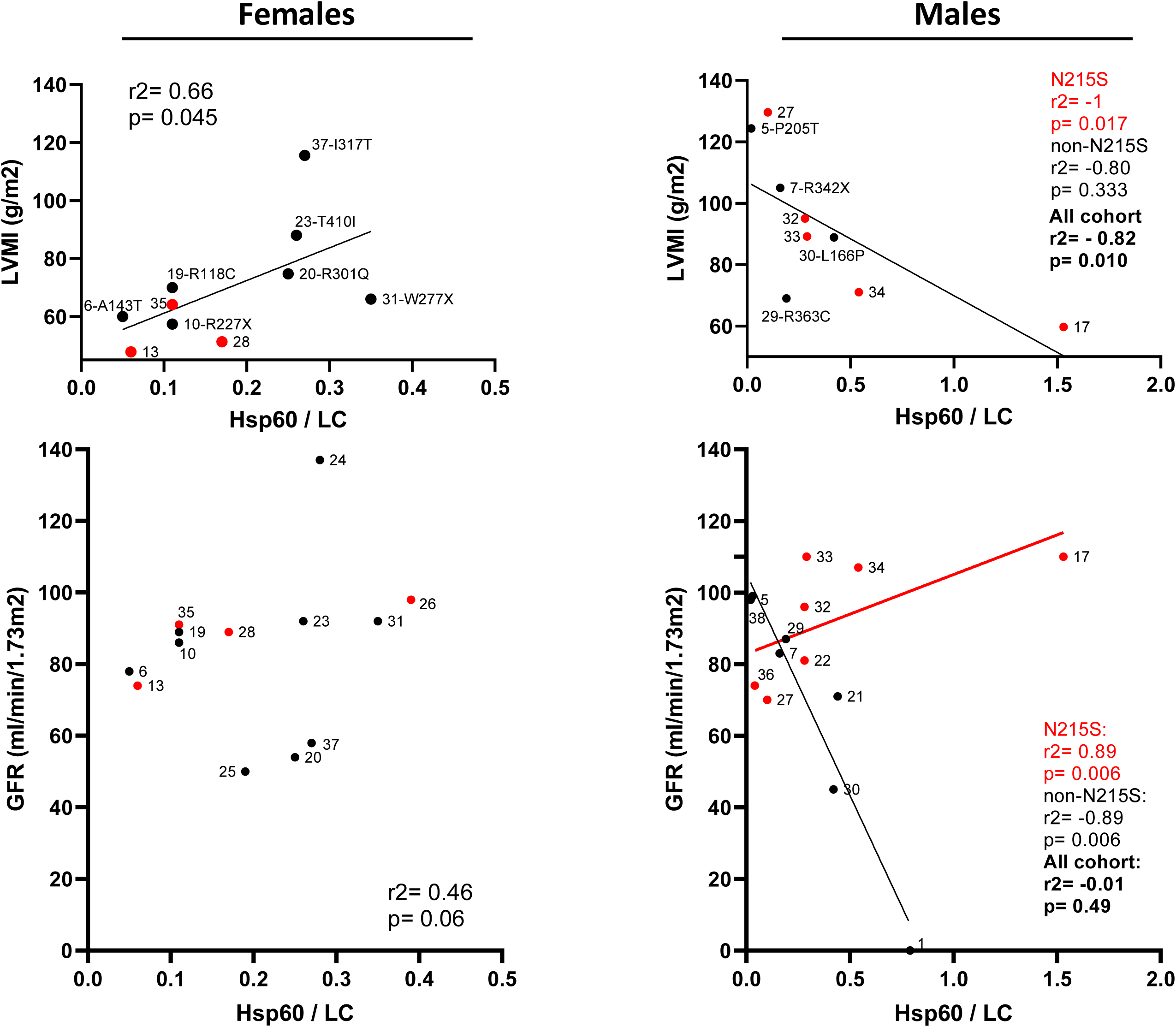
Relationship between Hsp60 and current left ventricular mass index (LVMI) and glomerular filtration rate (GFR) in participants with Fabry disease. LVMI was obtained through cardiac magnetic resonance. Data

To further understand the relation between GLA variants and the mtUPR, exploratory linear regression models were built including intracellular Hsp60 as the dependent variable and GLA protein, GLA activity, age at sampling, and plasma lyso-Gb3 as the independent ones (lyso-Gb3 data was only available for males). While for males the only statistically significant variable was GLA protein (p= 0.002, 95% CI: 0.3 to 0.98, model 5, table A), for females this was age at the time the sample was taken (p= 0.029, 95% CI: -0.01 to -0.0004, table C). The best model for males explained 58.1% of Hsp60 variance (AICc= -30.5, F(1,12)= 16.7, p= 0.002, model 5, table A) and for females it only explained 36% of Hsp60 variance (AICc= -55.8, F(1,11)= 6.2, p= 0.0029, model 3, table C). When looking at the data for males with the N215S genotype (n=7), the best model indicated that GLA protein, activity, and age at sampling were all significant variables for predicting intracellular Hsp60 variance in this group, i.e., model 2, table B, with GLA protein being the most strongly related independent variable.

**Table A.** Multiple linear regression analysis for Hsp60 - All male cohort.

| Model | Independent variables | $\beta$ coefficient | 95% CI | *p*-value | $r^2$ | Adj. $r^2$ | AICc | F | *p*-value |
| --- | --- | --- | --- | --- | --- | --- | --- | --- | --- |
| 1 | Intercept | 0.700 | 0.11 to 1.29 | 0.025 | 0.76 | 0.64 | -16.49 | F(4,8)= 6.2 | 0.014 |
|  | GLA protein | 0.812 | 0.42 to 1.20 | 0.001 |  |  |  |  |  |
|  | GLA activity | -0.040 | -0.08 to 0.002 | 0.060 |  |  |  |  |  |
|  | Plasma lysoGb3 | -0.008 | -0.02 to 0.01 | 0.203 |  |  |  |  |  |
|  | Age | -0.004 | -0.01 to 0.004 | 0.282 |  |  |  |  |  |
| 2 | Intercept | 0.471 | 0.11 to 0.84 | 0.017 | 0.72 | 0.62 | -21.9 | F(3,9)= 7.6 | 0.008 |
|  | GLA protein | 0.790 | 0.40 to 1.18 | 0.001 |  |  |  |  |  |
|  | GLA activity | -0.037 | -0.08 to 0.004 | 0.072 |  |  |  |  |  |
|  | Plasma lysoGb3 | -0.008 | -0.022 to 0.01 | 0.211 |  |  |  |  |  |
| 3 | Intercept | 0.294 | 0.06 to 0.53 | 0.020 | 0.66 | 0.59 | -29.2 | F(2,11)= 10.5 | 0.003 |
|  | GLA protein | 0.775 | 0.40 to 1.15 | 0.001 |  |  |  |  |  |
|  | GLA activity | -0.025 | -0.06 to 0.01 | 0.152 |  |  |  |  |  |
| 4 | Intercept | 0.185 | -0.02 to 0.39 | 0.073 | 0.58 | 0.55 | -30.5 | F(1,12)= 16.7 | 0.002 |
|  | GLA protein | 0.641 | 0.30 to 0.98 | 0.002 |  |  |  |  |  |

**Table B.** Multiple linear regression analysis for Hsp60 - For N215S males.

| Model | Independent variables | $\beta$ coefficient | 95% CI | $p$-value | $r^2$ | Adj. $r^2$ | AICc | F | $p$-value |
| --- | --- | --- | --- | --- | --- | --- | --- | --- | --- |
| 1 | Intercept | 0.949 | 0.79 to 1.11 | 0.002 | 0.99 | 0.99 | n/a | $F(4,2)=1217$ | 0.0008 |
|  | GLA protein | 0.778 | 0.72 to 0.84 | 0.000 |  |  |  |  |  |
|  | GLA activity | -0.029 | -0.04 to -0.02 | 0.004 |  |  |  |  |  |
|  | Plasma lysoGb3 | -0.009 | -0.03 to 0.01 | 0.207 |  |  |  |  |  |
|  | Age | -0.010 | -0.01 to -0.01 | 0.005 |  |  |  |  |  |
| 2 | Intercept | 0.983 | 0.85 to 1.12 | 0.0002 | 0.99 | 0.99 | 11.2 | $F(3,3)=901.9$ | <0.0001 |
|  | GLA protein | 0.774 | 0.72 to 0.83 | <0.0001 |  |  |  |  |  |
|  | GLA activity | -0.030 | -0.04 to -0.02 | 0.001 |  |  |  |  |  |
|  | Age | -0.011 | -0.01 to -0.01 | 0.0003 |  |  |  |  |  |
| 3 | Intercept | 0.651 | 0.05 to 1.26 | 0.041 | 0.94 | 0.91 | -2.1 | $F(2,4)=30.5$ | 0.004 |
|  | GLA protein | 0.711 | 0.40 to 1.02 | 0.003 |  |  |  |  |  |
|  | Age | -0.009 | -0.02 to 0.001 | 0.075 |  |  |  |  |  |
| 4 | Intercept | 0.156 | -0.09 to 0.41 | 0.168 | 0.85 | 0.82 | -9.9 | $F(1,5)=28.5$ | 0.003 |
|  | GLA protein | 0.790 | 0.41 to 1.17 | 0.003 |  |  |  |  |  |

**Table C.** Multiple linear regression analysis for Hsp60 - All female cohort.

| Model | Independent variables | $\beta$ coefficient | 95% CI | *p*-value | $r^2$ | Adj. $r^2$ | AICc | F | *p*-value |
| --- | --- | --- | --- | --- | --- | --- | --- | --- | --- |
| 1 | Intercept | 0.433 | 0.06 to 0.81 | 0.028 | 0.54 | 0.34 | -34.9 | F(3,7)= 2.7 | 0.127 |
|  | GLA protein | 0.047 | -0.19 to 0.28 | 0.644 |  |  |  |  |  |
|  | GLA activity | -0.003 | -0.02 to 0.01 | 0.624 |  |  |  |  |  |
|  | Age | -0.005 | -0.01 to 0.001 | 0.070 |  |  |  |  |  |
| 2 | Intercept | 0.490 | 0.18 to 0.80 | 0.006 | 0.49 | 0.38 | -47.8 | F(2,9)= 4.4 | 0.047 |
|  | GLA activity | -0.002 | -0.01 to 0.01 | 0.521 |  |  |  |  |  |
|  | Age | -0.005 | -0.01 to -0.001 | 0.023 |  |  |  |  |  |
| 3 | Intercept | 0.391 | 0.21 to 0.57 | 0.001 | 0.36 | 0.3 | -55.8 | F(1,11)= 6.2 | 0.029 |
|  | Age | -0.004 | -0.01 to -0.0004 | 0.029 |  |  |  |  |  |
| 4 | Intercept | 0.380 | 0.06 to 0.70 | 0.0260 | 0.39 | 0.26 | -45.8 | F(2,9)= 2.9 | 0.108 |
|  | GLA protein | 0.005 | -0.15 to -0.16 | 0.945 |  |  |  |  |  |
|  | Age | -0.004 | -0.009 to 0.001 | 0.111 |  |  |  |  |  |

### 3.3 Serum mitokines FGF-21 and GDF-15 in FD

Mitokines FGF-21 and GDF-15 were measured in a total of 60 serum samples from Fabry patients (18 males and 17 females) and from apparently healthy age-matched controls (17 males and 8 females, fig. 6C). A comparison of the mitokines serum levels was done between groups and no differences were found except for males GDF-15 values, where Fabry patients had twice the levels of the controls (934.9 vs 559.3 pg/ml, p= 0.002, fig. 6). No differences were found between genotype groups for subjects with FD. The lack of statistical difference between biological sex groups is in agreement with published data (Conte et al. 2021; Conte et al. 2019), therefore males and females were pooled together to study the relation between FGF-21, GDF-15, and aging in our cohort of Fabry patients and controls, as these mitokines are known to increase with age (Conte et al. 2019). While serum levels of FGF-21 and GDF-15 were positively and significantly correlated with age in subjects with FD (FGF-21: r2= 0.39 p= 0.02, GDF-15: r2= 0.64 p< 0.0001, fig. 7), only GDF-15 correlated with age in HCs (r2= 0.63 p= 0.0007, fig. 7).

**Fig. 6.**
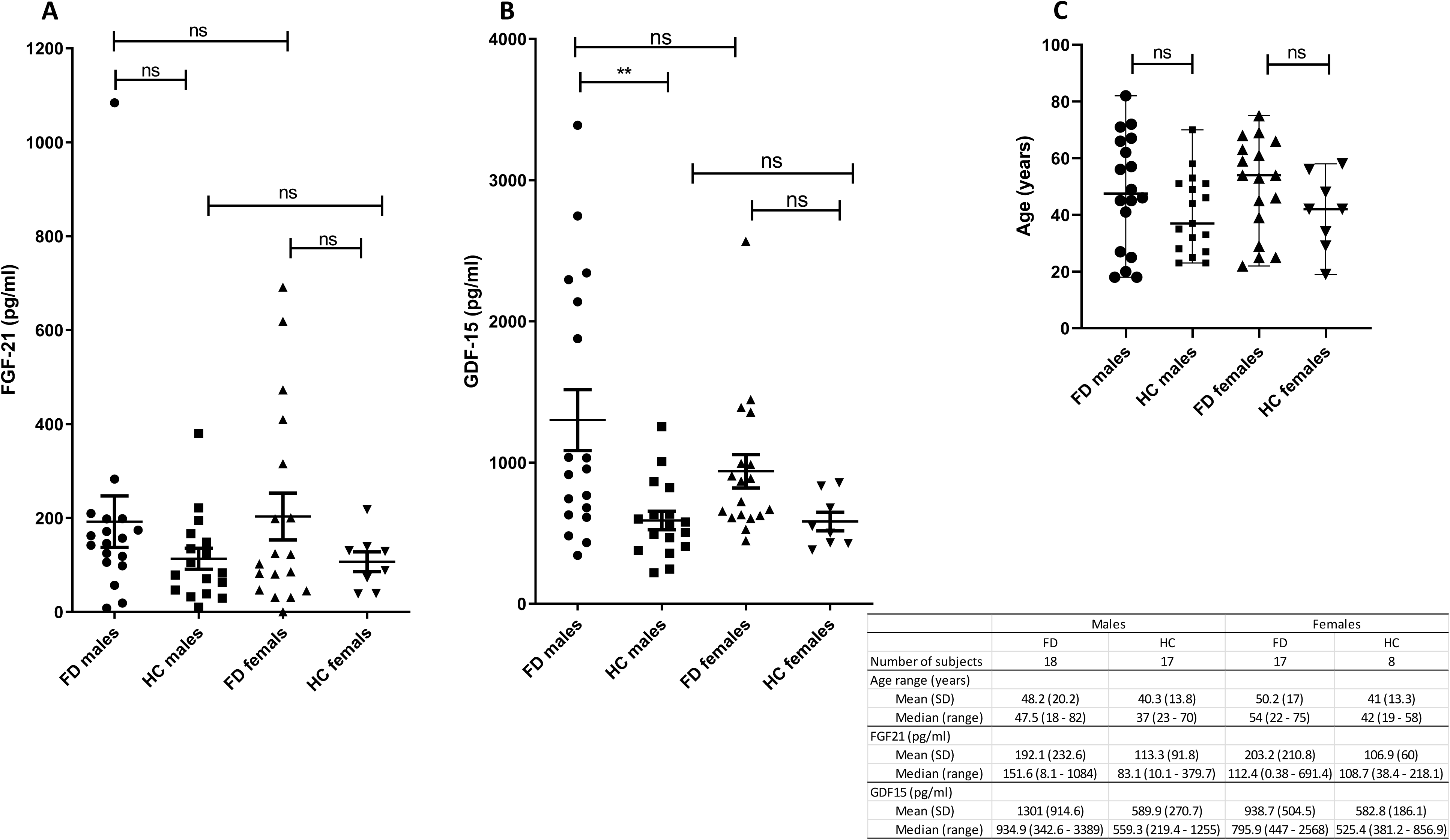
Serum levels of fibroblast growth factor 21 (FGF-21) and growth differentiation factor 15 (GDF-15) in healthy controls and participants with Fabry disease. **(A)** FGF-21 and **(B)** GDF-15 were measured by enzyme-linked immunosorbent assay (ELISA) in patients with Fabry disease (FD, males n= 18, females n= 17) and in healthy controls (HC, males n= 17, females n= 8) age and gender matched **(C)** Median age (years): males: FD 47.5 vs. HC 37, females: FD 54 vs. 42.

**Fig. 7.**
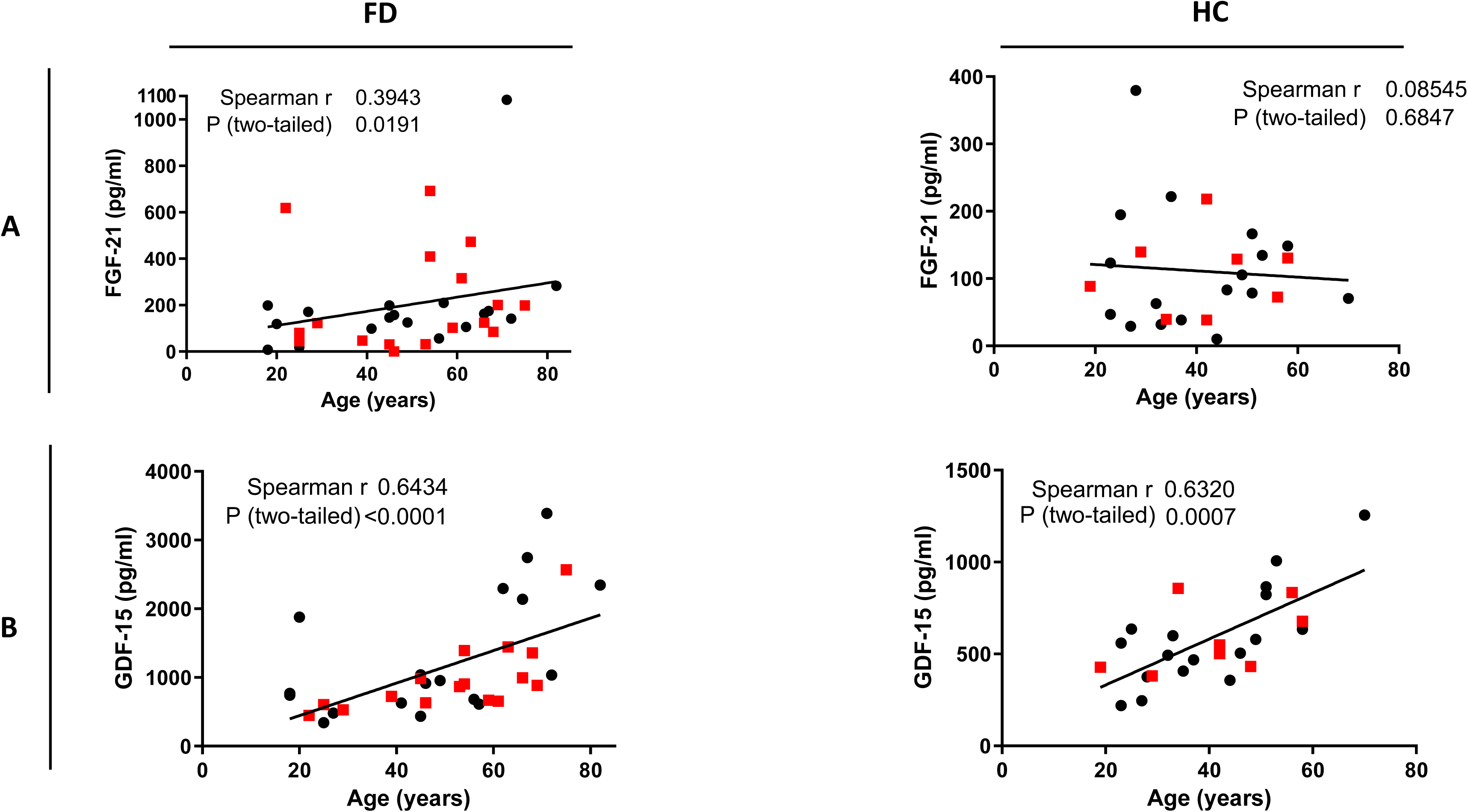
Relation between serum mitokines and aging. **(A)** FGF-21 and **(B)** GDF-15 were measured by enzyme-linked immunosorbent assay (ELISA) in patients with Fabry disease (FD, males n= 18, females n= 17) and in healthy controls (HC, males n= 17, females n= 8). In **red** females and in **black** males.

Correlation analyses between mitokines and intracellular Hsp60 levels showed that for male patients, higher levels of Hsp60 corresponded with lower levels of FGF-21 (r2= -0.75 p= 0.004, fig. 8). Although this was not the case for females, these did show a statistically significant correlation between the heat shock protein and GDF-15 (r2= -0.57 p= 0.05, fig. 8). So, to investigate the association of mitokines serum levels, aging, and Hsp60 in FD, multiple linear regression exploratory modelling was employed. For FGF-21 a statistically significant model was only found for males, though this only explained 46% of the variance in FGF-21 serum levels (AICc= 113.5, F (2,10) = 4.3, p= 0.045, model 4, table D). This model only included 2 independent variables, Hsp60 and GLA protein, but only the former was statistically significant. When looking at males with the N215S genotype, the model that achieved statistical significance included Hsp60, GLA protein, GLA activity, and age at sampling as independent variables (table E, model 2). Statistically significant models were found for both sex groups when testing Hsp60, GLA protein, GLA activity, and age as independent variables on GDF-15 serum levels. For males, regardless of genotype, the only statistically significant independent variable was age of the subject at the time the sample was taken (tables G and H). When looking only at females, age was the only statistically significant variable as well (table I).

**Fig. 8.**
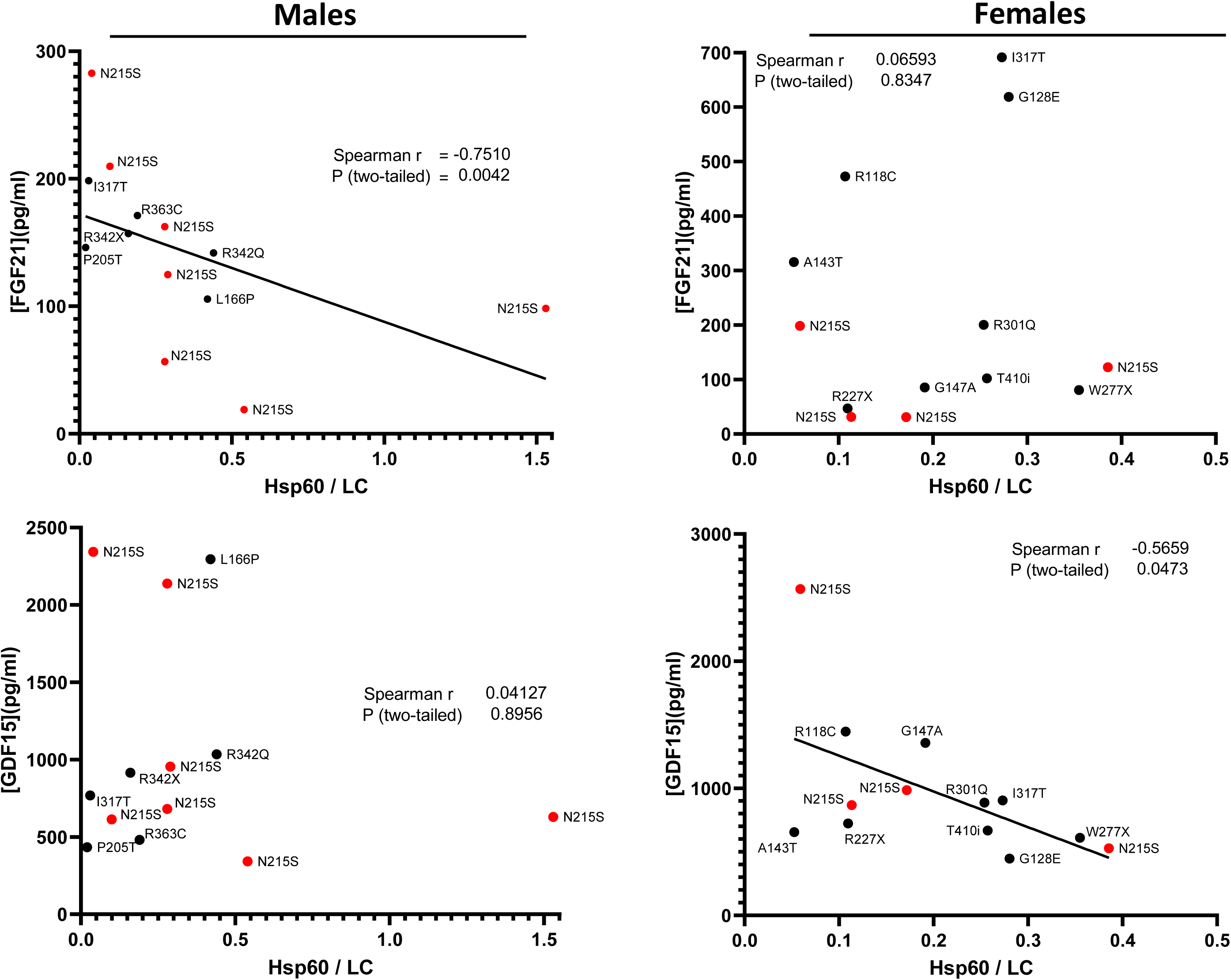
Relation between serum mitokines and Hsp60. FGF-21 and GDF-15

**Table D.** Multiple linear regression analysis for FGF-21 - All male cohort.

| Model | Independent variables | β coefficient | 95% CI | p-value | r<sup>2</sup> | Adj. r<sup>2</sup> | AICc | F | p-value |
| --- | --- | --- | --- | --- | --- | --- | --- | --- | --- |
| 1 | Intercept | 250.800 | 17.15 to 484.5 | 0.039 | 0.61 | 0.28 | 131.9 | F(5,6)=<br>1.9 | 0.235 |
|  | Hsp60 | -314.200 | -615.0 to -13.37 | 0.043 |  |  |  |  |  |
|  | GLA protein | 218.900 | -42.59 to 480.5 | 0.086 |  |  |  |  |  |
|  | GLA activity | -7.188 | -21.30 to 6.922 | 0.259 |  |  |  |  |  |
|  | Plasma lysoGb3 | -2.173 | -6.652 to 2.307 | 0.280 |  |  |  |  |  |
|  | Age | 0.041 | -2.532 to 2.614 | 0.970 |  |  |  |  |  |
| 2 | Intercept | 253.900 | 130.1 to 377.6 | 0.002 | 0.61 | 0.38 | 118.7 | F(4,7)=<br>2.7 | 0.118 |
|  | Hsp60 | -316.500 | -552.9 to -80.15 | 0.016 |  |  |  |  |  |
|  | GLA protein | 221.000 | 15.71 to 426.2 | 0.038 |  |  |  |  |  |
|  | GLA activity | -7.273 | -18.98 to 4.44 | 0.185 |  |  |  |  |  |
|  | Plasma lysoGb3 | -2.195 | -6.01 to 1.62 | 0.216 |  |  |  |  |  |
| 3 | Intercept | 198.300 | 129.8 to 266.7 | 0.0001 | 0.51 | 0.34 | 118 | F(3,9)=<br>3.1 | 0.08 |
|  | Hsp60 | -238.300 | -421.7 to -54.80 | 0.017 |  |  |  |  |  |
|  | GLA protein | 157.000 | -6.76 to 320.8 | 0.058 |  |  |  |  |  |
|  | GLA activity | -3.518 | -12.32 to 5.28 | 0.389 |  |  |  |  |  |
| 4 | Intercept | 178.100 | 132.9 to 223.2 | <0.0001 | 0.46 | 0.35 | 113.5 | F(2,10)=<br>4.3 | 0.045 |
|  | Hsp60 | -214.400 | -383.8 to -45.09 | 0.018 |  |  |  |  |  |
|  | GLA protein | 123.200 | -13.67 to 260.0 | 0.073 |  |  |  |  |  |
| 5 | Intercept | 172.200 | 122.4 to 222.1 | <0.0001 | 0.24 | 0.18 | 113.6 | F(1,11)=<br>3.6 | 0.086 |
|  | Hsp60 | -84.450 | -183.1 to 14.18 | 0.086 |  |  |  |  |  |

**Table E.** Multiple linear regression analysis for FGF-21 - For N215S males.

| Model | Independent variables | β coefficient | 95% CI | p-value | r<sup>2</sup> | Adj. r<sup>2</sup> | AICc | F | p-value |
| --- | --- | --- | --- | --- | --- | --- | --- | --- | --- |
| 1 | Intercept | 1833.000 | -12626 to 16291 | 0.354 | 0.98 | 0.89 | n/a | F(5,1)=<br>10.6 | 0.229 |
|  | Hsp60 | -1985.000 | -17197 to 13226 | 0.345 |  |  |  |  |  |
|  | GLA protein | 1540.000 | -10300 to 13380 | 0.346 |  |  |  |  |  |
|  | GLA activity | -56.990 | -493.5 to 379.5 | 0.345 |  |  |  |  |  |
|  | Plasma lysoGb3 | 7.140 | -173.2 to 187.5 | 0.703 |  |  |  |  |  |
|  | Age | -17.670 | -168.8 to 133.5 | 0.377 |  |  |  |  |  |
| 2 | Intercept | 2276.000 | -171.4 to 4724 | 0.057 | 0.98 | 0.93 | n/a | F(4,2)=<br>21.1 | 0.046 |
|  | Hsp60 | -2463.000 | -4946 to 19.58 | 0.051 |  |  |  |  |  |
|  | GLA protein | 1913.000 | -9.34 to 3836 | 0.051 |  |  |  |  |  |
|  | GLA activity | -70.240 | -145.03 to 4.81 | 0.057 |  |  |  |  |  |
|  | Age | -22.110 | -49.22 to 5.00 | 0.073 |  |  |  |  |  |
| 3 | Intercept | 281.800 | 114.5 to 449.2 | 0.013 | 0.83 | 0.67 | 119.4 | F(3,3)=<br>5.0 | 0.109 |
|  | Hsp60 | -447.100 | -821.5 to -72.71 | 0.032 |  |  |  |  |  |
|  | GLA protein | 354.500 | 17.78 to 691.1 | 0.044 |  |  |  |  |  |
|  | GLA activity | -9.625 | -26.51 to 7.255 | 0.167 |  |  |  |  |  |
| 4 | Intercept | 202.200 | 101.3 to 303.1 | 0.0051 | 0.65 | 0.48 | 82.6 | F(2,4)=<br>3.8 | 0.121 |
|  | Hsp60 | -364.800 | -742.8 to 13.23 | 0.055 |  |  |  |  |  |
|  | GLA protein | 262.900 | -60.92 to 586.8 | 0.087 |  |  |  |  |  |
| 5 | Intercept | 172.000 | 54.97 to 289.0 | 0.0129 | 0.21 | 0.05 | 74.4 | F(1,5)=<br>1.3 | 0.301 |
|  | Hsp60 | -81.720 | -264.0 to 100.5 | 0.301 |  |  |  |  |  |

**Table F.** Multiple linear regression analysis for FGF-21 - All female cohort.

| Model | Independent variables | β coefficient | 95% CI | p-value | r<sup>2</sup> | Adj. r<sup>2</sup> | AICc | F | p-value |
| --- | --- | --- | --- | --- | --- | --- | --- | --- | --- |
| 1 | Intercept | 214.100 | -1209 to 1637 | 0.725 | 0.09 | -0.52 | 147.1 | F(4,6)=<br>0.1 | 0.959 |
|  | Hsp60 | -361.600 | -2731 to 2008 | 0.722 |  |  |  |  |  |
|  | GLA protein | -32.990 | -659.0 to 593.1 | 0.902 |  |  |  |  |  |
|  | GLA activity | 6.093 | -26.60 to 38.78 | 0.664 |  |  |  |  |  |
|  | Age | -1.181 | -18.36 to 16.00 | 0.872 |  |  |  |  |  |
| 2 | Intercept | 218.500 | -1306 to 1743 | 0.749 | 0.01 | -0.36 | 150 | F(3,8)=<br>0.04 | 0.99 |
|  | Hsp60 | 225.500 | -2152 to 2603 | 0.832 |  |  |  |  |  |
|  | GLA activity | -1.671 | -23.21 to 19.87 | 0.863 |  |  |  |  |  |
|  | Age | 0.064 | -18.18 to 18.30 | 0.994 |  |  |  |  |  |
| 3 | Intercept | 223.700 | -200.5 to 647.9 | 0.264 | 0.01 | -0.21 | 143.7 | F(2,9)=<br>0.1 | 0.943 |
|  | Hsp60 | 219.800 | -1405 to 1844 | 0.767 |  |  |  |  |  |
|  | GLA activity | -1.714 | -18.17 to 14.75 | 0.819 |  |  |  |  |  |
| 9 | Intercept | 198.500 | -111.3 to 508.2 | 0.186 | 0.01 | -0.1 | 148.5 | F(1,11)=<br>0.1 | 0.802 |
|  | Hsp60 | 160.100 | -1211 to 1531 | 0.802 |  |  |  |  |  |

**Table G.**
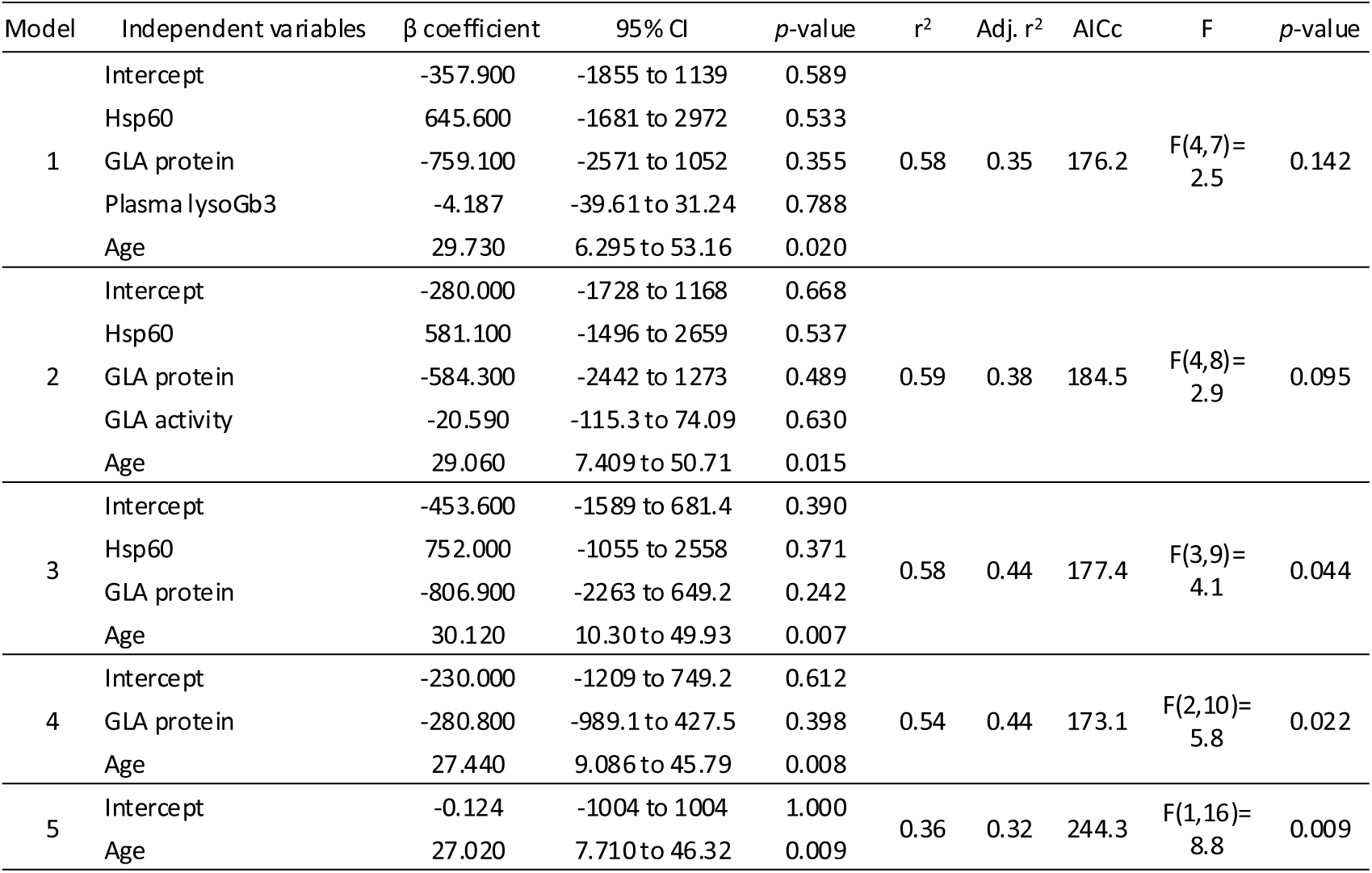
Multiple linear regression analysis for GDF-15 - All male cohort.

| Model | Independent variables | β coefficient | 95% CI | p-value | r<sup>2</sup> | Adj. r<sup>2</sup> | AICc | F | p-value |
| --- | --- | --- | --- | --- | --- | --- | --- | --- | --- |
| 1 | Intercept | -357.900 | -1855 to 1139 | 0.589 | 0.58 | 0.35 | 176.2 | F(4,7)=<br>2.5 | 0.142 |
|  | Hsp60 | 645.600 | -1681 to 2972 | 0.533 |  |  |  |  |  |
|  | GLA protein | -759.100 | -2571 to 1052 | 0.355 |  |  |  |  |  |
|  | Plasma lysoGb3 | -4.187 | -39.61 to 31.24 | 0.788 |  |  |  |  |  |
|  | Age | 29.730 | 6.295 to 53.16 | 0.020 |  |  |  |  |  |
| 2 | Intercept | -280.000 | -1728 to 1168 | 0.668 | 0.59 | 0.38 | 184.5 | F(4,8)=<br>2.9 | 0.095 |
|  | Hsp60 | 581.100 | -1496 to 2659 | 0.537 |  |  |  |  |  |
|  | GLA protein | -584.300 | -2442 to 1273 | 0.489 |  |  |  |  |  |
|  | GLA activity | -20.590 | -115.3 to 74.09 | 0.630 |  |  |  |  |  |
|  | Age | 29.060 | 7.409 to 50.71 | 0.015 |  |  |  |  |  |
| 3 | Intercept | -453.600 | -1589 to 681.4 | 0.390 | 0.58 | 0.44 | 177.4 | F(3,9)=<br>4.1 | 0.044 |
|  | Hsp60 | 752.000 | -1055 to 2558 | 0.371 |  |  |  |  |  |
|  | GLA protein | -806.900 | -2263 to 649.2 | 0.242 |  |  |  |  |  |
|  | Age | 30.120 | 10.30 to 49.93 | 0.007 |  |  |  |  |  |
| 4 | Intercept | -230.000 | -1209 to 749.2 | 0.612 | 0.54 | 0.44 | 173.1 | F(2,10)=<br>5.8 | 0.022 |
|  | GLA protein | -280.800 | -989.1 to 427.5 | 0.398 |  |  |  |  |  |
|  | Age | 27.440 | 9.086 to 45.79 | 0.008 |  |  |  |  |  |
| 5 | Intercept | -0.124 | -1004 to 1004 | 1.000 | 0.36 | 0.32 | 244.3 | F(1,16)=<br>8.8 | 0.009 |
|  | Age | 27.020 | 7.710 to 46.32 | 0.009 |  |  |  |  |  |

**Table H.**
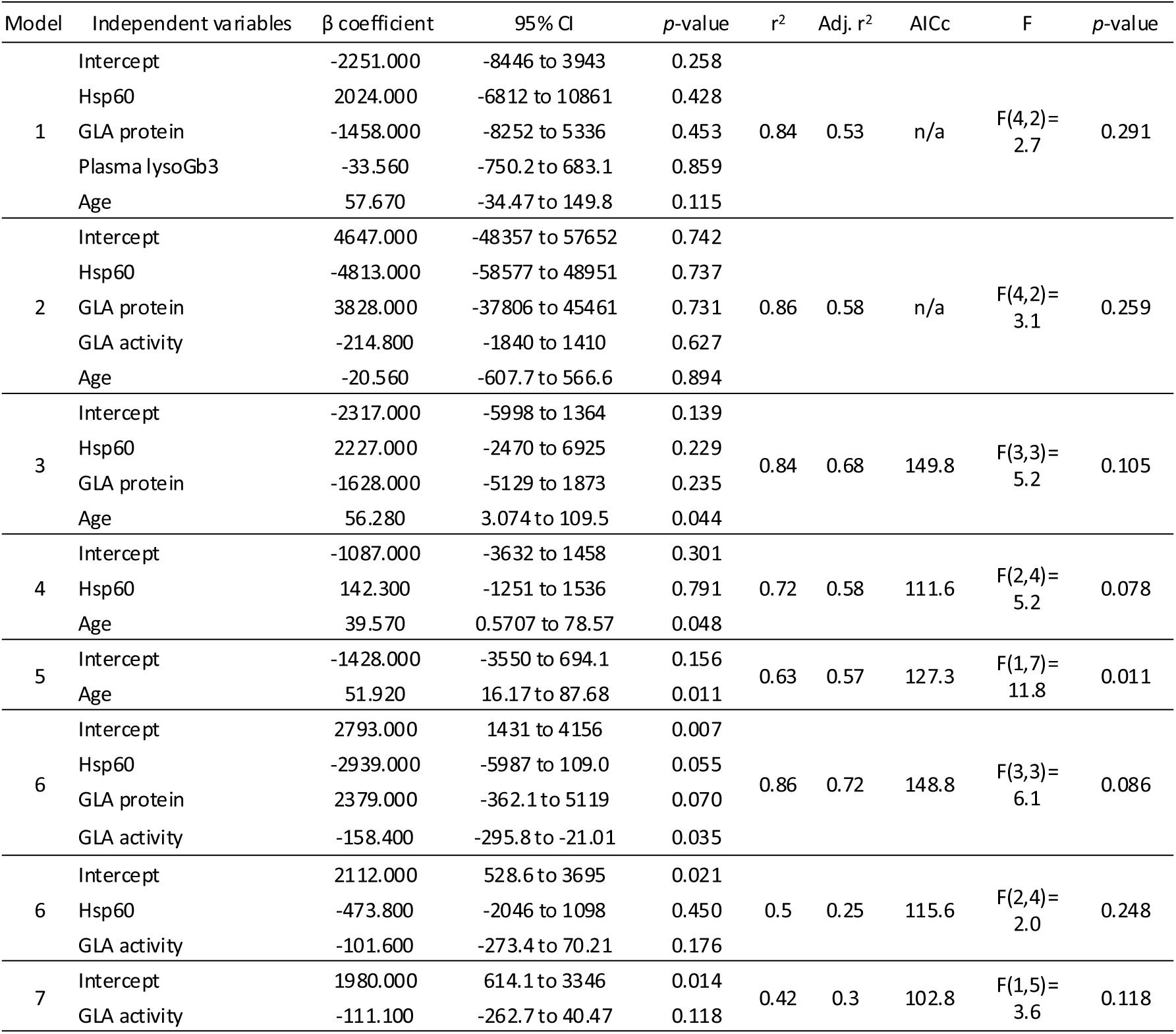
Multiple linear regression analysis for GDF-15 - For N215S males.

In terms of overall clinical severity, data for males showed positive statistically significant correlations between the MSSI and both mitokines (FGF-21: r2= 0.59 p= 0.01, fig. 9, and GDF-15: r2= 0.67 p= 0.002, fig. 10). When looking at the N215S male group, the correlations between both severity scores and FGF-21 strengthened, suggesting that higher levels of these mitokines were found in subjects with higher severity scores for this genotype group (AASS: r2= 0.65 p= 0.06 and MSSI: r2= 0.87 p= 0.005, fig. 9). For females, no correlation was found between levels of serum mitokines and overall clinical severity scores (figures 9 and 10).

**Fig. 9.**
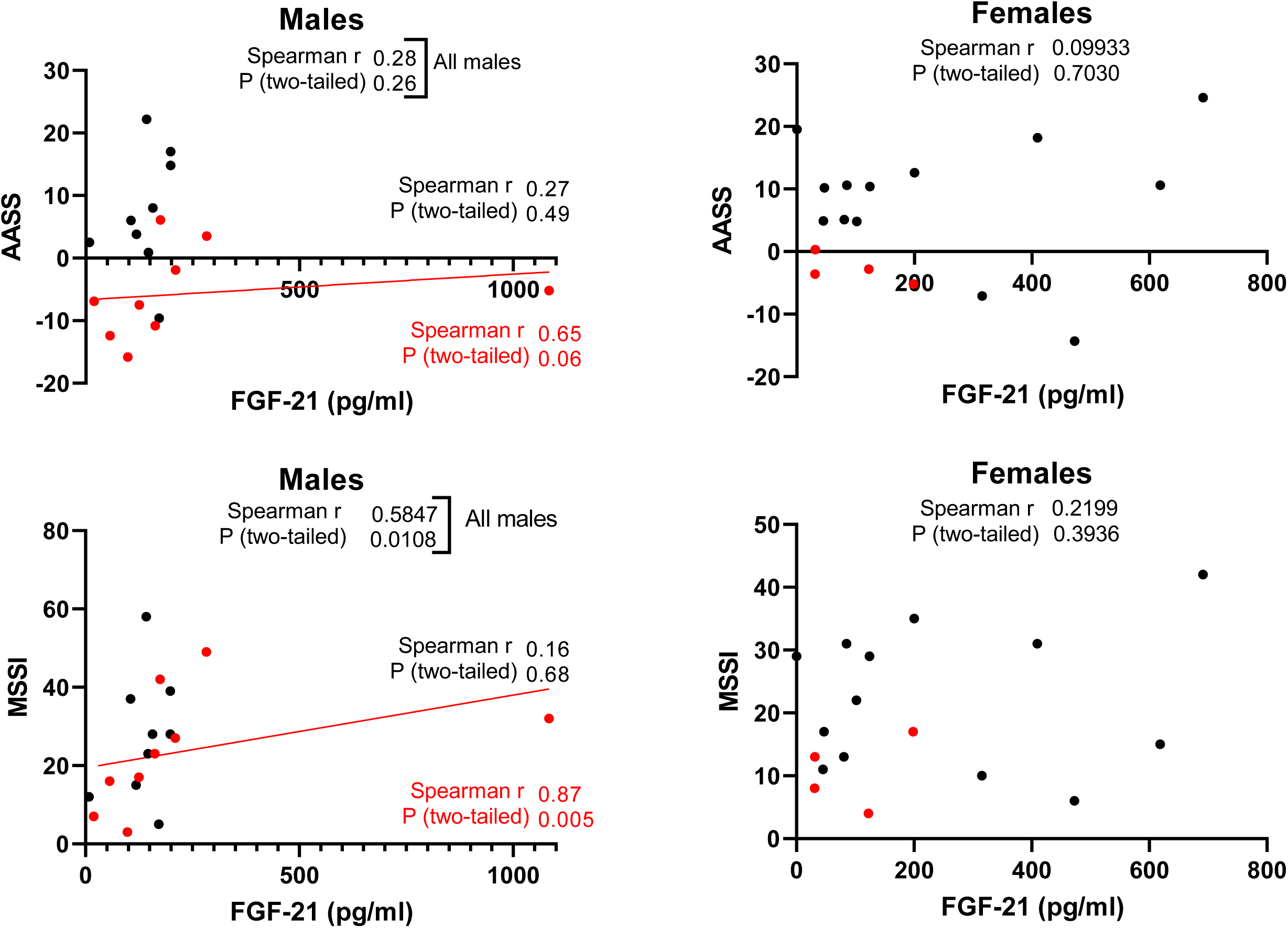
Relationship between serum FGF-21 levels and clinical severity by the current Mainz Severity Score Index (MSSI) and the age-adjusting severity score (AASS).

**Fig. 10.**
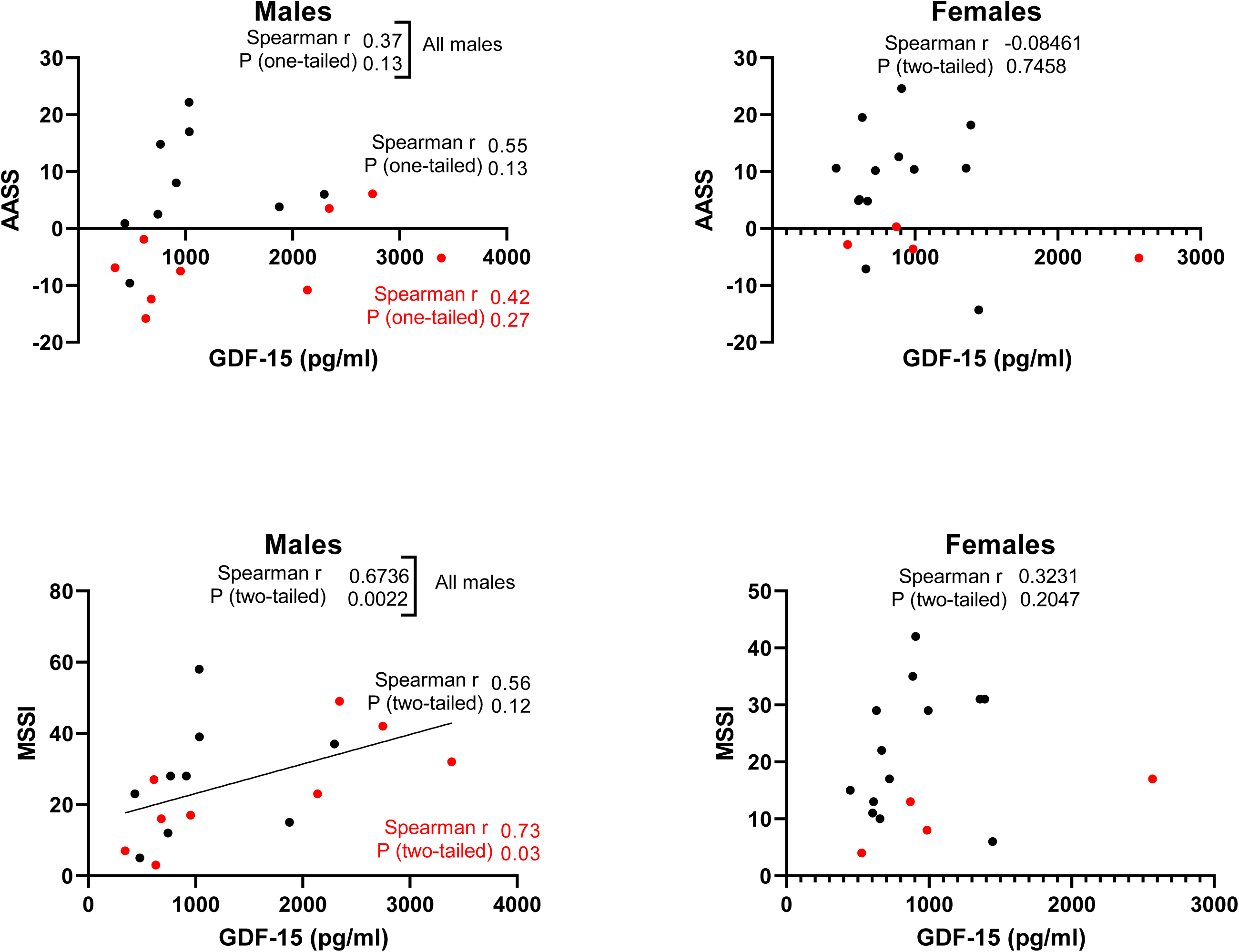
Relationship between serum GDF-15 levels and clinical severity by the current Mainz Severity Score Index (MSSI) and the age-adjusting severity score (AASS).

When patients were grouped according to the presence of target organ injury, i.e., FD with or without cardiomyopathy and nephropathy (as defined in the methods section), serum levels of GDF-15 were significantly higher in those patients with cardiomyopathy, associated or not with nephropathy, than in patients without cardiomyopathy (934.9 vs 610.1 pg/ml, p= 0.02, fig. 11), in agreement with published data (Gregório et al. 2022). GDF-15 levels were also higher in those individuals who developed either a serious renal event (1827 vs 743.3 pg/ml, p= 0.007, fig. 11) or a serious cardiac event (1197 vs 723.3 pg/ml, p= 0.001, fig. 12), see tables 1 and 2 for individual patient data. When participants were classified upon the presence of LVH, both mitokines were higher in those with LVH documented on an image scan (FGF-21: 159.7 vs 98.3 pg/ml, p= 0.04, and GDF-15: 974.6 vs 655.8 pg/ml, p= 0.01, fig. 12).

**Fig. 11.**
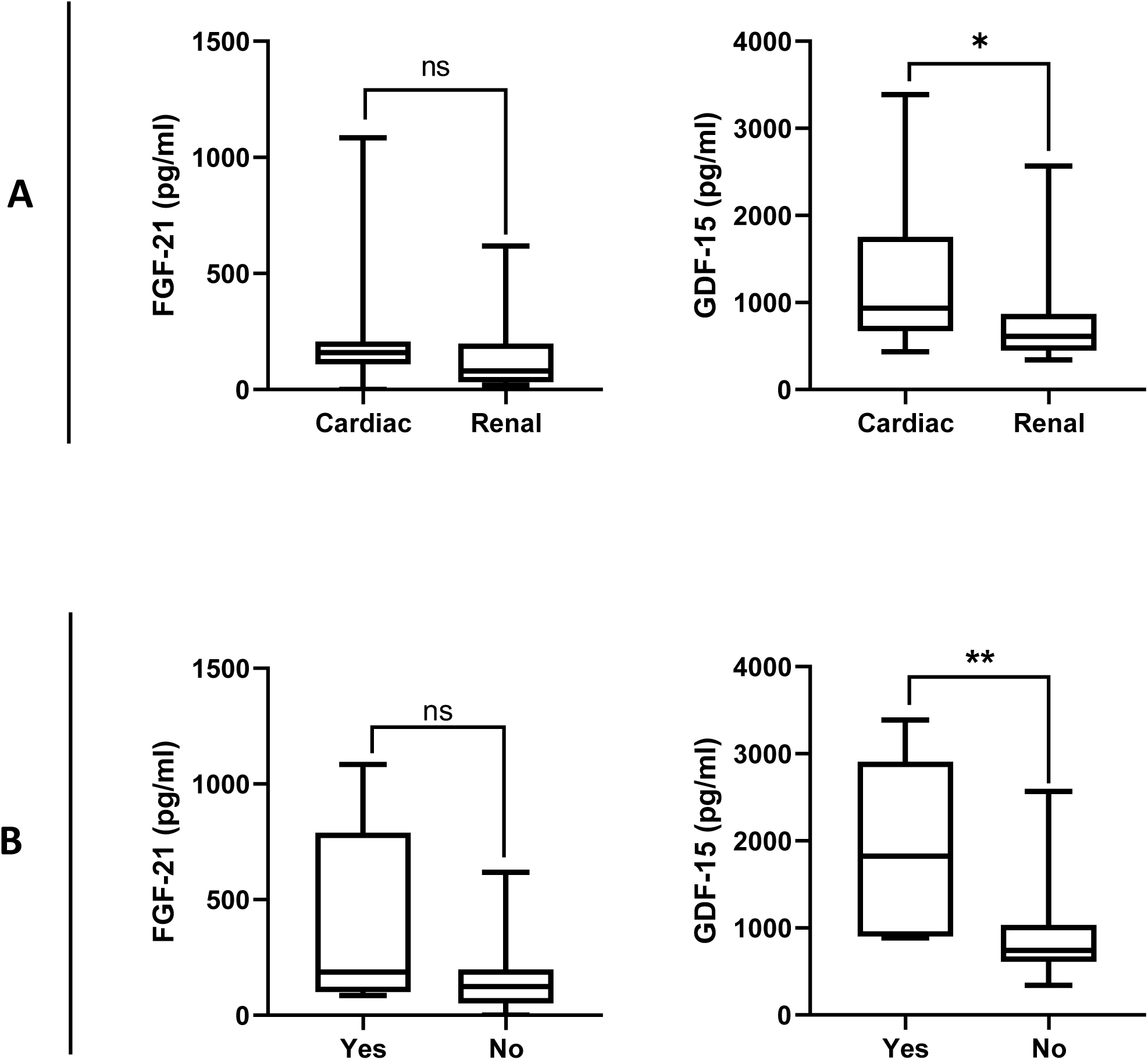
Serum levels of fibroblast growth factor 21 (FGF-21) and growth differentiation factor 15 (GDF-15) in participants with Fabry disease with or without (A) cardiomyopathy and nephropathy, and (B) the development of a serious renal event. FGF-21 and GDF-15 were measured by enzyme-linked immunosorbent assay (ELISA) in patients with Fabry disease (FD, males n= 18, females n= 17). Only 7 patients had nephropathy without Associated cardiomyopathy. Male participant 16 and females number 26, 28, and 19 have not shown signs of neither nephropathy or cardiomyopathy. Serious renal event as defined in the methods section. Data per participant is on tables 1 and 2.

**Fig. 12.**
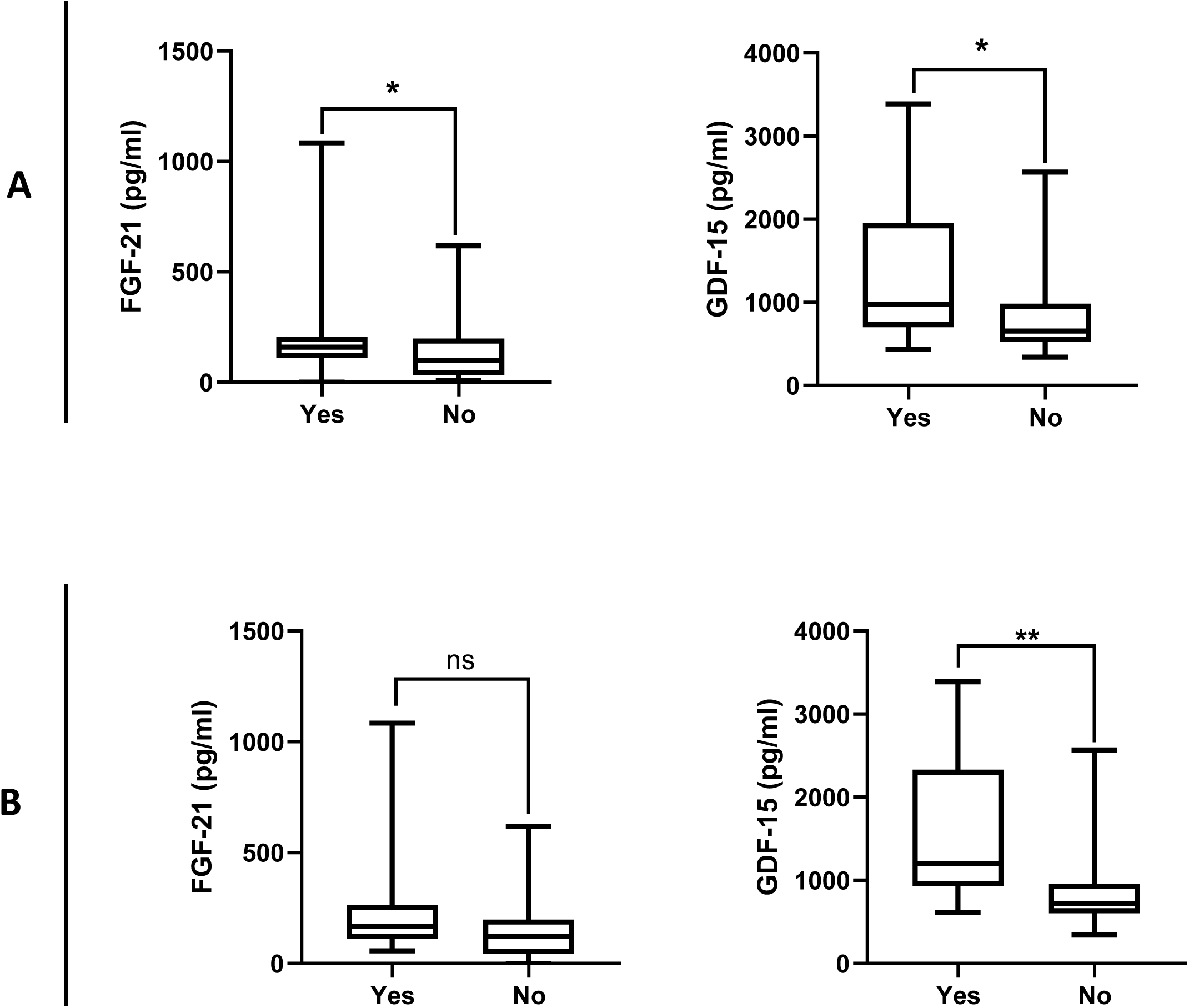
Serum levels of fibroblast growth factor 21 (FGF-21) and growth differentiation factor 15 (GDF-15) in participants with Fabry disease with or without (A) left ventricular hypertrophy (LVH) on image, and (B) the development of a serious cardiac event. FGF-21 and GDF-15 were measured by enzyme-linked immunosorbent assay (ELISA) in patients with Fabry disease (FD, males n= 18, females n= 17). Serious cardiac event as defined in the methods section. Data per participant is on tables 1 and 2.

Exploratory multivariate linear regression modelling including data for all males, indicated that 76% of MSSI variation was associated positively to GDF-15 (β= 0.01 pg/ml, 95% CI= 0.003 to 0.02, p= 0.009), GLA protein (β= 37.3 GLA/LC, 95% CI= 13.8 to 60.8, p= 0.006), and negatively to Hsp60 (β= -52.2 Hsp60/LC, 95% CI= -81.3 to -23.1, p= 0.003)(table J, model 4, AICc= 71.5, F(3,9)= 9.5, p= 0.004). For N215S males, a statistically significant model was found (table K, model 4, AICc= 47.5, F (2,4) = 22.5, p= 0.007), where the only statistically significant independent variable was FGF-21 (β= 0.12 pg/ml, 95% CI= 0.05 to 0.19, p= 0.01, table K) and explained 92% of MSSI variance for this group. For females, two models explained MSSI variance, model 6 explained 33% and the only independent variable was GLA activity (β= -0.56 nmol/hr/mg, 95% CI= -1.11 to 0.01, p= 0.05, table L, model 6). Additionally, model 10 explained 72% of MSSI variance, and the only significant independent variable was plasma lyso-Gb3 (β= 3.1 ng/ml, 95% CI= 1.14 to 4.95, p= 0.008, table L, model 10). Statistically significant multivariate regression models for the AASS were found only for females, all including plasma lyso-Gb3 as the only statistically significant independent variable (table O, models 8, 9, and 10).

**Table I.** Multiple linear regression analysis for GDF-15 - All female cohort.

| Model | Independent variables | β coefficient | 95% CI | p-value | r<sup>2</sup> | Adj. r<sup>2</sup> | AICc | F | p-value |
| --- | --- | --- | --- | --- | --- | --- | --- | --- | --- |
| 1 | Intercept | -200.000 | -3346 to 2946 | 0.882 | 0.55 | 0.24 | 164.6 | F(4,6)=<br>1.8 | 0.247 |
|  | Hsp60 | -247.600 | -5486 to 4990 | 0.912 |  |  |  |  |  |
|  | GLA protein | 136.700 | -1247 to 1520 | 0.817 |  |  |  |  |  |
|  | GLA activity | -4.230 | -76.49 to 68.03 | 0.891 |  |  |  |  |  |
|  | Age | 24.980 | -12.99 to 62.95 | 0.159 |  |  |  |  |  |
| 2 | Intercept | -307.200 | -2259 to 1645 | 0.721 | 0.55 | 0.35 | 153.6 | F(3,7)=<br>2.8 | 0.119 |
|  | GLA protein | 125.000 | -1094 to 1344 | 0.815 |  |  |  |  |  |
|  | GLA activity | -3.580 | -67.12 to 59.96 | 0.898 |  |  |  |  |  |
|  | Age | 26.110 | -0.3397 to 52.56 | 0.052 |  |  |  |  |  |
| 4 | Intercept | -291.200 | -1876 to 1294 | 0.687 | 0.47 | 0.35 | 158.1 | F(2,9)=<br>4.0 | 0.058 |
|  | GLA protein | 93.830 | -647.3 to 834.9 | 0.781 |  |  |  |  |  |
|  | Age | 23.810 | 0.5232 to 47.10 | 0.046 |  |  |  |  |  |
| 5 | Intercept | -53.880 | -709.9 to 602.2 | 0.863 | 0.44 | 0.41 | 209.1 | F(1,5)=<br>12.0 | 0.0035 |
|  | Age | 20.150 | 7.725 to 32.57 | 0.004 |  |  |  |  |  |

**Table J.** Multiple linear regression analysis for MSSI - All male cohort.

| Model | Independent variables | $\beta$ coefficient | 95% CI | $p$-value | $r^2$ | Adj. $r^2$ | AICc | F | $p$-value |
| --- | --- | --- | --- | --- | --- | --- | --- | --- | --- |
| 1 | Intercept | 24.75 | -16.81 to 66.31 | 0.1954 | 0.82 | 0.63 | 101.5 | F(6,6)= 4.4 | 0.046 |
|  | Age | 0.15 | -0.47 to 0.76 | 0.5833 |  |  |  |  |  |
|  | GDF15 | 0.01 | -0.01 to 0.03 | 0.1501 |  |  |  |  |  |
|  | FGF21 | -0.07 | -0.24 to 0.09 | 0.3474 |  |  |  |  |  |
|  | Hsp60 | -64.47 | -122.30 to -6.69 | 0.0342 |  |  |  |  |  |
|  | GLA protein | 44.53 | -1.21 to 90.27 | 0.0546 |  |  |  |  |  |
|  | GLA activity | -0.22 | -2.03 to 1.60 | 0.7783 |  |  |  |  |  |
| 2 | Intercept | 22.37 | -10.59 to 55.34 | 0.1525 | 0.81 | 0.68 | 86.1 | F(5,7)= 6.1 | 0.017 |
|  | Age | 0.15 | -0.40 to 0.70 | 0.5392 |  |  |  |  |  |
|  | GDF15 | 0.01 | -0.003 to 0.02 | 0.1164 |  |  |  |  |  |
|  | FGF21 | -0.07 | -0.21 to 0.08 | 0.3279 |  |  |  |  |  |
|  | Hsp60 | -61.97 | -110.50 to -13.40 | 0.0195 |  |  |  |  |  |
|  | GLA protein | 41.83 | 5.91 to 77.74 | 0.0284 |  |  |  |  |  |
| 3 | Intercept | 28.55 | 6.03 to 51.06 | 0.0192 | 0.80 | 0.70 | 76.5 | F(4,8)= 8.1 | 0.006 |
|  | GDF15 | 0.01 | 0.005 to 0.02 | 0.0066 |  |  |  |  |  |
|  | FGF21 | -0.08 | -0.21 to 0.05 | 0.2284 |  |  |  |  |  |
|  | Hsp60 | -68.21 | -108.40 to -27.97 | 0.0045 |  |  |  |  |  |
|  | GLA protein | 46.96 | 18.22 to 75.70 | 0.0055 |  |  |  |  |  |
| 4 | Intercept | 17.79 | 5.51 to 30.07 | 0.0095 | 0.76 | 0.68 | 71.5 | F(3,9)= 9.5 | 0.004 |
|  | GDF15 | 0.01 | 0.003 to 0.02 | 0.0091 |  |  |  |  |  |
|  | Hsp60 | -52.19 | -81.32 to -23.07 | 0.0029 |  |  |  |  |  |
|  | GLA protein | 37.29 | 13.76 to 60.83 | 0.0059 |  |  |  |  |  |
| 5 | Intercept | 19.33 | -22.54 to 61.20 | 0.2887 | 0.85 | 0.66 | 108.6 | F(6,5)= 4.5 | 0.059 |
|  | Age | 0.14 | -0.5115 to 0.7927 | 0.6033 |  |  |  |  |  |
|  | GDF15 | 0.01 | -0.01 to 0.03 | 0.1367 |  |  |  |  |  |
|  | FGF21 | -0.07 | -0.25 to 0.11 | 0.3732 |  |  |  |  |  |
|  | Hsp60 | -59.63 | -121.6 to 2.30 | 0.0562 |  |  |  |  |  |
|  | GLA protein | 41.69 | -3.27 to 86.65 | 0.0629 |  |  |  |  |  |
|  | Plasma lyso-Gb3 | 0.14 | -0.50 to 0.78 | 0.6013 |  |  |  |  |  |
| 6 | Intercept | 25.18 | -3.351 to 53.72 | 0.0741 | 0.84 | 0.70 | 87.3 | F(5,6)= 6.1 | 0.024 |
|  | GDF15 | 0.02 | 0.005 to 0.03 | 0.0116 |  |  |  |  |  |
|  | FGF21 | -0.08 | -0.23 to 0.07 | 0.2618 |  |  |  |  |  |
|  | Hsp60 | -65.71 | -115.1 to -16.37 | 0.0173 |  |  |  |  |  |
|  | GLA protein | 46.67 | 12.15 to 81.20 | 0.0162 |  |  |  |  |  |
|  | Plasma lyso-Gb3 | 0.13 | -0.44 to 0.71 | 0.5899 |  |  |  |  |  |

**Table K.**
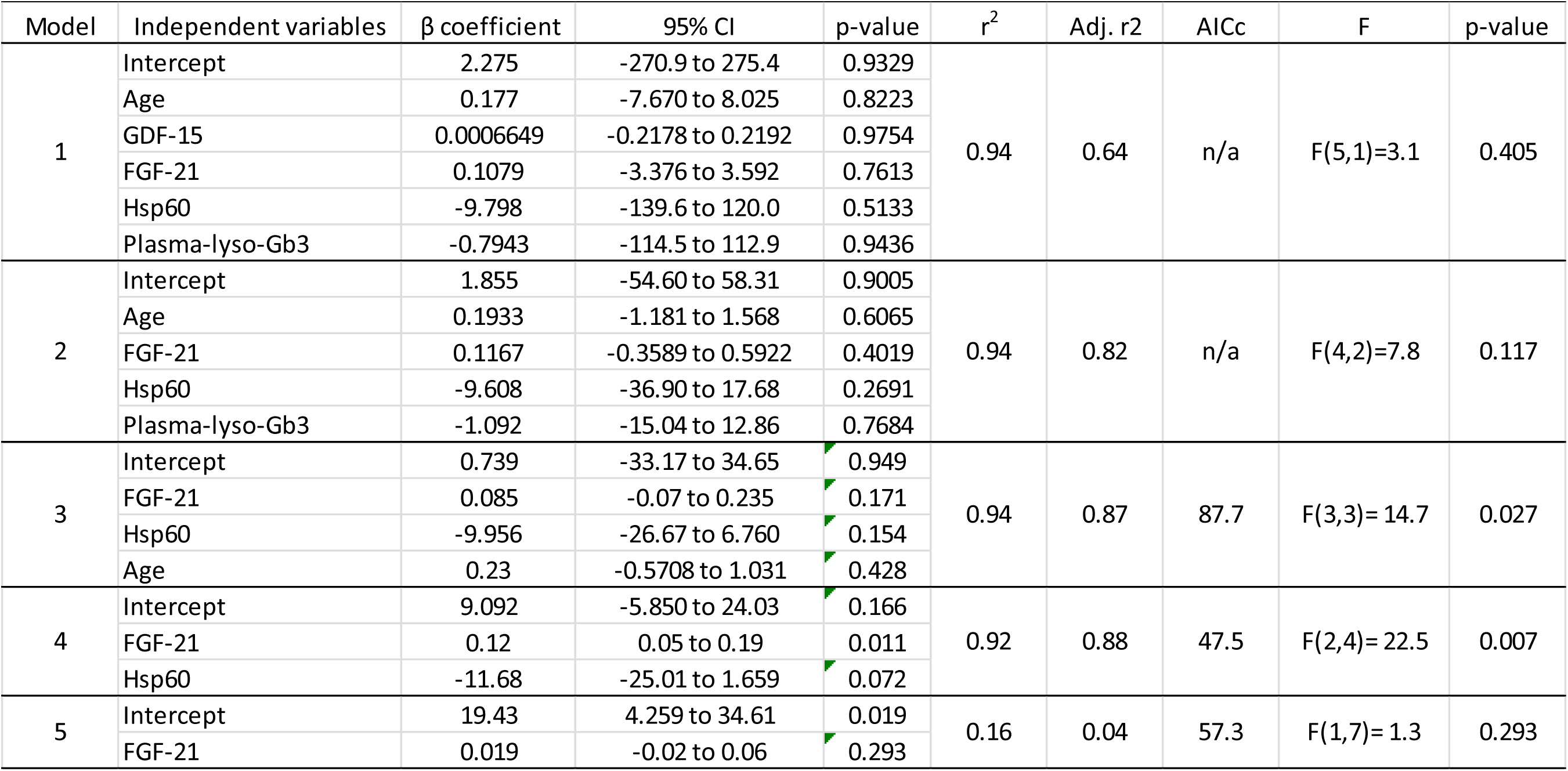
Multiple linear regression analysis for MSSI - N215S males.

| Table K. Multiple linear regression analysis for MSSl - N215S males |  |  |  |  |  |  |  |  |  |
| --- | --- | --- | --- | --- | --- | --- | --- | --- | --- |
| Model | Independent variables | β coefficient | 95% CI | p-value | r<sup>2</sup> | Adj. r2 | AICc | F | p-value |
| 1 | Intercept | 2.275 | -270.9 to 275.4 | 0.9329 | 0.94 | 0.64 | n/a | F(5,1)=3.1 | 0.405 |
|  | Age | 0.177 | -7.670 to 8.025 | 0.8223 |  |  |  |  |  |
|  | GDF-15 | 0.0006649 | -0.2178 to 0.2192 | 0.9754 |  |  |  |  |  |
|  | FGF-21 | 0.1079 | -3.376 to 3.592 | 0.7613 |  |  |  |  |  |
|  | Hsp60 | -9.798 | -139.6 to 120.0 | 0.5133 |  |  |  |  |  |
|  | Plasma-lyso-Gb3 | -0.7943 | -114.5 to 112.9 | 0.9436 |  |  |  |  |  |
| 2 | Intercept | 1.855 | -54.60 to 58.31 | 0.9005 | 0.94 | 0.82 | n/a | F(4,2)=7.8 | 0.117 |
|  | Age | 0.1933 | -1.181 to 1.568 | 0.6065 |  |  |  |  |  |
|  | FGF-21 | 0.1167 | -0.3589 to 0.5922 | 0.4019 |  |  |  |  |  |
|  | Hsp60 | -9.608 | -36.90 to 17.68 | 0.2691 |  |  |  |  |  |
|  | Plasma-lyso-Gb3 | -1.092 | -15.04 to 12.86 | 0.7684 |  |  |  |  |  |
| 3 | Intercept | 0.739 | -33.17 to 34.65 | 0.949 | 0.94 | 0.87 | 87.7 | F(3,3)= 14.7 | 0.027 |
|  | FGF-21 | 0.085 | -0.07 to 0.235 | 0.171 |  |  |  |  |  |
|  | Hsp60 | -9.956 | -26.67 to 6.760 | 0.154 |  |  |  |  |  |
|  | Age | 0.23 | -0.5708 to 1.031 | 0.428 |  |  |  |  |  |
| 4 | Intercept | 9.092 | -5.850 to 24.03 | 0.166 | 0.92 | 0.88 | 47.5 | F(2,4)= 22.5 | 0.007 |
|  | FGF-21 | 0.12 | 0.05 to 0.19 | 0.011 |  |  |  |  |  |
|  | Hsp60 | -11.68 | -25.01 to 1.659 | 0.072 |  |  |  |  |  |
| 5 | Intercept | 19.43 | 4.259 to 34.61 | 0.019 | 0.16 | 0.04 | 57.3 | F(1,7)= 1.3 | 0.293 |
|  | FGF-21 | 0.019 | -0.02 to 0.06 | 0.293 |  |  |  |  |  |

**Table L.** Multiple linear regression analysis for MSSI - Female cohort.

| Table L. Multiple linear regression analysis for MSSl - Female cohort |  |  |  |  |  |  |  |  |  |
| --- | --- | --- | --- | --- | --- | --- | --- | --- | --- |
| Model | Independent variables | $\beta$ coefficient | 95% CI | $p$-value | $r^2$ | Adj. $r^2$ | AIC | F | $p$-value |
| 1 | Intercept | 8.62 | -44.50 to 61.74 | 0.6757 | 0.58 | -0.05 | 122.2 | F(6,4)= 0.9 | 0.557 |
|  | Age | 0.26 | -0.50 to 1.01 | 0.4032 |  |  |  |  |  |
|  | GDF15 | 0.00 | -0.01 to 0.01 | 0.7341 |  |  |  |  |  |
|  | FGF21 | 0.00 | -0.03 to 0.03 | 0.8992 |  |  |  |  |  |
|  | Hsp60 | 11.63 | -76.64 to 99.91 | 0.733 |  |  |  |  |  |
|  | GLA protein | 13.77 | -9.42 to 36.97 | 0.1747 |  |  |  |  |  |
|  | GLA activity | -0.79 | -2.02 to 0.43 | 0.1472 |  |  |  |  |  |
| 2 | Intercept | 8.22 | -35.34 to 51.78 | 0.6481 | 0.58 | 0.16 | 85.6 | F(5,5)= 1.4 | 0.367 |
|  | Age | 0.26 | -0.36 to 0.88 | 0.3362 |  |  |  |  |  |
|  | GDF15 | 0.00 | -0.01 to 0.01 | 0.6913 |  |  |  |  |  |
|  | Hsp60 | 12.26 | -60.21 to 84.73 | 0.6817 |  |  |  |  |  |
|  | GLA protein | 13.84 | -5.37 to 33.06 | 0.1234 |  |  |  |  |  |
|  | GLA activity | -0.80 | -1.80 to 0.19 | 0.0936 |  |  |  |  |  |
| 3 | Intercept | 8.67 | -29.77 to 47.11 | 0.6009 | 0.57 | 0.28 | 67.7 | F(4,6)= 1.9 | 0.222 |
|  | Age | 0.20 | -0.26 to 0.66 | 0.3249 |  |  |  |  |  |
|  | Hsp60 | 12.82 | -51.19 to 76.83 | 0.6414 |  |  |  |  |  |
|  | GLA protein | 13.53 | -3.38 to 30.44 | 0.098 |  |  |  |  |  |
|  | GLA activity | -0.79 | -1.67 to 0.08 | 0.0699 |  |  |  |  |  |
| 4 | Intercept | 14.23 | -10.07 to 38.53 | 0.2088 | 0.55 | 0.35 | 57.1 | F(3,7)= 2.8 | 0.117 |
|  | Age | 0.14 | -0.18 to 0.47 | 0.3327 |  |  |  |  |  |
|  | GLA protein | 14.14 | -1.03 to 29.32 | 0.0635 |  |  |  |  |  |
|  | GLA activity | -0.83 | -1.6 to -0.03 | 0.0424 |  |  |  |  |  |
| 5 | Intercept | 23.88 | 13.66 to 34.11 | 0.0007 | 0.48 | 0.35 | 51.3 | F(2,8)= 3.7 | 0.075 |
|  | GLA protein | 12.17 | -2.04 to 26.39 | 0.0838 |  |  |  |  |  |
|  | GLA activity | -0.88 | -1.64 to -0.11 | 0.0288 |  |  |  |  |  |
| 6 | Intercept | 27.43 | 14.83 to 40.04 | 0.0007 | 0.33 | 0.26 | 60.5 | F(1,10)= 4.8 | 0.053 |
|  | GLA activity | -0.56 | -1.11 to 0.01 | 0.0525 |  |  |  |  |  |
| 7 | Intercept | -2.24 | -34.52 to 30.03 | 0.8392 | 0.85 | 0.66 | 115.3 | F(4,3)= 4.3 | 0.129 |
|  | Age | 0.20 | -0.4497 to 0.8463 | 0.402 |  |  |  |  |  |
|  | GDF15 | 0.00 | -0.01398 to 0.01429 | 0.974 |  |  |  |  |  |
|  | FGF21 | 0.01 | -0.02084 to 0.04997 | 0.2817 |  |  |  |  |  |
|  | Plasma lyso-Gb3 | 3.38 | 0.7424 to 6.024 | 0.0266 |  |  |  |  |  |
| 8 | Intercept | -2.36 | -25.33 to 20.60 | 0.7893 | 0.85 | 0.74 | 59.3 | F(3,4)= 7.7 | 0.039 |
|  | Age | 0.20 | -0.1407 to 0.5475 | 0.1761 |  |  |  |  |  |
|  | FGF21 | 0.01 | -0.01116 to 0.04050 | 0.19 |  |  |  |  |  |
|  | Plasma lyso-Gb3 | 3.38 | 1.389 to 5.373 | 0.0092 |  |  |  |  |  |
| 9 | Intercept | 5.94 | -12.73 to 24.61 | 0.4504 | 0.76 | 0.66 | 44.5 | F(2,5)= 7.9 | 0.028 |
|  | Age | 0.12 | -0.2083 to 0.4452 | 0.3941 |  |  |  |  |  |
|  | Plasma lyso-Gb3 | 3.08 | 1.057 to 5.110 | 0.0113 |  |  |  |  |  |
| 10 | Intercept | 12.27 | 6.042 to 18.50 | 0.0029 | 0.72 | 0.67 | 36.4 | F(1,6)= 15.3 | 0.008 |
|  | Plasma lyso-Gb3 | 3.05 | 1.142 to 4.953 | 0.0079 |  |  |  |  |  |

**Table M.** Multiple linear regression analysis for AASS - All male cohort.

| Table M. Multiple linear regression analysis for AASS - All male cohort |  |  |  |  |  |  |  |  |  |
| --- | --- | --- | --- | --- | --- | --- | --- | --- | --- |
| Model | Independent variables | $\beta$ coefficient | 95% CI | $p$-value | $r^2$ | Adj. $r^2$ | AICc | F | $p$-value |
| 1 | Intercept | 25.17 | -16.72 to 67.07 | 0.1918 | 0.61 | 0.23 | 101.7 | F(6,6)= 1.6 | 0.296 |
|  | Age | -0.30 | -0.9203 to 0.3239 | 0.2853 |  |  |  |  |  |
|  | GDF15 | 0.01 | -0.007396 to 0.02713 | 0.2114 |  |  |  |  |  |
|  | FGF21 | -0.08 | -0.2491 to 0.08982 | 0.294 |  |  |  |  |  |
|  | Hsp60 | -63.76 | -122.0 to -5.523 | 0.0366 |  |  |  |  |  |
|  | GLA protein | 45.56 | -0.5485 to 91.67 | 0.052 |  |  |  |  |  |
|  | GLA activity | -0.40 | -2.237 to 1.432 | 0.6108 |  |  |  |  |  |
| 2 | Intercept | 20.81 | -12.96 to 54.59 | 0.1884 | 0.59 | 0.30 | 86.7 | F(5,7)=2.1 | 0.188 |
|  | Age | -0.29 | -0.8570 to 0.2798 | 0.269 |  |  |  |  |  |
|  | GDF15 | 0.01 | -0.005690 to 0.02588 | 0.1742 |  |  |  |  |  |
|  | FGF21 | -0.07 | -0.2245 to 0.07985 | 0.298 |  |  |  |  |  |
|  | Hsp60 | -59.16 | -108.9 to -9.400 | 0.0261 |  |  |  |  |  |
|  | GLA protein | 40.60 | 3.801 to 77.40 | 0.035 |  |  |  |  |  |
| 3 | Intercept | 8.48 | -12.95 to 29.92 | 0.3881 | 0.52 | 0.28 | 78.5 | F(4,8)=2.2 | 0.163 |
|  | Age | -0.22 | -0.7652 to 0.3248 | 0.3787 |  |  |  |  |  |
|  | GDF15 | 0.01 | -0.007220 to 0.02005 | 0.3095 |  |  |  |  |  |
|  | Hsp60 | -42.14 | -76.38 to -7.895 | 0.0219 |  |  |  |  |  |
|  | GLA protein | 29.71 | 1.168 to 58.25 | 0.0432 |  |  |  |  |  |
| 4 | Intercept | 1.61 | -11.08 to 14.30 | 0.781 | 0.47 | 0.29 | 72.4 | F(3,9)=2.6 | 0.113 |
|  | GDF15 | 0.002 | -0.006465 to 0.01099 | 0.5717 |  |  |  |  |  |
|  | Hsp60 | -36.19 | -66.30 to -6.083 | 0.0236 |  |  |  |  |  |
|  | GLA protein | 24.14 | -0.1963 to 48.47 | 0.0515 |  |  |  |  |  |
| 5 | Intercept | 3.02 | -7.717 to 13.75 | 0.5487 | 0.04 | -0.14 | 81.2 | F(2,11)=0.2 | 0.813 |
|  | Hsp60 | -7.46 | -38.53 to 23.62 | 0.608 |  |  |  |  |  |
|  | GLA protein | 1.86 | -24.27 to 28.00 | 0.8783 |  |  |  |  |  |
| 6 | Intercept | 16.94 | -21.52 to 55.40 | 0.309 | 0.71 | 0.37 | 106.6 | F(6,5)=2.1 | 0.219 |
|  | Age | -0.31 | -0.9047 to 0.2933 | 0.2465 |  |  |  |  |  |
|  | GDF15 | 0.01 | -0.005332 to 0.02816 | 0.1401 |  |  |  |  |  |
|  | FGF21 | -0.08 | -0.2399 to 0.08767 | 0.2858 |  |  |  |  |  |
|  | Hsp60 | -56.60 | -113.5 to 0.2817 | 0.0508 |  |  |  |  |  |
|  | GLA protein | 40.92 | -0.3729 to 82.22 | 0.0514 |  |  |  |  |  |
|  | Plasma lyso-Gb3 | 0.18 | -0.4085 to 0.7706 | 0.4656 |  |  |  |  |  |

**Table N.**
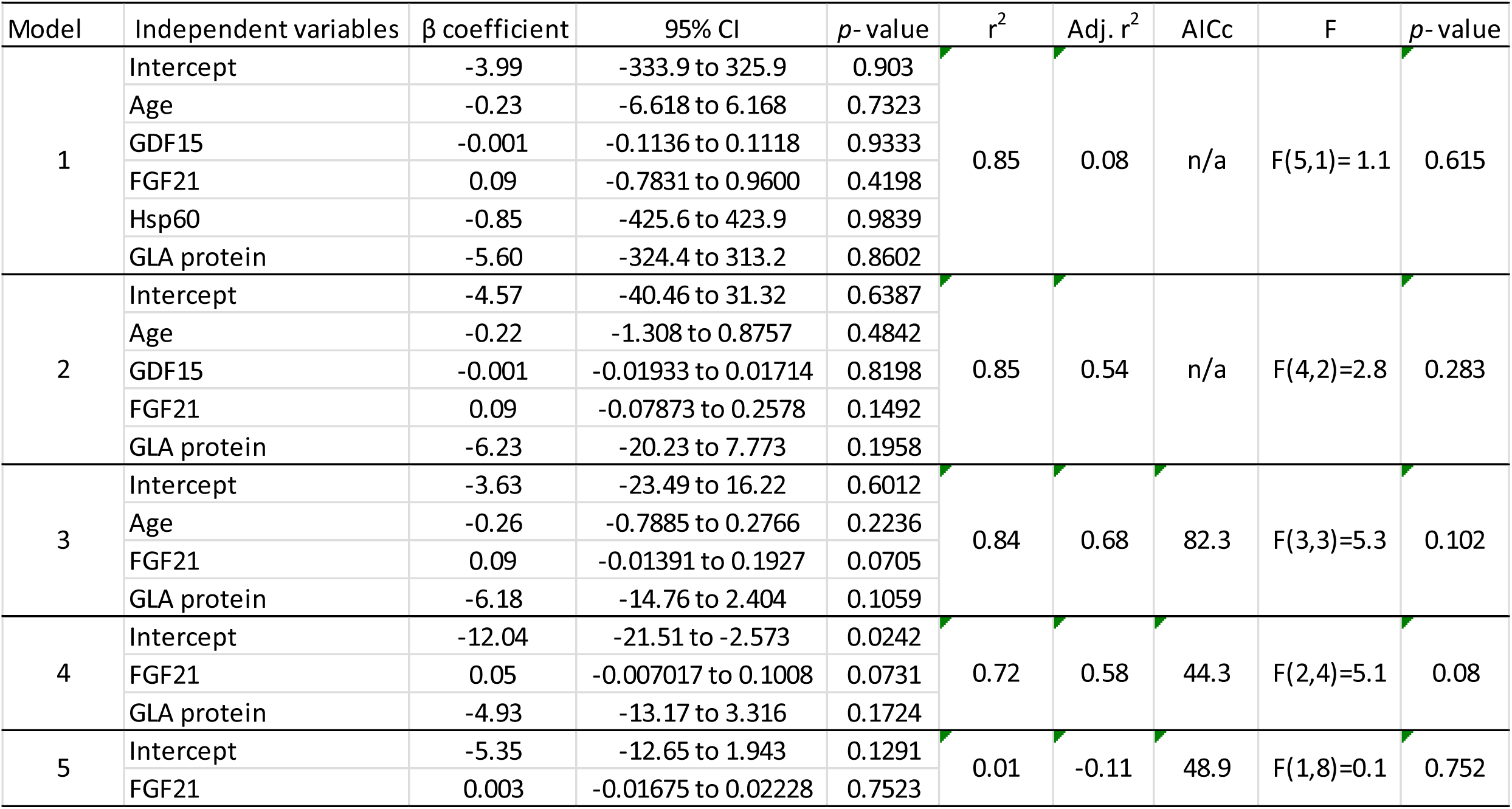
Multiple linear regression analysis for AASS - N215S males.

**Table O.** Multiple linear regression analysis for AASS - All female cohort.

|  |  |  |  |  |  |  |  |  |  |
| --- | --- | --- | --- | --- | --- | --- | --- | --- | --- |
| Model | Independent variables | $\beta$ coefficient | 95% CI | $p$- value | $r^2$ | Adj. $r^2$ | AICc | F | $p$- value |
| 1 | Intercept | 19.46 | -36.57 to 75.49 | 0.3894 | 0.59 | -0.02 | 123.4 | F(6,4)=1.0 | 0.538 |
|  | Age | -0.14 | -0.9443 to 0.6605 | 0.6491 |  |  |  |  |  |
|  | GDF15 | 0.00 | -0.02048 to 0.01474 | 0.6749 |  |  |  |  |  |
|  | FGF21 | -0.01 | -0.04407 to 0.03378 | 0.7322 |  |  |  |  |  |
|  | Hsp60 | -2.69 | -95.80 to 90.43 | 0.94 |  |  |  |  |  |
|  | GLA protein | 14.31 | -10.15 to 38.78 | 0.1796 |  |  |  |  |  |
|  | GLA activity | -0.85 | -2.141 to 0.4474 | 0.1434 |  |  |  |  |  |
| 2 | Intercept | 18.30 | -13.82 to 50.41 | 0.2029 | 0.59 | 0.18 | 86.7 | F(5,5)=1.5 | 0.347 |
|  | Age | -0.13 | -0.7038 to 0.4434 | 0.5849 |  |  |  |  |  |
|  | GDF15 | 0.00 | -0.01743 to 0.01174 | 0.637 |  |  |  |  |  |
|  | FGF21 | 0.00 | -0.03689 to 0.02693 | 0.7048 |  |  |  |  |  |
|  | GLA protein | 14.19 | -5.783 to 34.17 | 0.1274 |  |  |  |  |  |
|  | GLA activity | -0.84 | -1.900 to 0.2180 | 0.0967 |  |  |  |  |  |
| 3 | Intercept | 17.95 | -10.34 to 46.23 | 0.1715 | 0.58 | 0.30 | 68.8 | F(4,6)=2.1 | 0.204 |
|  | Age | -0.13 | -0.6331 to 0.3790 | 0.5616 |  |  |  |  |  |
|  | GDF15 | 0.00 | -0.01588 to 0.009765 | 0.5808 |  |  |  |  |  |
|  | GLA protein | 14.47 | -3.097 to 32.03 | 0.0905 |  |  |  |  |  |
|  | GLA activity | -0.88 | -1.790 to 0.03584 | 0.057 |  |  |  |  |  |
| 4 | Intercept | 18.88 | -6.873 to 44.64 | 0.1266 | 0.56 | 0.36 | 58.4 | F(3,7)=2.9 | 0.111 |
|  | Age | -0.207 | -0.5559 to 0.1421 | 0.2038 |  |  |  |  |  |
|  | GLA protein | 14.09 | -2.002 to 30.17 | 0.0772 |  |  |  |  |  |
|  | GLA activity | -0.87 | -1.704 to -0.02755 | 0.0446 |  |  |  |  |  |
| 5 | Intercept | 5.10 | -6.319 to 16.51 | 0.3334 | 0.43 | 0.29 | 53.8 | F(2,8)=3.0 | 0.106 |
|  | GLA protein | 16.89 | 1.018 to 32.76 | 0.0397 |  |  |  |  |  |
|  | GLA activity | -0.79 | -1.642 to 0.06729 | 0.0664 |  |  |  |  |  |
| 6 | Intercept | -0.19 | -9.147 to 8.760 | 0.9625 | 0.04 | -0.06 | 58.9 | F(1,10)=0.4 | 0.544 |
|  | GLA protein | 2.90 | -7.369 to 13.16 | 0.5436 |  |  |  |  |  |
| 7 | Intercept | 1.87 | -28.58 to 32.33 | 0.8573 | 0.90 | 0.77 | 114.3 | F(4,3)=6.7 | 0.08 |
|  | Age | -0.14 | -0.7547 to 0.4681 | 0.5098 |  |  |  |  |  |
|  | GDF15 | -0.000004 | -0.01334 to 0.01333 | 0.9992 |  |  |  |  |  |
|  | FGF21 | 0.01 | -0.02023 to 0.04659 | 0.2982 |  |  |  |  |  |
|  | Plasma lyso-Gb3 | 3.73 | 1.237 to 6.220 | 0.0176 |  |  |  |  |  |
| 8 | Intercept | 1.88 | -19.79 to 23.54 | 0.8218 | 0.90 | 0.83 | 58.3 | F(3,4)=12 | 0.02 |
|  | Age | -0.14 | -0.4681 to 0.1811 | 0.2871 |  |  |  |  |  |
|  | FGF21 | 0.01 | -0.01119 to 0.03755 | 0.2077 |  |  |  |  |  |
|  | Plasma lyso-Gb3 | 3.73 | 1.850 to 5.608 | 0.0053 |  |  |  |  |  |
| 9 | Intercept | -6.99 | -14.93 to 0.9532 | 0.0732 | 0.86 | 0.81 | 42.2 | F(2,5)=15.6 | 0.007 |
|  | FGF21 | 0.02 | -0.003465 to 0.03918 | 0.0839 |  |  |  |  |  |
|  | Plasma lyso-Gb3 | 3.86 | 2.057 to 5.662 | 0.0027 |  |  |  |  |  |
| 10 | Intercept | -2.40 | -9.343 to 4.537 | 0.4293 | 0.73 | 0.69 | 38.2 | F(1,6)=16.6 | 0.007 |
|  | Plasma lyso-Gb3 | 3.53 | 1.407 to 5.651 | 0.0066 |  |  |  |  |  |

**Table P.**
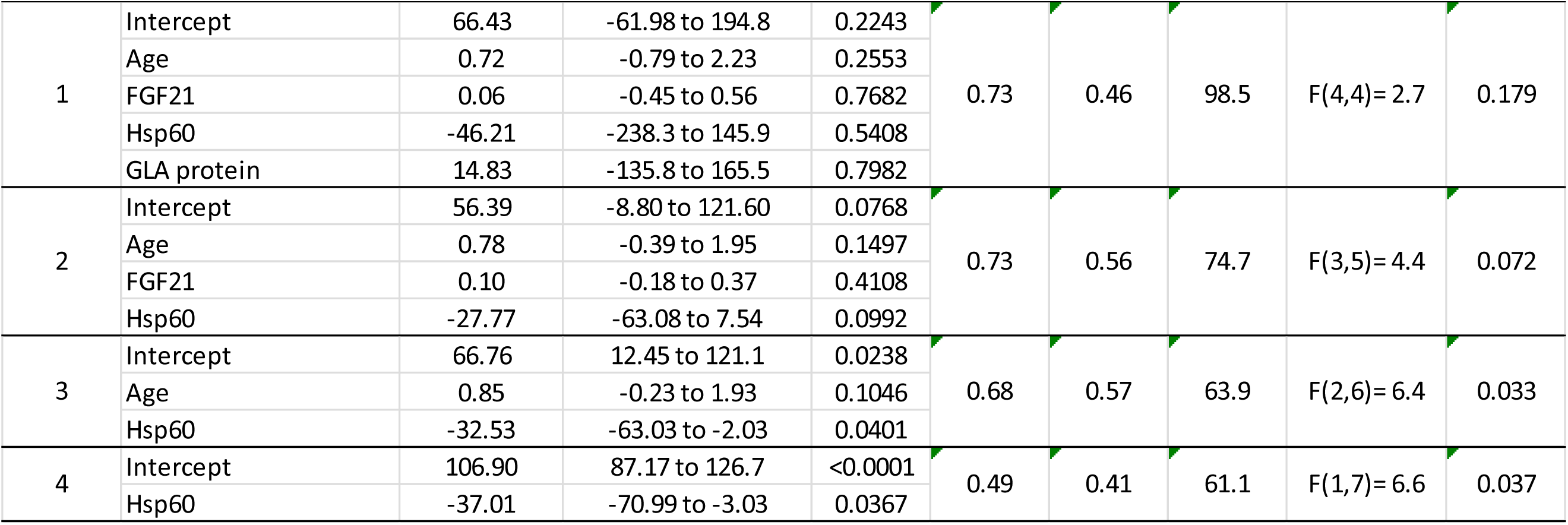
Multiple linear regression analysis for LVMI - All male cohort.

**Table Q.**
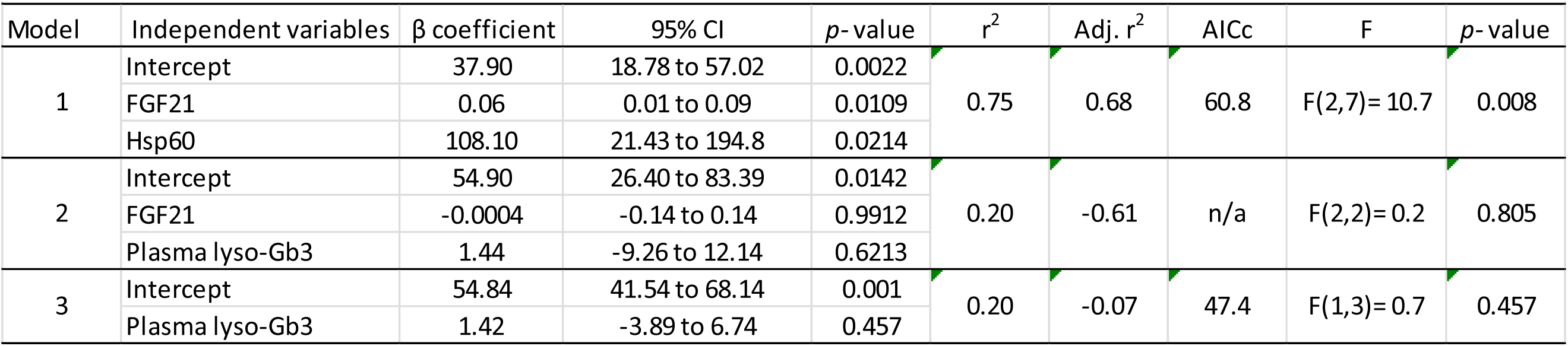
Multiple linear regression analysis for LVMI - All female cohort.

Correlation analysis between the mitokines and patients’ LVMIs suggested that higher serum levels of FGF-21 corresponded with higher LVMI magnitudes (males: r2= 0.66 p= 0.02 and females: r2= 0.47 p= 0.07, fig 13). For males, the best model only included one statistically significant independent variable, hsp60 and explained 49% of LVMI variance (AICc= 61.1, F (1,7) = 6.6, p= 0.04, model 4, table P). Conversely, for females the best model included both FGF-21 and Hsp60 as independent and significant variables, and explained 75% of LVMI variance (table Q, model 1). Correlation analyses between GDF-15 and GFR suggested higher levels of the mitokine corresponded with lower renal function rates for both sex groups (males: r2= -0.63 p= 0.005, females r2= -0.59 p= 0.01, fig. 14). FGF-21 correlation with GFR was only significant in males (r2= -0.56 p= 0.02, fig. 14). Multivariate modelling suggested higher GDF-15 serum values were significantly predictive of lower GFR only for males (β= - 0.02 pg/ml, 95% CI= -0.03 to -0.003, p= 0.02, table R, model 5), while for females this was age at the time the sample was taken (β= -0.68 years, 95% CI= -1.28 to -0.07, p= 0.03, table T, model 4). This model also indicated higher levels of serum FGF-21 related to higher GFR rates in females (β= 0.05 pg/ml, 95% CI= 0.002 to 0.092, p= 0.05, table Q, model 4).

**Fig. 13.**
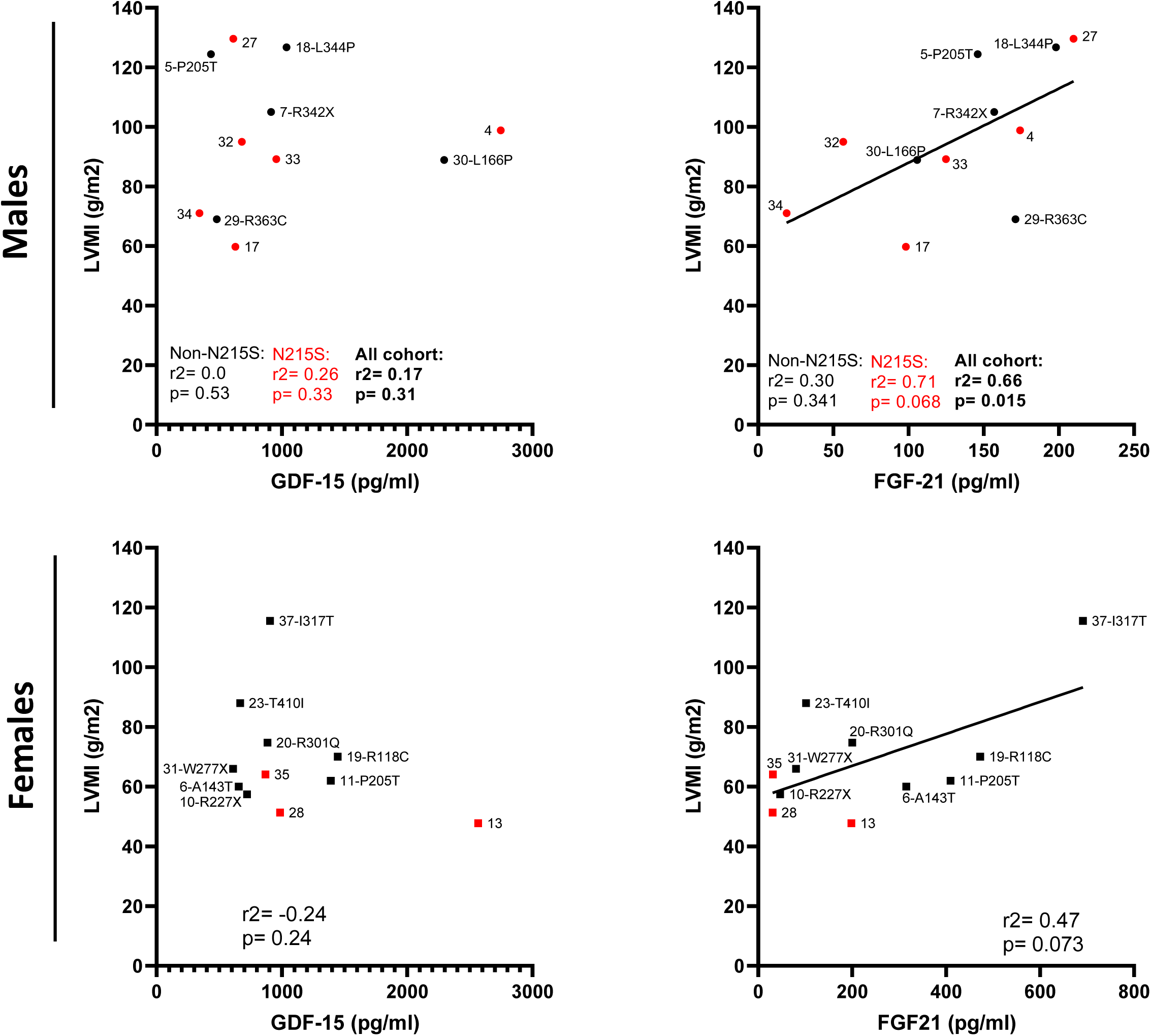
Relationship between serum levels of mitokines and left ventricular mass index (LVMI) in participants with Fabry disease. LVMI data was obtained via cardiac magnetic resonance (CMR) and corresponds to the most recently assessment available for each participant. In **red** = N215S

**Fig. 14.**
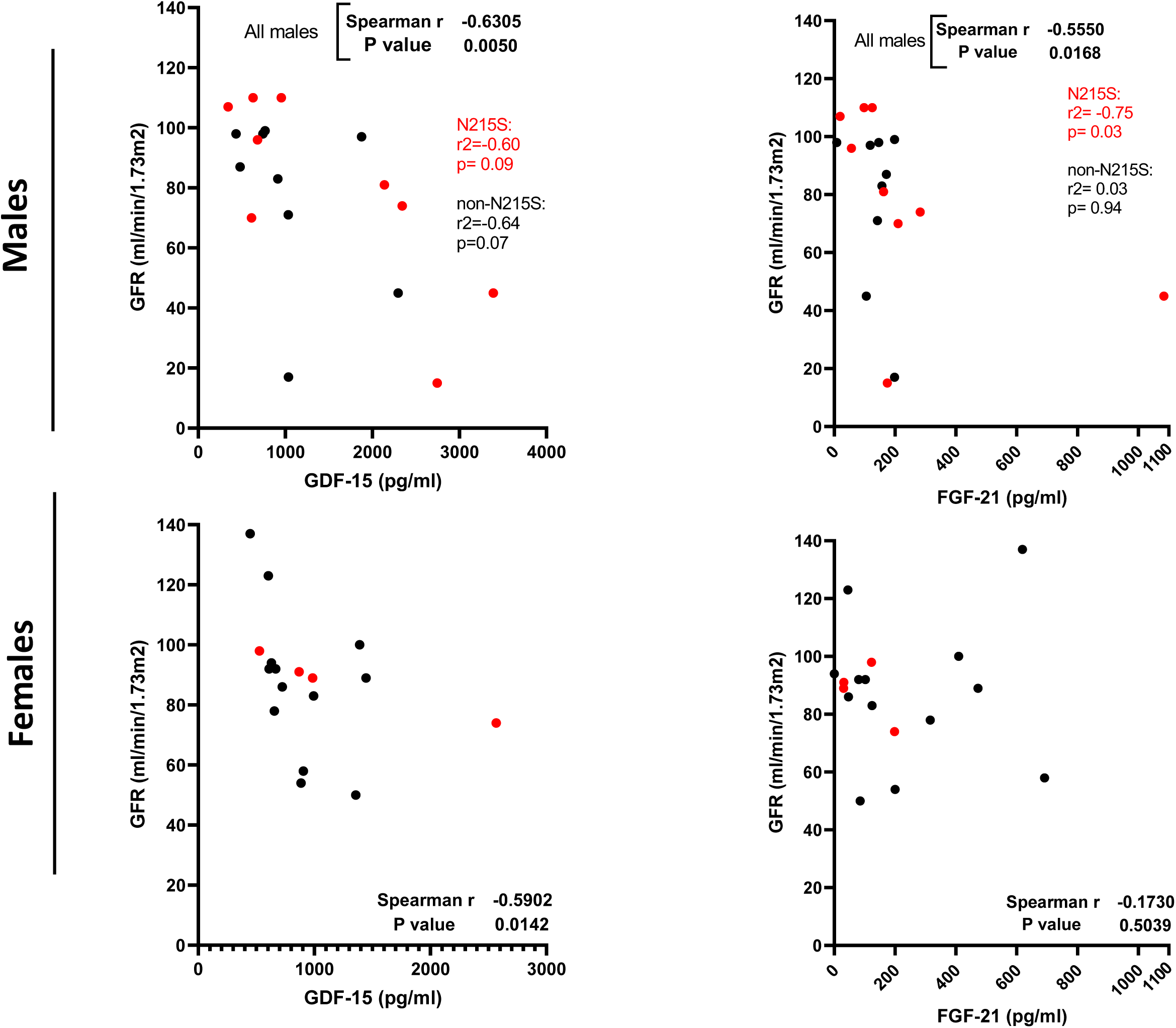
Relationship between serum levels of mitokines and current glomerular filtration rate (GFR) in participants with Fabry disease. In **red** = N215S individuals.

**Table R.** Multiple linear regression analysis for GFR - All male cohort.

| Table R. Multiple linear regression analysis for GFR - All male cohort |  |  |  |  |  |  |  |  |  |
| --- | --- | --- | --- | --- | --- | --- | --- | --- | --- |
| Model | Independent variables | $\beta$ coefficient | 95% CI | p-value | r<sup>2</sup> | Adj. r<sup>2</sup> | AICc | F | p-value |
| 1 | Intercept | 99.94 | 21.75 to 178.1 | 0.0204 | 0.52 | 0.04 | 118 | F(6,6)= 1.1 | 0.460 |
|  | Age | -0.23 | -1.394 to 0.9282 | 0.6409 |  |  |  |  |  |
|  | GDF15 | -0.01 | -0.04416 to 0.02028 | 0.3995 |  |  |  |  |  |
|  | FGF21 | 0.02 | -0.2925 to 0.3400 | 0.8605 |  |  |  |  |  |
|  | Hsp60 | 21.85 | -86.85 to 130.6 | 0.6402 |  |  |  |  |  |
|  | GLA protein | -11.34 | -97.41 to 74.72 | 0.758 |  |  |  |  |  |
|  | GLA activity | 0.61 | -2.813 to 4.036 | 0.6774 |  |  |  |  |  |
| 2 | Intercept | 104.40 | 58.43 to 150.3 | 0.001 | 0.52 | 0.17 | 102.4 | F(5,7)= 1.5 | 0.299 |
|  | Age | -0.26 | -1.261 to 0.7498 | 0.5665 |  |  |  |  |  |
|  | GDF15 | -0.01 | -0.03636 to 0.01474 | 0.3504 |  |  |  |  |  |
|  | Hsp60 | 15.92 | -50.86 to 82.69 | 0.5906 |  |  |  |  |  |
|  | GLA protein | -7.29 | -67.38 to 52.80 | 0.7825 |  |  |  |  |  |
|  | GLA activity | 0.56 | -2.451 to 3.574 | 0.6728 |  |  |  |  |  |
| 3 | Intercept | 107.50 | 72.56 to 142.4 | 0.0001 | 0.51 | 0.27 | 92.2 | F(4,8)= 2.1 | 0.172 |
|  | Age | -0.31 | -1.140 to 0.5231 | 0.4171 |  |  |  |  |  |
|  | GDF15 | -0.01 | -0.03275 to 0.01267 | 0.3378 |  |  |  |  |  |
|  | Hsp60 | 8.74 | -19.75 to 37.23 | 0.4993 |  |  |  |  |  |
|  | GLA activity | 0.39 | -2.037 to 2.807 | 0.7234 |  |  |  |  |  |
| 4 | Intercept | 109.30 | 78.56 to 140.1 | <0.0001 | 0.50 | 0.34 | 84.9 | F(3,9)= 3.1 | 0.085 |
|  | Age | -0.28 | -1.030 to 0.4762 | 0.4269 |  |  |  |  |  |
|  | GDF15 | -0.01 | -0.03115 to 0.008571 | 0.2306 |  |  |  |  |  |
|  | Hsp60 | 9.88 | -15.84 to 35.60 | 0.4073 |  |  |  |  |  |
| 5 | Intercept | 100.90 | 80.98 to 120.8 | <0.0001 | 0.47 | 0.36 | 80.4 | F(2,10)= 4.4 | 0.043 |
|  | GDF15 | -0.02 | -0.03013 to -0.002708 | 0.0236 |  |  |  |  |  |
|  | Hsp60 | 9.99 | -14.95 to 34.92 | 0.3932 |  |  |  |  |  |
| 6 | Intercept | 107.90 | 92.29 to 123.6 | <0.0001 | 0.55 | 0.52 | 108.9 | F(1,16)= 19.6 | 0.0004 |
|  | GDF15 | -0.02 | -0.03066 to -0.01081 | 0.0004 |  |  |  |  |  |
| 7 | Intercept | 111.00 | 25.59 to 196.5 | 0.0205 | 0.52 | -0.04 | 125.7 | F(6,5)= 0.9 | 0.548 |
|  | Age | -0.24 | -1.575 to 1.086 | 0.6568 |  |  |  |  |  |
|  | GDF15 | -0.01 | -0.05010 to 0.02430 | 0.4135 |  |  |  |  |  |
|  | FGF21 | 0.01 | -0.3546 to 0.3732 | 0.9501 |  |  |  |  |  |
|  | Hsp60 | 9.83 | -116.5 to 136.2 | 0.8494 |  |  |  |  |  |
|  | GLA protein | -1.69 | -93.43 to 90.05 | 0.964 |  |  |  |  |  |
|  | Plasma lyso-Gb3 | -0.19 | -1.499 to 1.120 | 0.7253 |  |  |  |  |  |
| 8 | Intercept | 112.70 | 64.49 to 160.9 | 0.0012 | 0.52 | 0.13 | 103.7 | F(5,6)= 1.3 | 0.367 |
|  | Age | -0.25 | -1.369 to 0.8626 | 0.5987 |  |  |  |  |  |
|  | GDF15 | -0.01 | -0.04058 to 0.01572 | 0.3214 |  |  |  |  |  |
|  | Hsp60 | 7.48 | -68.02 to 82.98 | 0.8166 |  |  |  |  |  |
|  | GLA protein | -0.18 | -61.10 to 60.75 | 0.9946 |  |  |  |  |  |
|  | Plasma lyso-Gb3 | -0.20 | -1.317 to 0.9270 | 0.6854 |  |  |  |  |  |
| 9 | Intercept | 112.80 | 74.25 to 151.3 | 0.0002 | 0.52 | 0.25 | 90.5 | F(4,7)= 1.9 | 0.211 |
|  | Age | -0.25 | -1.138 to 0.6280 | 0.5169 |  |  |  |  |  |
|  | GDF15 | -0.01 | -0.03599 to 0.01118 | 0.2537 |  |  |  |  |  |
|  | Hsp60 | 7.29 | -24.52 to 39.09 | 0.6048 |  |  |  |  |  |
|  | Plasma lyso-Gb3 | -0.20 | -1.167 to 0.7754 | 0.6479 |  |  |  |  |  |

**Table S.** Multiple linear regression analysis for GFR - N215S males.

| Table S. Multiple linear regression analysis for GFR - N215S males |  |  |  |  |  |  |  |  |  |
| --- | --- | --- | --- | --- | --- | --- | --- | --- | --- |
| Model | Independent variables | $\beta$ coefficient | 95% CI | *p*-value | $r^2$ | Adj. $r^2$ | AICc | F | *p*-value |
| 1 | Intercept | 113.40 | -31.17 to 258.0 | 0.078 | 0.75 | 0.25 | n/a | F(4,2)= 1.5 | 0.438 |
|  | GDF-15 | 0.01 | -0.05760 to 0.06753 | 0.765 |  |  |  |  |  |
|  | FGF-21 | -0.10 | -0.6670 to 0.4608 | 0.514 |  |  |  |  |  |
|  | Age | -0.32 | -4.217 to 3.588 | 0.762 |  |  |  |  |  |
|  | Hsp60 | 10.64 | -52.85 to 74.12 | 0.546 |  |  |  |  |  |
| 2 | Intercept | 108.00 | 28.78 to 187.2 | 0.023 | 0.74 | 0.47 | 99.6 | F(3,3)= 2.8 | 0.212 |
|  | FGF-21 | -0.10 | -0.4535 to 0.2471 | 0.418 |  |  |  |  |  |
|  | Age | -0.12 | -1.989 to 1.754 | 0.854 |  |  |  |  |  |
|  | Hsp60 | 11.34 | -27.71 to 50.40 | 0.423 |  |  |  |  |  |
| 3 | Intercept | 103.70 | 72.66 to 134.8 | 0.001 | 0.73 | 0.60 | 57.7 | F(2,4)= 5.4 | 0.072 |
|  | FGF-21 | -0.12 | -0.2766 to 0.03436 | 0.097 |  |  |  |  |  |
|  | Hsp60 | 12.22 | -15.51 to 39.96 | 0.288 |  |  |  |  |  |
| 4 | Intercept | 90.89 | 60.61 to 121.2 | 0.0 | 0.25 | 0.14 | 69.7 | F(1.7)= 2.3 | 0.171 |
|  | FGF-21 | -0.05 | -0.1269 to 0.02746 | 0.1714 |  |  |  |  |  |

**Table T.** Multiple linear regression analysis for GFR - All female cohort.

| Table T. Multiple linear regression analysis for GFR - All female cohort |  |  |  |  |  |  |  |  |  |
| --- | --- | --- | --- | --- | --- | --- | --- | --- | --- |
| Model | Independent variables | $\beta$ coefficient | 95% CI | $p$-value | $r^2$ | Adj. $r^2$ | AICc | F | $p$-value |
| 1 | Intercept | 110.70 | 12.54 to 208.9 | 0.0351 | 0.80 | 0.51 | 135.7 | F(6,4)= 2.7 | 0.176 |
|  | Age | -0.64 | -2.044 to 0.7672 | 0.2758 |  |  |  |  |  |
|  | GDF15 | 0.00 | -0.03264 to 0.02906 | 0.8797 |  |  |  |  |  |
|  | FGF21 | 0.05 | -0.02050 to 0.1159 | 0.1241 |  |  |  |  |  |
|  | Hsp60 | -18.30 | -181.4 to 144.8 | 0.771 |  |  |  |  |  |
|  | GLA protein | 11.14 | -31.72 to 54.00 | 0.5105 |  |  |  |  |  |
|  | GLA activity | -0.12 | -2.391 to 2.144 | 0.8873 |  |  |  |  |  |
| 2 | Intercept | 120.80 | 56.51 to 185.0 | 0.0037 | 0.81 | 0.66 | 95.2 | F(5,6)= 5.2 | 0.035 |
|  | Age | -0.86 | -1.789 to 0.06133 | 0.0624 |  |  |  |  |  |
|  | GDF15 | 0.00 | -0.02135 to 0.02444 | 0.8743 |  |  |  |  |  |
|  | FGF21 | 0.04 | -0.008379 to 0.09600 | 0.0858 |  |  |  |  |  |
|  | Hsp60 | -40.23 | -153.1 to 72.68 | 0.4169 |  |  |  |  |  |
|  | GLA protein | 9.94 | -12.33 to 32.20 | 0.3167 |  |  |  |  |  |
| 3 | Intercept | 120.90 | 63.30 to 178.5 | 0.0016 | 0.81 | 0.70 | 82.1 | F(4,7)= 7.5 | 0.011 |
|  | Age | -0.83 | -1.554 to -0.1123 | 0.0292 |  |  |  |  |  |
|  | FGF21 | 0.04 | -0.002881 to 0.09069 | 0.062 |  |  |  |  |  |
|  | Hsp60 | -41.79 | -140.9 to 57.29 | 0.3518 |  |  |  |  |  |
|  | GLA protein | 10.09 | -9.782 to 29.95 | 0.269 |  |  |  |  |  |
| 4 | Intercept | 104.60 | 62.94 to 146.3 | 0.0004 | 0.78 | 0.70 | 74.9 | F(3,8)= 9.7 | 0.005 |
|  | Age | -0.68 | -1.282 to -0.07367 | 0.0323 |  |  |  |  |  |
|  | FGF21 | 0.05 | 0.001461 to 0.09180 | 0.0445 |  |  |  |  |  |
|  | GLA protein | 9.75 | -9.601 to 29.11 | 0.2787 |  |  |  |  |  |
| 5 | Intercept | 134.30 | 107.3 to 161.3 | <0.0001 | 0.56 | 0.50 | 101.5 | F(2,14)= 8.9 | 0.003 |
|  | Age | -0.97 | -1.464 to -0.4745 | 0.0009 |  |  |  |  |  |
|  | FGF21 | 0.01 | -0.03030 to 0.04786 | 0.6375 |  |  |  |  |  |
| 6 | Intercept | 135.90 | 110.6 to 161.1 | <0.0001 | 0.55 | 0.52 | 98.3 | F(1,15)= 18.4 | 0.0006 |
|  | Age | -0.96 | -1.442 to -0.4853 | 0.0006 |  |  |  |  |  |
| 7 | Intercept | 136.40 | 94.26 to 178.6 | 0.002 | 0.94 | 0.87 | 119.5 | F(4,3)= 12.6 | 0.032 |
|  | Age | -1.06 | -1.905 to -0.2126 | 0.0283 |  |  |  |  |  |
|  | GDF15 | 0.00 | -0.01432 to 0.02259 | 0.5273 |  |  |  |  |  |
|  | FGF21 | 0.03 | -0.01579 to 0.07667 | 0.1271 |  |  |  |  |  |
|  | Plasma lyso-Gb3 | 0.77 | -2.679 to 4.217 | 0.529 |  |  |  |  |  |
| 8 | Intercept | 134.90 | 107.2 to 162.7 | <0.0001 | 0.58 | 0.48 | 105.0 | F(3,13)= 5.9 | 0.009 |
|  | Age | -1.12 | -1.798 to -0.4399 | 0.0035 |  |  |  |  |  |
|  | GDF15 | 0.01 | -0.01505 to 0.02998 | 0.4865 |  |  |  |  |  |
|  | FGF21 | 0.01 | -0.03251 to 0.04789 | 0.6862 |  |  |  |  |  |
| 9 | Intercept | 136.30 | 110.5 to 162.1 | <0.0001 | 0.57 | 0.51 | 101.1 | F(2,14)= 9.3 | 0.003 |
|  | Age | -1.12 | -1.775 to -0.4673 | 0.0025 |  |  |  |  |  |
|  | GDF15 | 0.01 | -0.01379 to 0.02943 | 0.4508 |  |  |  |  |  |

### 3.4 Impact of treatment

No differences became apparent when subjects were analysed based on treatment type (fig. 15). Nonetheless, correlation analyses between age at treatment commencement and serum mtUPR markers showed that those males who started treatment at an older age had elevated levels of serum mitokines (FGF-21: r2=0.43 p= 0.04 and GDF-15: r2= 0.58 p= 0.006; fig. 16). Females showed the same association only with GDF-15 (r2=0.70 p= 0.007, fig. 16). When males were stratified based on treatment type, correlation analyses showed that while subjects who started PCT at an older age had higher levels of both mitokines (FGF-21: r2=0.65 p= 0.002, and GDF-15: r2= 0.72, p= 0.008, fig. 16), those who started ERT older had higher levels of GDF-15 only (r2=0.93 p= 0.003, fig. 16).

**Fig. 15.**
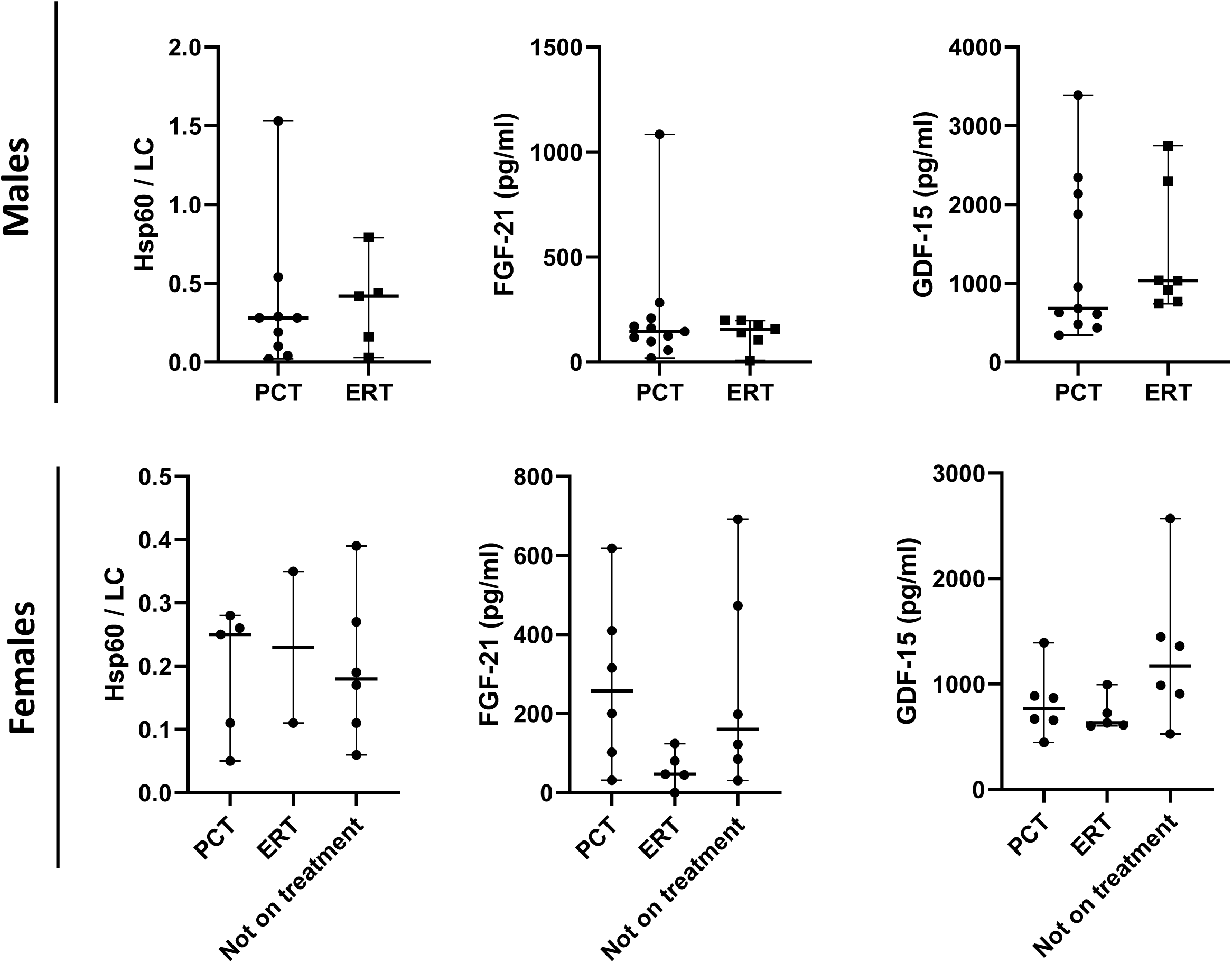
Impact of treatment on mtUPR markers: levels of Hsp60, FGF-21, and GDF-15 were compared in subjects with FD.

**Fig. 16.**
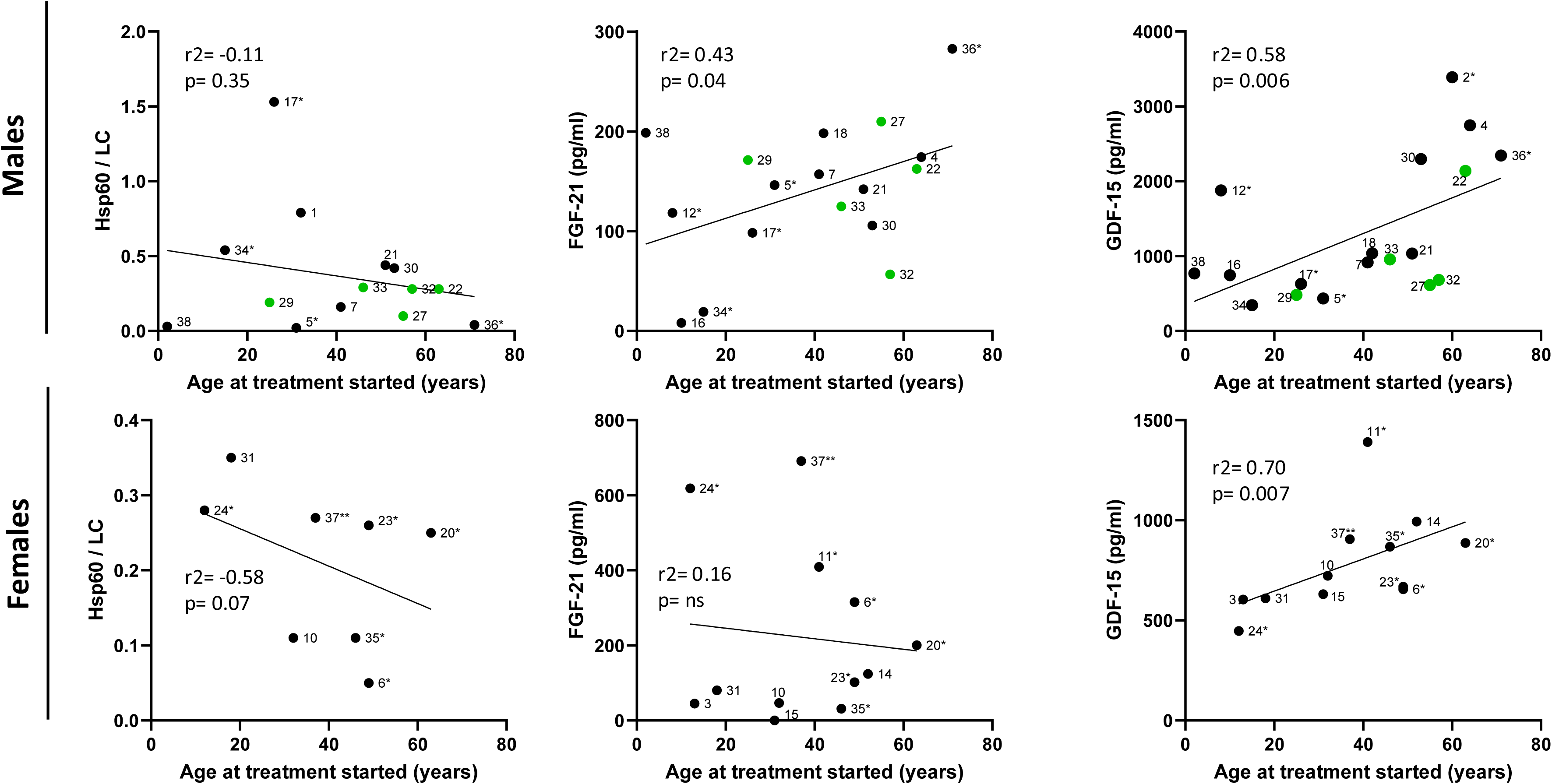
Impact of treatment on mtUPR markers: correlation analyses between markers of mtUPR and age at treatment commencement. Patients who started on enzyme replacement therapy (ERT) are in black whilst those who started on pharmacological chaperon therapy (PCT) are in green. Subjects with * started on ERT and switched to PCT.

## 4.0 Discussion

FD is clinically heterogeneous and currently age is the most significant correlate of phenotypic diversity. While mitochondrial dysfunction is associated with normal aging and has been reported in FD, it has not been systematically studied. Considering this knowledge gap, this study focuses on the mtUPR and its potential impact on clinical expression in FD. It has been suggested that this response has a cytoprotective role, and higher mtUPR intracellular marker levels are reported to associate with less cell death, lower inflammation markers, and less symptom severity in individuals with osteoarthritis and with myocardial chronic pressure overload (Zhou et al. 2022; Smyrnias et al. 2019).

In this study, higher hsp60 intracellular protein levels corresponded with lower LVMI magnitudes in males (fig. 5 and table P, model 4). While this effect seemed to impact on global severity scores and kidney function in N215S males only, their non-N215S counterparts showed the opposite, with higher levels of hsp60 relating to higher MSSI, AASS, and lower GFR (figs. 4, 5, supplementary fig. 3 and 4). Given that the N215S genotype is associated with a cardiac phenotype in FD (table 1)(Lavalle et al. 2018; Germain et al. 2018), these results could suggest a cytoprotective role for hsp60 in male cardiomyocytes, reflected in lower overall severity scores with higher levels of hsp60 for this patient group. Conversely, for the non-N215S group, who presented more classical cardinal features and higher overall severity scores (table 1), higher levels of hsp60 may result in lower LVMI magnitudes without impacting on global severity scores (supplementary fig. 3). Accordingly, multivariate models for MSSI suggested an indirect relation between this score and hsp60 in both male groups (tables J and K). These findings could support a cytoprotective role for the mtUPR in cardiomyocytes in individuals with FD.

Conversely, higher hsp60 intracellular protein levels were associated with higher LVMI magnitudes in females (fig. 5 and table Q, model 1). No association was found with the MSSI (supplementary fig. 3). Instead, females hsp60 levels showed a significant association with the AASS, suggesting higher levels of this heat shock protein relate to higher clinical severity adjusted for age (fig. 4). Additionally, females showed a significant negative correlation between hsp60 levels and age (supplementary fig. 6), in contrast to HC who showed a positive trend between these parameters (fig. 3A). While no difference was found in hsp60 amounts between sex groups, these strong associations were absent in males regardless of their genotype (fig. 4 and supplementary fig. 6). These observations are opposed to the purported cytoprotective role hypothesised for the mtUPR and to the findings reported for male subjects.

The analysis based on the predictive value of intracellular levels of Hsp60 suggested a difference between biological sex groups. For males, total GLA protein was the only variable that consistently achieved statistical significance regardless of genotype in all the models produced (table A and B). However, for females this was age at sampling (table C). Therefore, to assess if the above discussed differences in hsp60 relate to the expression of the mutant GLA gene, the relation between hsp60 and the amount of GLA endo-H 50 kDa sensitive ER form was studied. A strong negative correlation was only found for females, suggesting higher levels of endo-H sensitive 50kDA ER form relate to lower hsp60 levels (r2= -0.79 p= 0.003, supplemental fig. 7). Work from the Germain group has shown the mtUPR response can be influenced by sex-specific factors (Riar et al. 2017). The group used a G93A-SOD1 transgenic mouse as a model of Amyotrophic Lateral Sclerosis (ALS) to study the relation between misfolded SOD1, the mtUPR, and sex differences in disease phenotype as females with ALS have a longer lifespan than males. Sex differences were observed in the mtUPR axes CHOP and ERα. In female transgenic mice, markers of the CHOP axis (i.e., CHOP and hsp60) were elevated at symptomatic stages, while in males these markers remained similar to or below control levels. In terms of the ERα axis, males exhibited an increase in several effector markers during pre-symptomatic stages only. Among these markers, females only showed an increase in OMI at the symptomatic stage. Additionally, male transgenic mice demonstrated decreased proteasomal activity and an increase in ubiquitinated proteins while females exhibited an increase in proteasomal activity and no increase in ubiquitinated proteins. Of note, the group confirmed that the results observed were not due to overexpression of SOD1, as they examined the levels of OMI and ubiquitinated proteins in mice overexp ressing WT SOD1. While OMI levels were lower in WT female mice, protein ubiquitination was lower in WT male mice, confirming that the effects of OMI and the proteasome are caused by mutant and not by WT SOD1 expression (Riar et al. 2017). The authors concluded that sex differences may influence the activation of the mtUPR and suggested that oestrogen and its receptors could potentially act as drivers through the mtUPR-ERα axis (Riar et al. 2017). Therefore, the contrasting results seen between males and females LVMI relations with hsp60 could be in line with the results from the Germain group. Additionally, the observation that higher hsp60 levels correspond to younger female patients, could also support the role proposed for oestrogens in driving these differences as the levels of these hormone are known to decrease with age.

FGF-21 and GDF-15 are reported to increase with age in healthy individuals, however, in this study, correlations between serum levels of these and age were stronger for Fabry patients (fig. 7). Given the continuum described between FD clinical progression and aging (Hughes et al. 2010), these findings could be in line with the fact that age-related disorders share the same basic molecular mechanisms with aging, the ‘seven pillars’ (Kennedy et al. 2014). Consistently, higher levels of both mitokines were associated with higher MSSI in males, with the N215S group showing stronger associations (fig. 9 and 10). While for both sex groups FGF-21 and GDF-15 were associated with LVMI and GFR, respectively, FGF-21 was associated with kidney function only in N215S males (figs. 13 and 14). Given that this group was the only one in which LVMI was strongly associated with age (supplemental fig. 2 and 3), taken together, these observations could suggest that FGF-21 serum levels might be reflecting the underlying mechanisms driving accelerated cardiac aging in male subjects with FD. Accordingly, hsp60 amounts resulted predictive of FGF-21 in males (table D, model 4) and levels of this mitokine resulted predictive for MSSI in the N215S male group only (table K, model 4). Overall, these could maybe suggest that hsp60 and FGF-21 might be involved in the pathways driving heart aging and some of the heart pathology in males with FD.

Nevertheless, MSSI modelling for the whole male cohort indicated as statistically relevant predictors GDF-15, total GLA protein, and hsp60 (table J, model 4). While predictor analyses suggested a link between the latter two and FGF-21 serum levels in males (tables D, model 4), for GDF-15 the statistically significant predictor was age (table G, model 4). GDF-15 has been acknowledged as a key protein linked to aging, multimorbidity, and frailty (Conte et al. 2020; Tanaka et al. 2020). Additionally, this mitokine has been associated with CKD progression (Nair et al. 2017) and more recently it was shown to enhance the expression of the KLB in this organ. While a compensatory increase in experimental CKD was noted, exogenous administration of GDF-15 was nephroprotective (Valiño-Rivas et al. 2022). In males, this mitokine was the only statistically significant predictor of GFR (table R, model 5), therefore the results could hint that GDF-15 might be implicated in both, kidney aging and kidney pathology in male subjects with FD. In females, MSSI and AASS modelling both suggested as predictors the level of plasma lyso-Gb3 (table O, model 10 and table L, model 10). However, age resulted predictive for GFR (table T, model 4) and for GDF-15 in this group (table I, model 5). Unlike for heart pathology, these results would be in line with results exhibited by males and support GDF-15 role in kidney aging and in Fabry kidney disease.

Regarding the influence of therapeutic interventions, the associations observed between age at treatment commencement and mitokines serum levels (fig. 16) could reflect FD aging phenotype and the effect of early treatment initiation. Individuals who started treatment at an earlier age could exhibit lower levels mitokines due to the treatment effect on Gb3 (fig. 16). Additionally, the distinct associations observed between PCT and ERT in relation to FGF-21 levels and age of treatment initiation, might also suggest an effect of treatment type on serum levels of this mitokine in males (fig. 17). Indeed, GLA protein proved to be a predictor for hsp60 (table A, model 5), which appeared to be predictive of FGF-21 serum levels (table D, model 4) and LVMI (table P, model 4) in males. These results could potentially suggest that treatment approaches that do not alleviate the accumulation of misfolded GLA protein may not be sufficient to tackle disease progression in males with missense GLA variants. As PCT successfully corrects the trafficking defects of amenable GLA missense variants (Yam, Zuber, and Roth 2005), new therapeutic strategies focusing on enhancing protein degradation are under investigation for FD (Seemann et al. 2020; Mohamed et al. 2017).

**Fig. 17.**
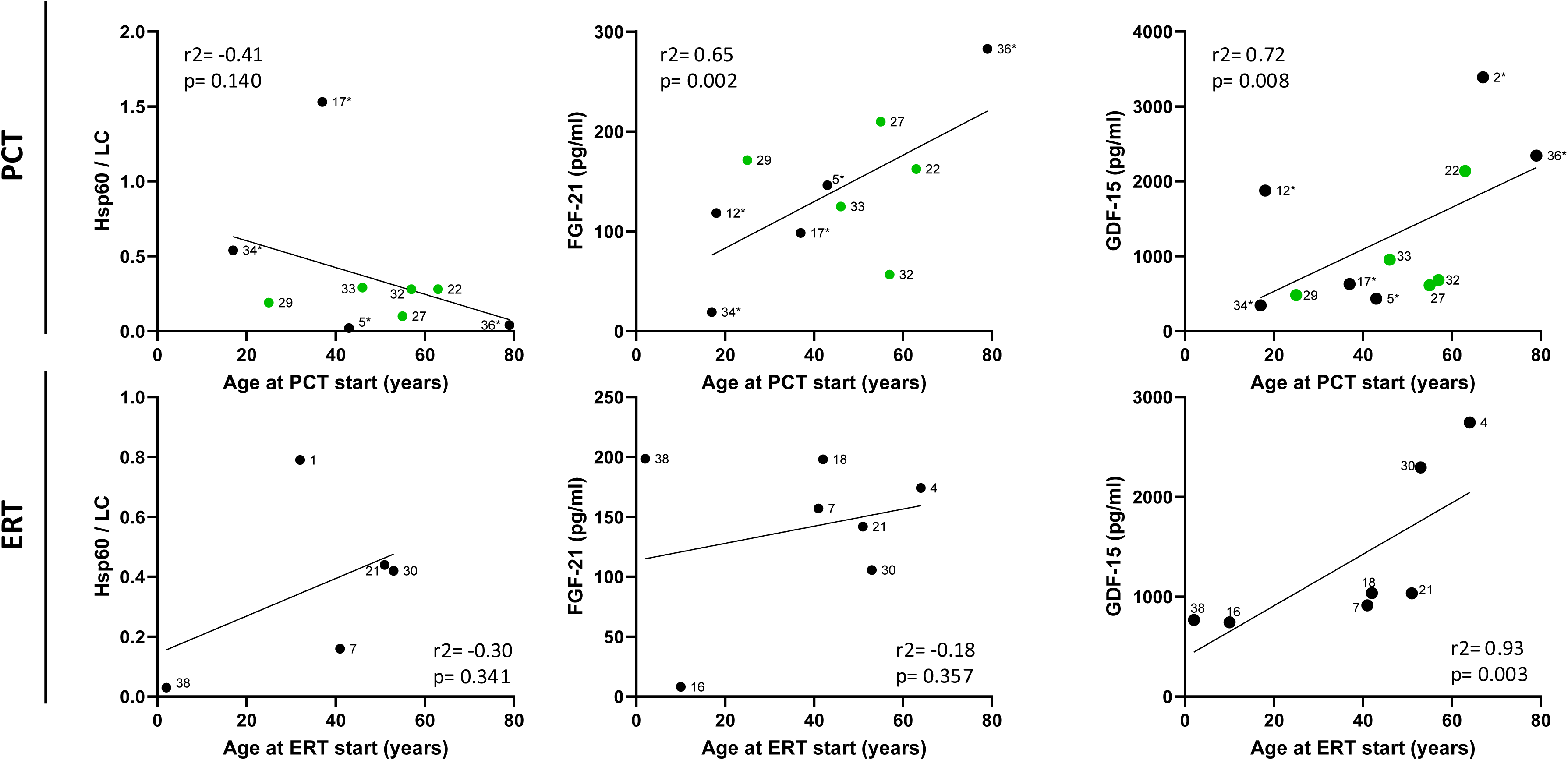
Impact of treatment type on mtUPR markers in male subjects with Fabry disease: correlation analyses between markers of mtUPR and age at treatment commencement. (A) Patients who are currently on pharmacological chaperon therapy (PCT) and (B) patients <u>who have only been on</u> enzyme replacement therapy (ERT). Patients who started on ERT are in black whilst those who started on PCT are in green. Subjects with * started on ERT and switched to PCT.

Activation of the mtUPR leads to an amelioration of pathophysiological alterations and a deceleration in the progression of pathology in numerous disease models including non-alcoholic fatty liver disease (Weng et al. 2023), Parkinson’s (Liu et al. 2020), and Huntington’s (Naia et al. 2021; Cilleros-Holgado et al. 2023). In the heart, recent in vivo and in vitro experiments showed that Carvedilol, a β-blocker, was able to downregulate microRNA-1 expression (a known pro-apoptotic miRNA and increased in T2D cardiomyocytes (Shan et al. 2010)), resulting in an increase expression of hsp60 and an improvement in cardiomyocytes viability as well as in cardiac function within infarcted rats (Hu et al. 2019). In the context of T2D, where intracellular reduced levels of hsp60 seem to contribute to the downregulation of the insulin-like growth factor 1 receptor in the myocardium (Chen et al. 2005) and potentially also in the hypothalamus, treatment with leptin was able to restored hsp60 levels and improve mitochondrial function in mice models of the disease (Kleinridders et al. 2013).

However, excessive activation may be detrimental, exacerbating both the pathology and symptoms across various diseases, such as cancer (Tang et al. 2022). It has been postulated that when heat shock proteins are overexpressed in stress response, they are relocated to the plasma membrane, acting as auto-antigens recognized by various immune cells and antibodies (Mayr et al. 1999), like a danger signal in the immune response, potentially causing inflammatory diseases (Zininga, Ramatsui, and Shonhai 2018). Therefore, its potential role as a candidate for vaccines or as an adjuvant is being studied for atherosclerosis, a chronic inflammatory condition (Hu et al. 2018; Li et al. 2012; van Puijvelde et al. 2007; Krishnan-Sivadoss et al. 2021) and maybe relevant for Fabry’s inflammatory cardiomyopathy (Augusto et al. 2019). Similarly, chronic mitokine signalling can lead to continuous mitochondrial stress response activation in different tissues, as seen in certain types of cancers (Franz et al. 2019) or heart failure (Tsai et al. 2018) (and in normal aging) (Conte et al. 2019). This may impair inter-organ mitochondrial stress signalling, for which pharmacological interventions to mitigate mitokine signalling may have potential benefits in various diseases (Burtscher et al. 2023), including Fabry.

Biomarkers are defined as indicators of normal biological and pathological processes (’FDA-NIH Biomarker Working Group. BEST (Biomarkers, EndpointS, and other Tools) Resource’ 2016). While diagnostic biomarkers are disease specific, allowing to identify individuals with a specific condition, in the context of age-related disorders, biomarkers of biological age have been proposed as prognostic biomarkers (Franceschi et al. 2018). As these provide information about the status of molecular mechanisms of age-related decline (the ‘seven pillars’), combined with diagnostic biomarkers, these could identify disease-specific aging trajectories, particularly important in conditions with subclinical incubation periods where treatment initiation is recommended before the onset of irreversible complications (Franceschi et al. 2018), like FD (Hughes 2017). In T2D, serum GDF-15 significantly improved the reliability of HbA1c in the assessment of glycemic control and in the diagnosis of complications (Conte et al. 2021). Given that the results presented in this work are cross-sectional and correspond to a single time point, the association found between mtUPR markers and clinical outcomes should be validated in bigger cohorts.

## 5.0 Limitations

This is a cross-sectional study and the laboratory results presented correspond to a single time point sample. Clinical data for all patient was assessed retrospectively and only a subset of patients had lyso-Gb3 levels and LVMI data available for analysis. Given the rarity of FD, the sample size was relatively small and the mtUPR predictor analyses included only a minor proportion of patients. In addition, the narrow number of patients restricted the number of independent variables included in the models, excluding other potentially relevant factors. Thus, the predictive value of these needs to be validated in bigger cohorts. Besides, this study did not quantify the impact of the environment, of other comorbidities, or of concomitant medication which highly likely influence the findings. Therefore, a more robust, appropriately designed study is needed to allow for the adjustment for any potential difference between groups in the models to help estimate causal effects from the observed data. Additionally, while hsp60 was measured by quantitative Western blotting, the linear dynamic range was assessed only for the anti-Na+/K+ ATPase antibody (LC). This omission introduces an additional potential source of variation in the quantification of hsp60, impacting the mtUPR predictor analyses.

## 6.0 Conclusion

This is the first study that investigates markers of both intracellular and systemic mtUPR activation in individuals with FD. While findings suggest these are associated with evidence of heart and kidney disease, their potential association with FD progression and therapy need to be further investigated. These results are in line with the proposed compensatory role for the mtUPR in proteostasis (Cilleros-Holgado et al. 2023) and suggest that variation in the activation levels relate to the clinical severity in individuals with FD.

