## Supplemental figures for "Mitochondrial stress in Fabry disease"

### Slide 1
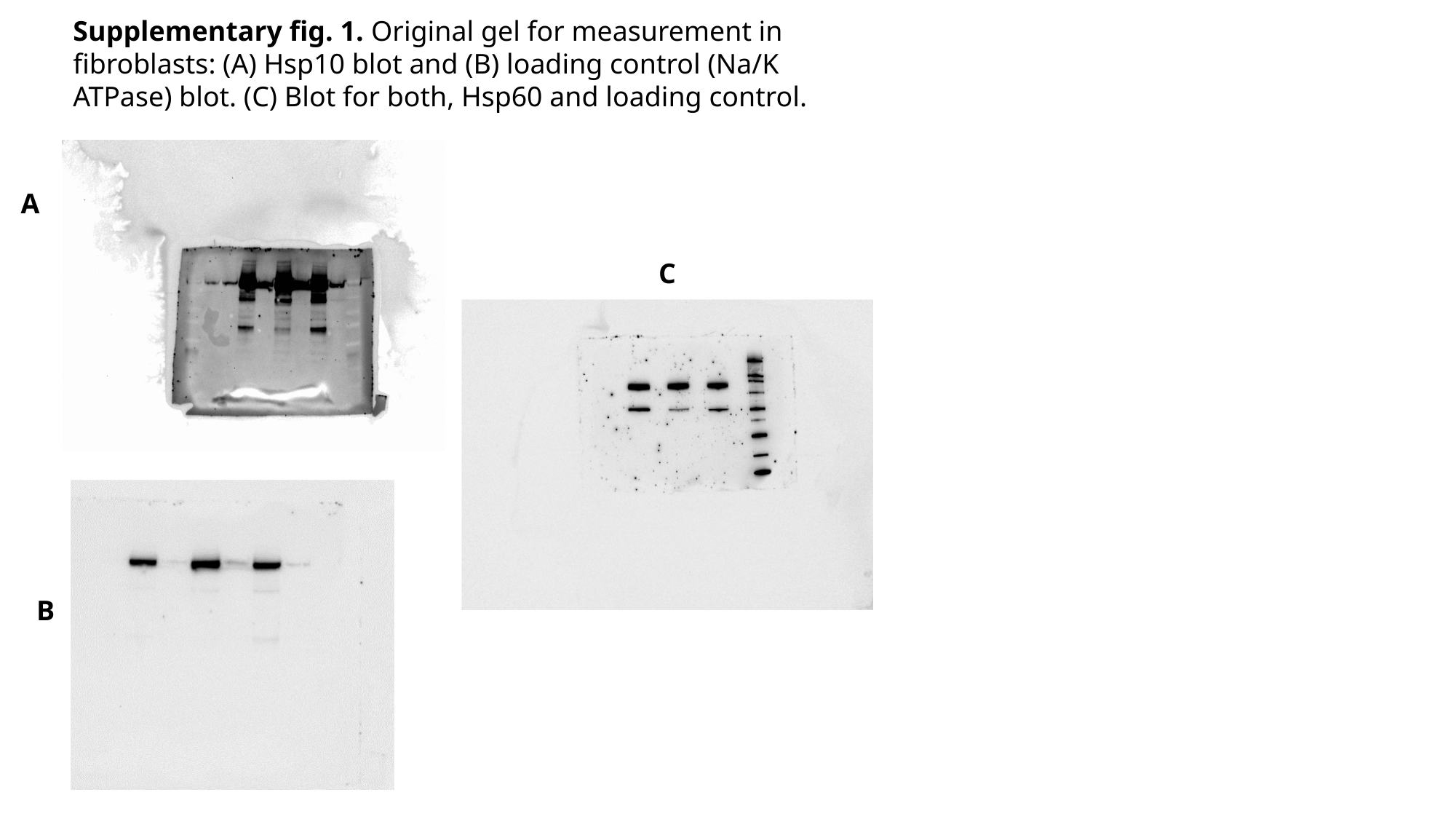

Supplementary fig. 1. Original gel for measurement in fibroblasts: (A) Hsp10 blot and (B) loading control (Na/K ATPase) blot. (C) Blot for both, Hsp60 and loading control.
A
C
B

### Slide 2
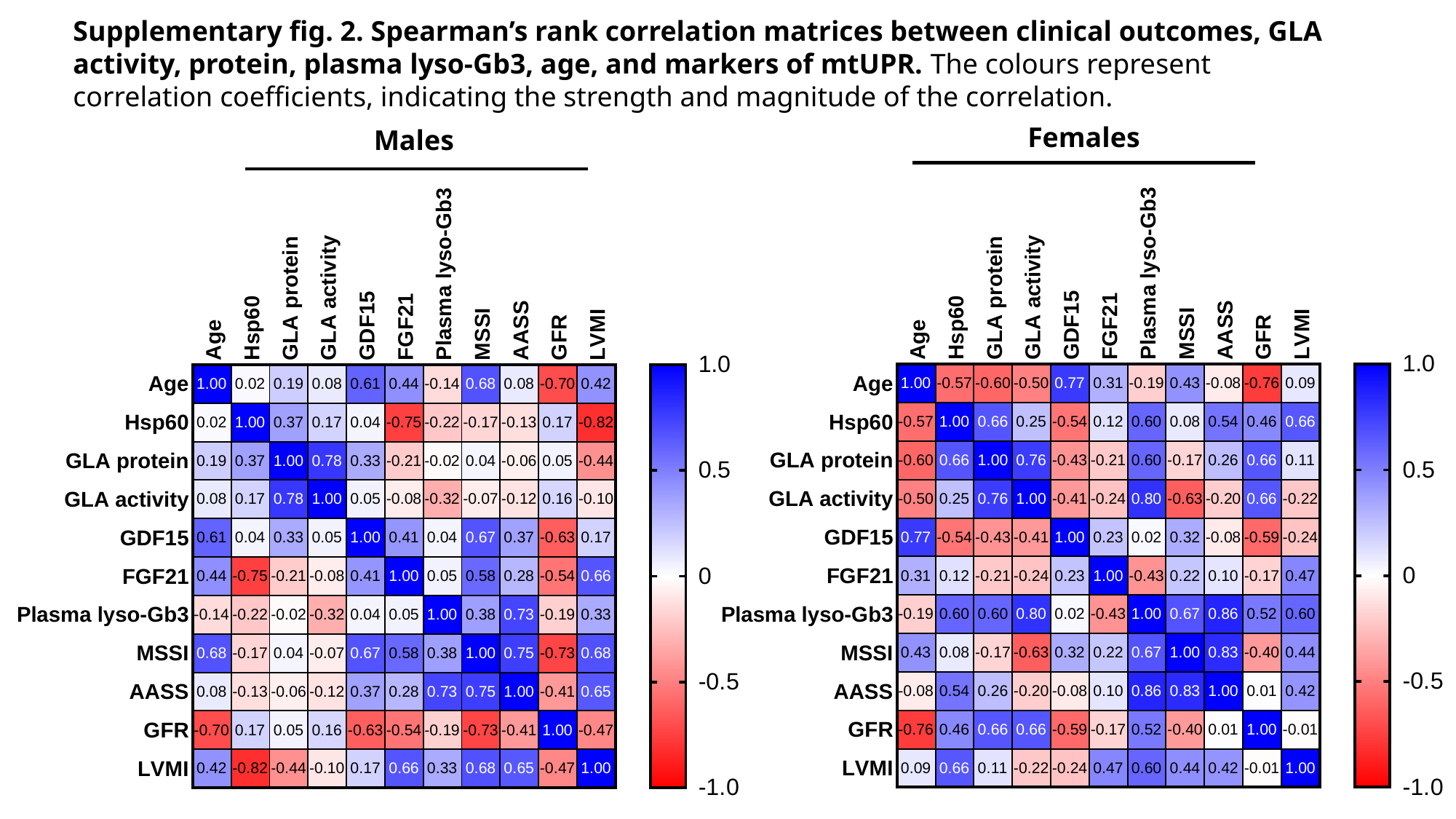

Supplementary fig. 2. Spearman’s rank correlation matrices between clinical outcomes, GLA activity, protein, plasma lyso-Gb3, age, and markers of mtUPR. The colours represent correlation coefficients, indicating the strength and magnitude of the correlation.
Females
Males

### Slide 3
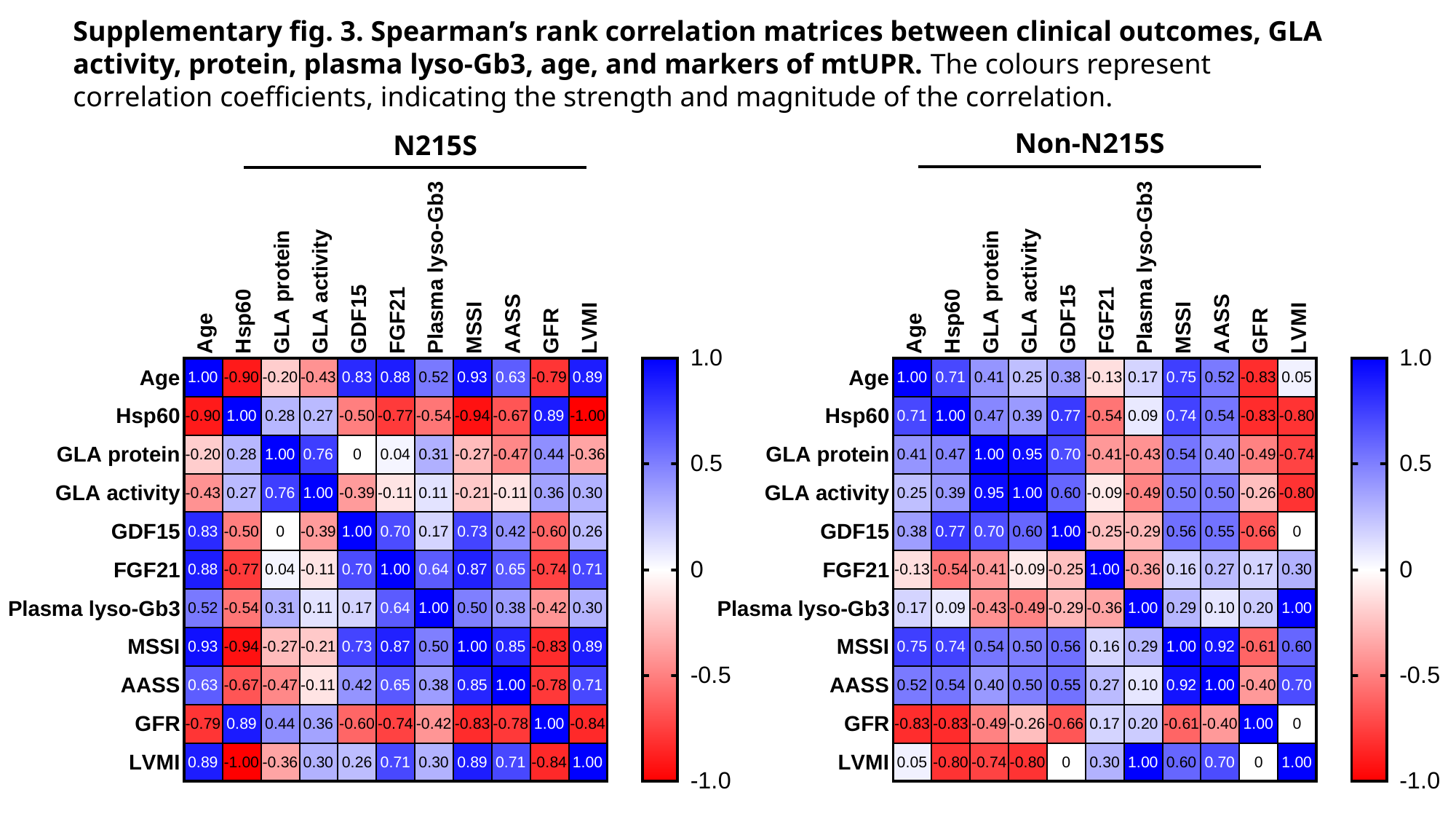

Supplementary fig. 3. Spearman’s rank correlation matrices between clinical outcomes, GLA activity, protein, plasma lyso-Gb3, age, and markers of mtUPR. The colours represent correlation coefficients, indicating the strength and magnitude of the correlation.
Non-N215S
N215S

### Slide 4
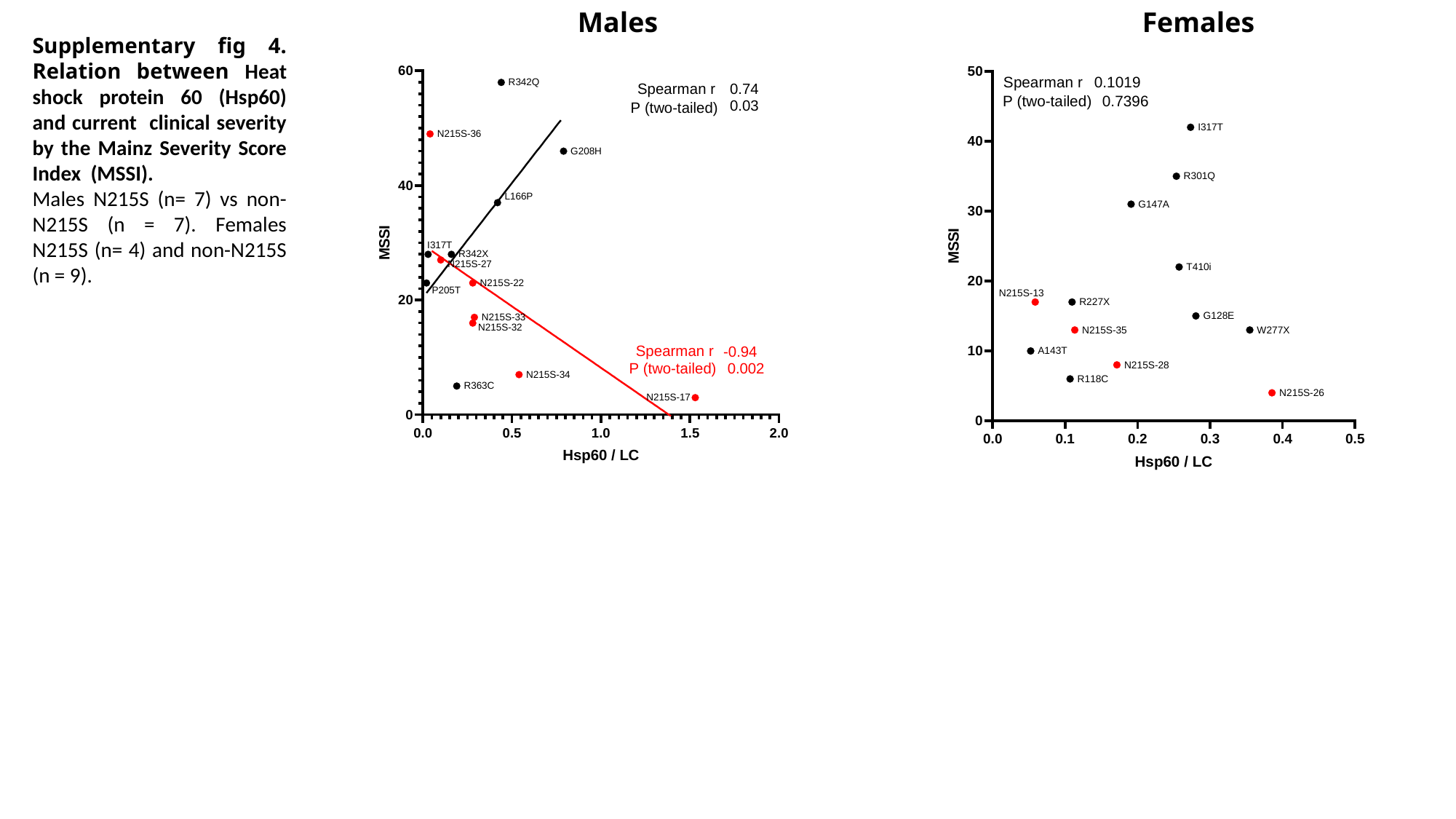

Males
Females
Supplementary fig 4. Relation between Heat shock protein 60 (Hsp60) and current clinical severity by the Mainz Severity Score Index (MSSI).
Males N215S (n= 7) vs non-N215S (n = 7). Females N215S (n= 4) and non-N215S (n = 9).

### Slide 5
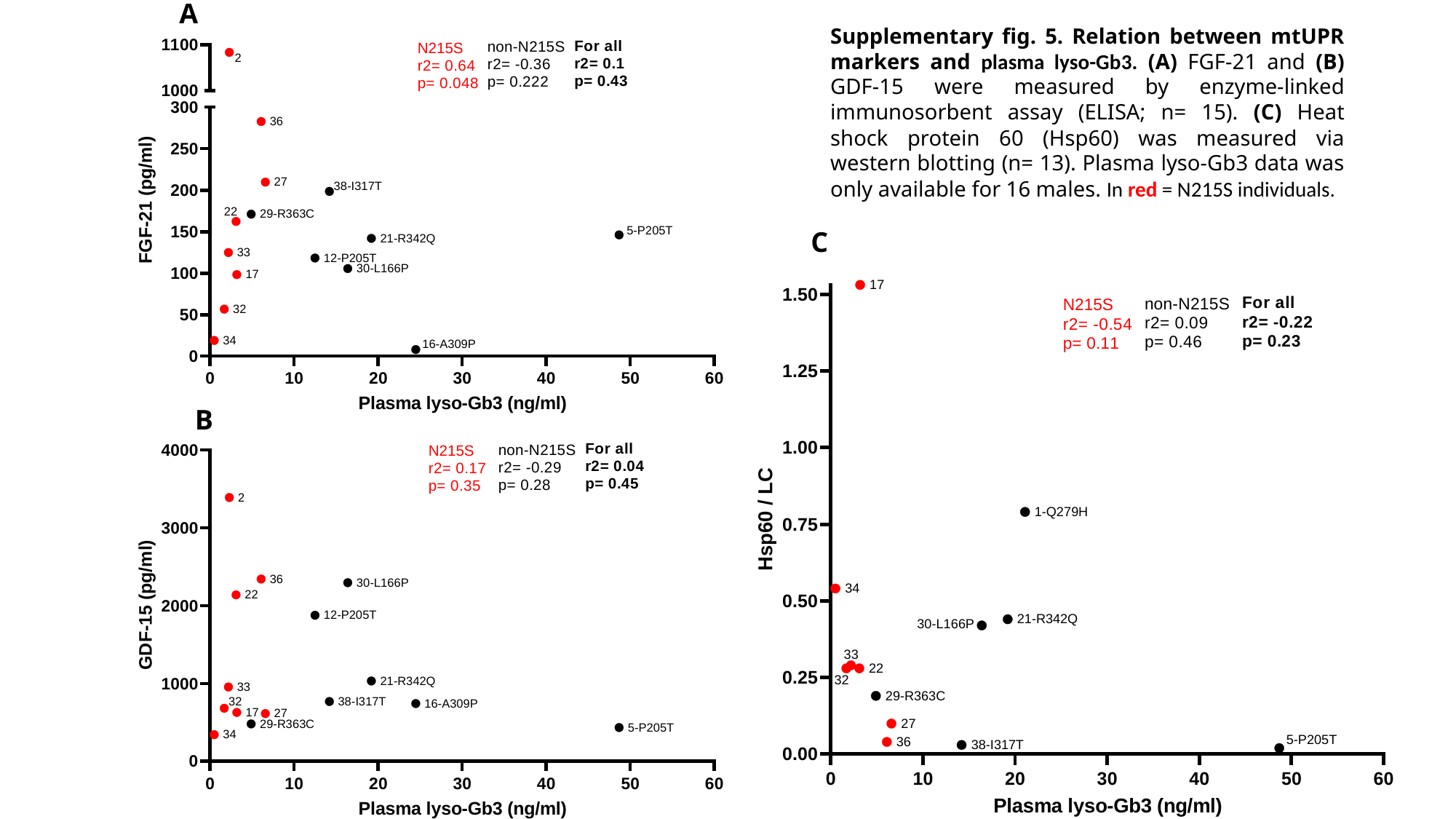

A
Supplementary fig. 5. Relation between mtUPR markers and plasma lyso-Gb3. (A) FGF-21 and (B) GDF-15 were measured by enzyme-linked immunosorbent assay (ELISA; n= 15). (C) Heat shock protein 60 (Hsp60) was measured via western blotting (n= 13). Plasma lyso-Gb3 data was only available for 16 males. In red = N215S individuals.
C
B

### Slide 6
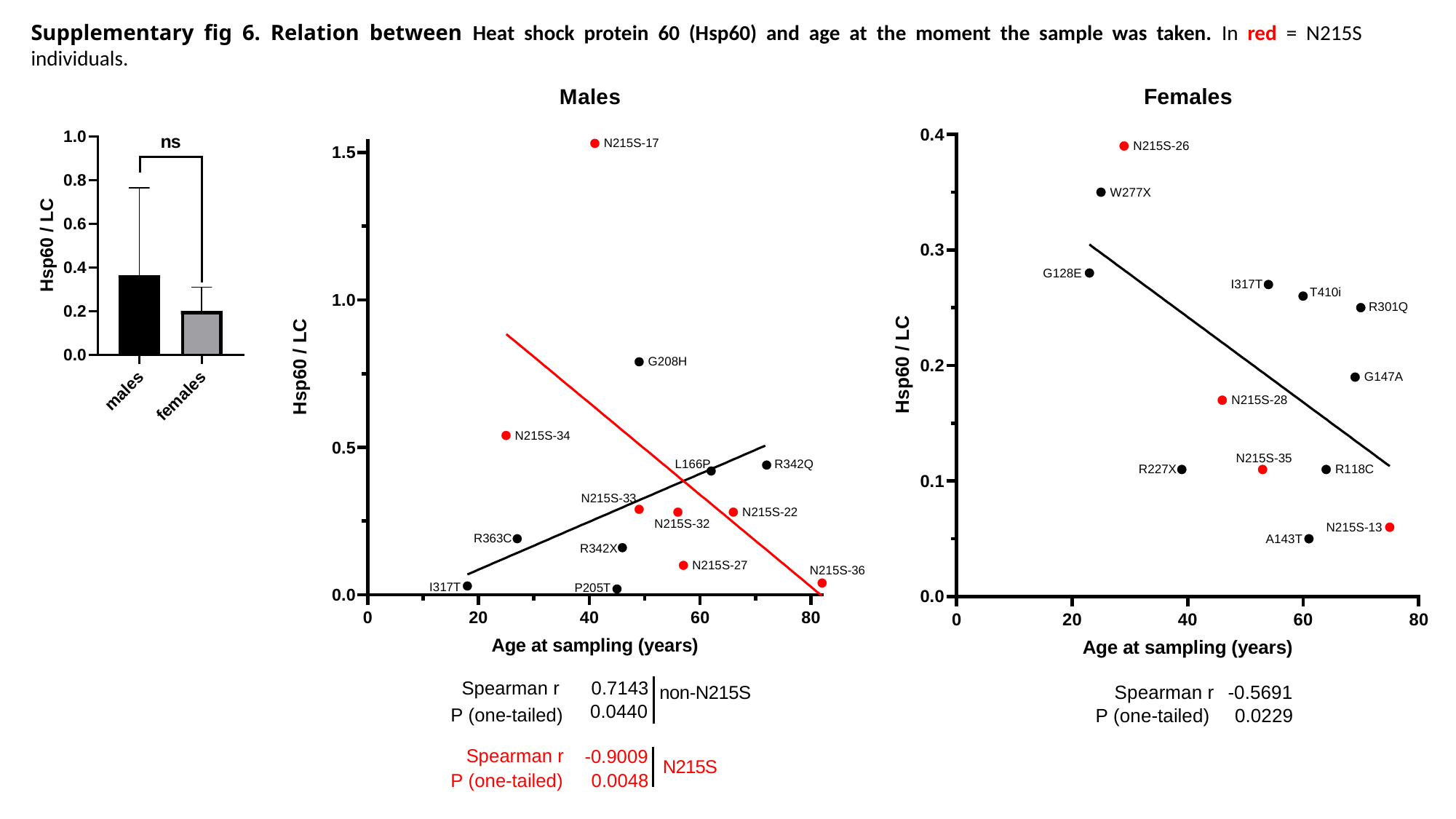

Supplementary fig 6. Relation between Heat shock protein 60 (Hsp60) and age at the moment the sample was taken. In red = N215S individuals.

### Slide 7
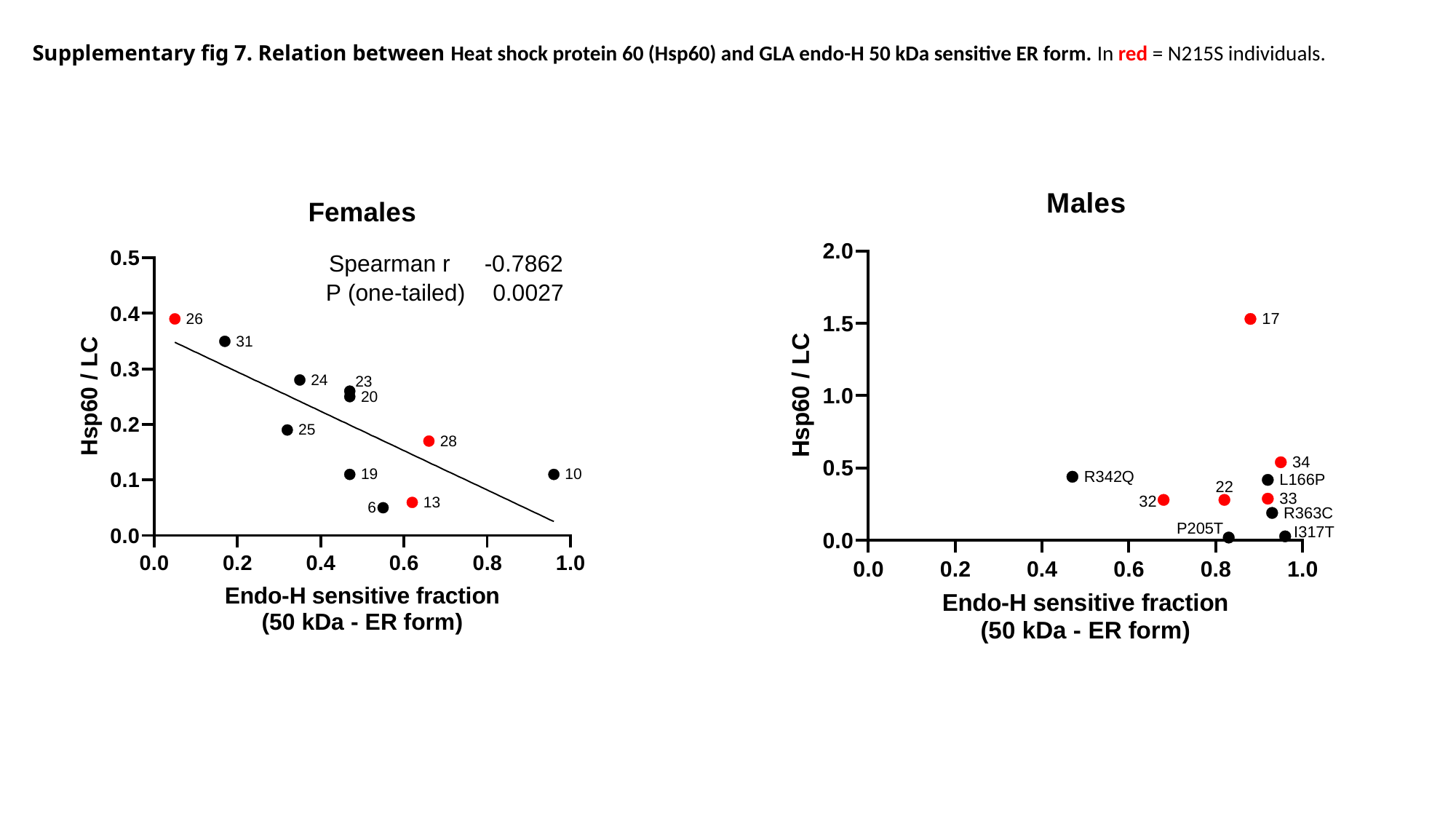

Supplementary fig 7. Relation between Heat shock protein 60 (Hsp60) and GLA endo-H 50 kDa sensitive ER form. In red = N215S individuals.

### Slide 8
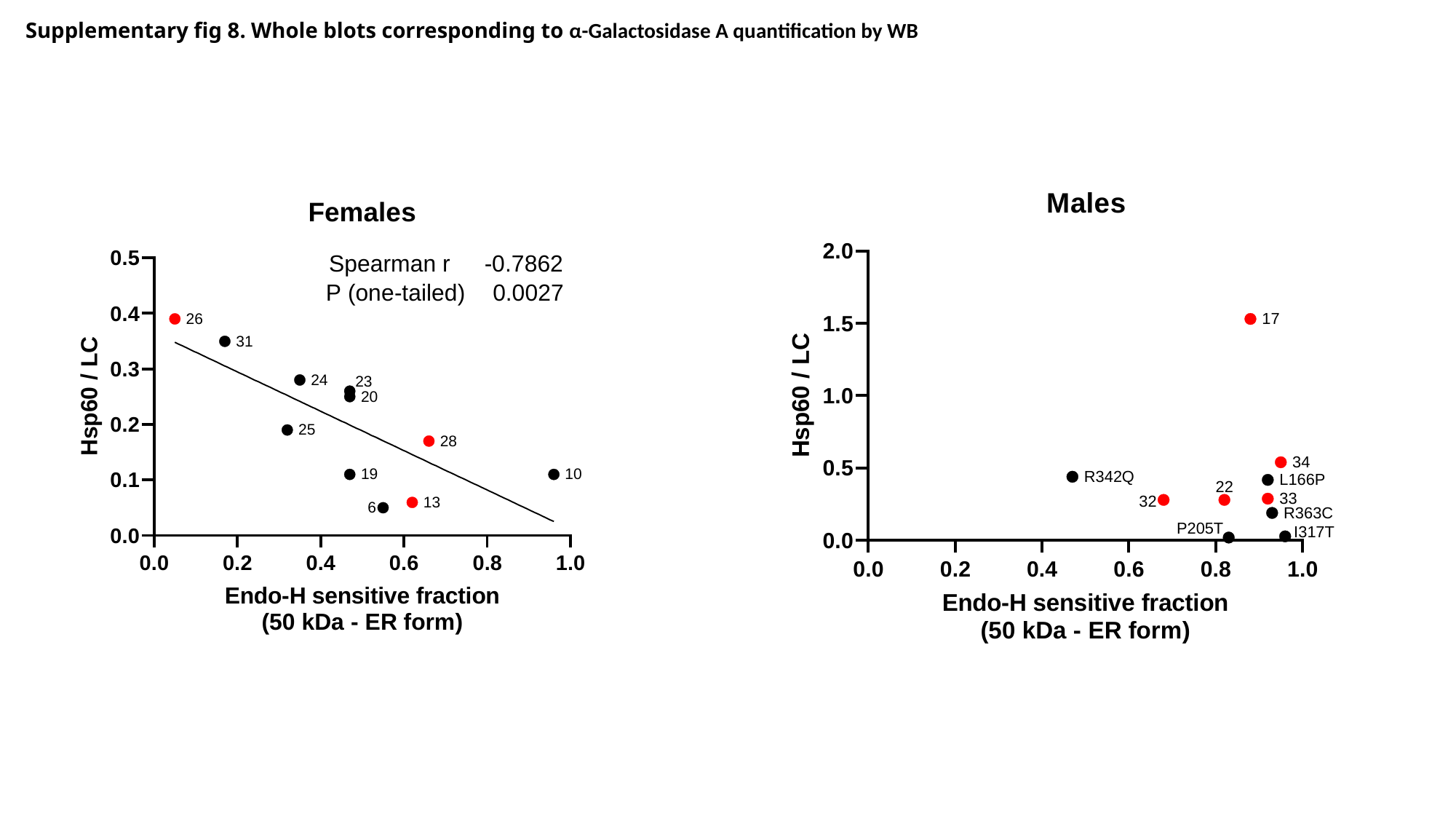

Supplementary fig 8. Whole blots corresponding to α-Galactosidase A quantification by WB

### Slide 9
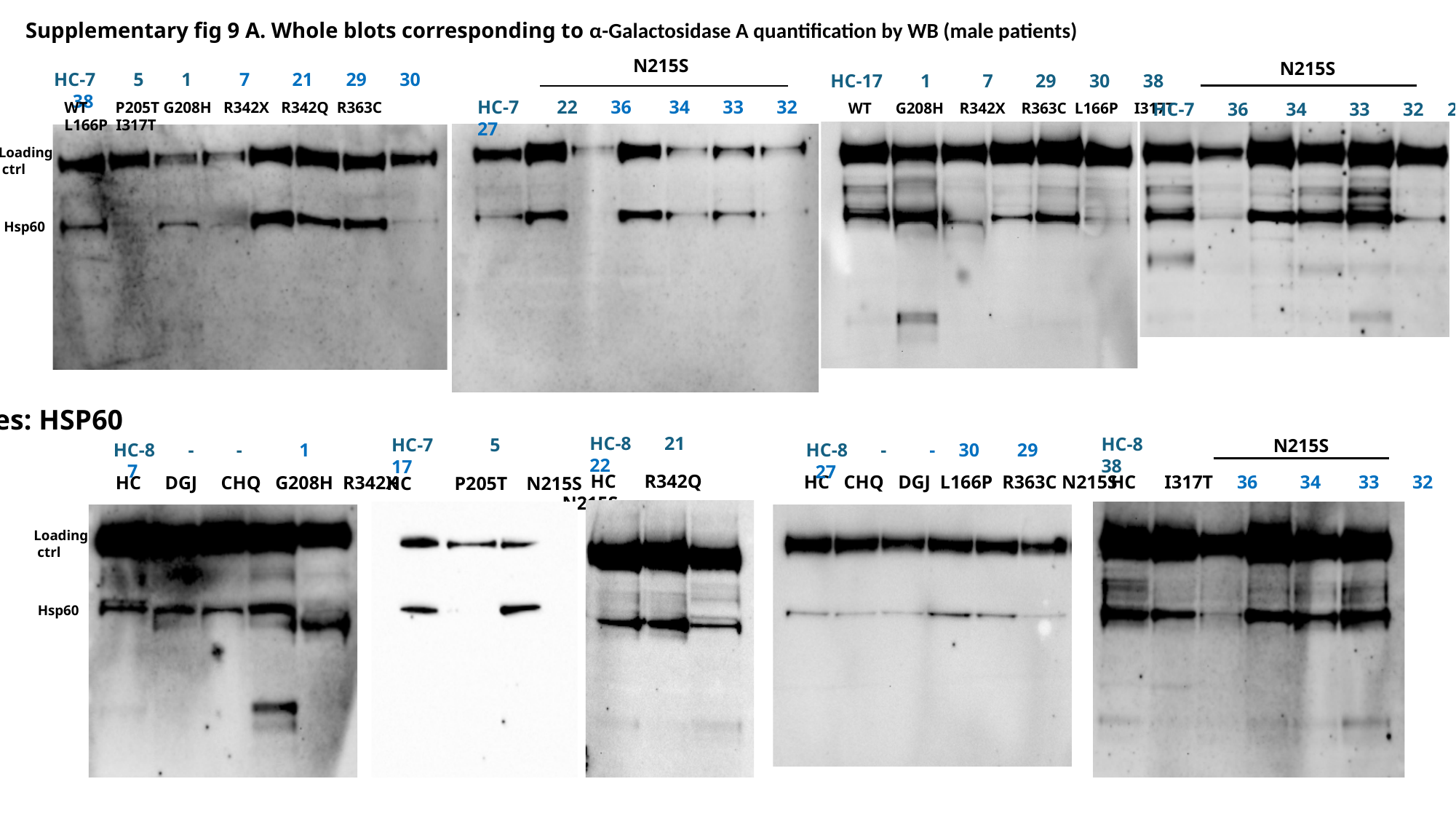

Supplementary fig 9 A. Whole blots corresponding to α-Galactosidase A quantification by WB (male patients)
N215S
HC-7 22 36 34 33 32 27
N215S
HC-7 5 1 7 21 29 30 38
WT P205T G208H R342X R342Q R363C L166P I317T
HC-17 1 7 29 30 38
WT G208H R342X R363C L166P I317T
HC-7 36 34 33 32 27
Loading
 ctrl
Hsp60
Males: HSP60
HC-8 21 22
HC-8 38
HC-7 5 17
N215S
HC I317T 36 34 33 32
HC-8 - - 1 7
HC-8 - - 30 29 27
 HC P205T N215S
 HC R342Q N215S
 HC CHQ DGJ L166P R363C N215S
 HC DGJ CHQ G208H R342X
Loading
 ctrl
Hsp60

### Slide 10
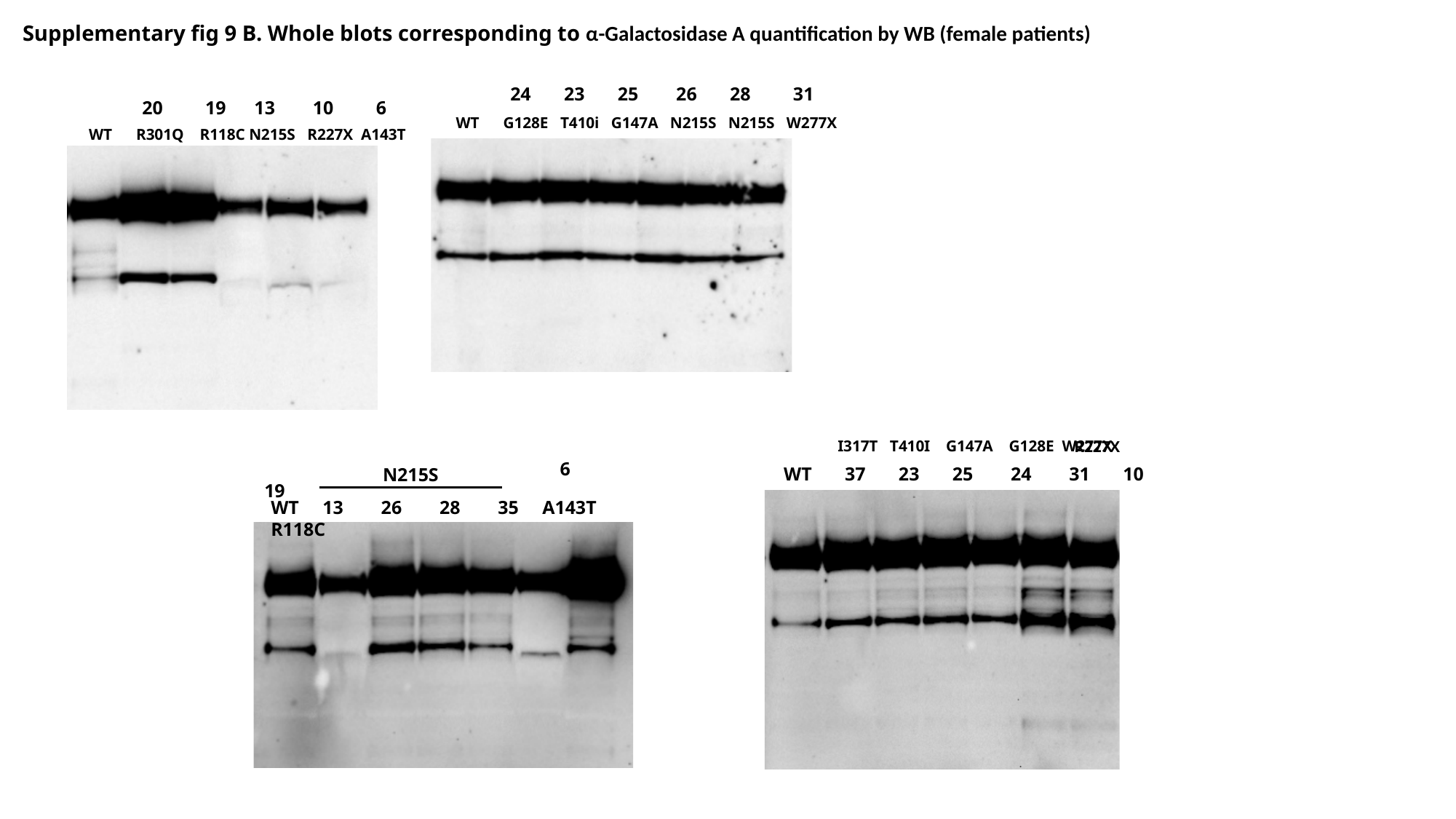

Supplementary fig 9 B. Whole blots corresponding to α-Galactosidase A quantification by WB (female patients)
 24 23 25 26 28 31
 WT G128E T410i G147A N215S N215S W277X
 20 19 13 10 6
 WT R301Q R118C N215S R227X A143T
 I317T T410I G147A G128E W277X
R227X
 WT 13 26 28 35 6 19
N215S
WT 13 26 28 35 A143T R118C
WT 37 23 25 24 31 10
